# Dental calculus as a record of Pleistocene reindeer oral, digestive and dietary flora

**DOI:** 10.1101/2025.08.02.668267

**Authors:** Fabian L. Kellner, Jaelle C. Brealey, Nicola Vogel, Vanessa C. Bieker, Sarah L. F. Martin, Martin Seiler, Bente Philippsen, Vebjørn Veiberg, Mikkel Winther Pedersen, Katerina Guschanski, Michael D. Martin

## Abstract

Dental calculus offers an extraordinary window into the past, preserving a rich archive of genetic material that can unlock new insights into ancient ecosystems. By capturing traces of oral microbiomes, dietary components, and even gut microbes regurgitated by ruminants, dental calculus provides an extraordinary window into the past to reconstruct ecological interactions and past environmental conditions. Here, we harness the power of ancient metagenomics to explore the oral microbiome, digestive microbiomes and diet of Pleistocene reindeer (*Rangifer tarandus*) from archaeological sites in France, a region that once served as a glacial refugium before reindeer disappeared from the area. We used shotgun metagenomic sequencing to assemble microbial genomes (MAGs) and classify microbial and dietary reads from dental calculus of 19 ancient reindeer (ca. 12,000 - 23,000 years BP) and 27 modern and historical (1861 - 1958 CE) Scandinavian reindeer. Notably, six bacterial taxa associated with the rumen microbiome were consistently detected across both ancient and modern samples, offering a rare glimpse into the continuity of digestive adaptations over thousands of years. Our recovery of oral microbial and putative dietary plant DNA suggest adaptive capability. Our results suggest a spatio-temporal turnover in oral microbiome and putative dietary plant DNA, which might be explained by ecological differences between Pleistocene France and contemporary Scandinavia. As soft tissue preservation is rare in ancient remains, dental calculus emerges as an exciting and powerful tool for reconstructing the environmental and ecological histories of both past and extinct populations.

## Introduction

With their circumpolar distribution in the Arctic and sub-Arctic, reindeer (*Rangifer tarandus*) are an iconic species uniquely adapted to some of the Earth’s harshest environments (Lin et al. 2019). Like other mammoth steppe megafauna, reindeer evolutionary history has been deeply intertwined with climatic fluctuations during the Pleistocene and Holocene, which shaped their distributions, genetic diversity, migration patterns, and ecological roles (Vereshchagin and Baryshnikov 1992; Hofreiter and Stewart 2009). As a keystone species, reindeer have influenced tundra ecosystems for millennia, while also serving as a vital resource for human populations, from prehistoric hunters to indigenous herders who remain dependent on the semi-domesticated reindeer herds for resources (Müller-Wille et al. 2006; Mustonen 2022). Past work on fossil and genetic evidence has provided insights into the reindeer’s substantial climate-driven range fluctuations throughout the Pleistocene and more recent northward range shift in the Holocene (Flagstad and Røed 2003; Lorenzen et al. 2011; Sommer et al. 2014; Hold et al. 2024).

In dental calculus, the mineralised dental plaque that continuously calcifies during an individual’s lifetime (Jin and Yip 2002; Key et al. 2017), the DNA of microbes and other materials present in the oral environment become encased and can be preserved over thousands of years (Mann et al. 2018). This property makes dental calculus a promising material for studying the temporal evolution of oral microbiomes (Warinner et al. 2015; Brealey et al. 2020; Modi et al. 2020) and for determining past dietary components that can also be preserved in the calculus (Moraitou et al. 2022). For example, ancient dental calculus has been used to track changes in microbiome diversity, especially in humans (Granehäll et al. 2021; Gancz et al. 2023) and their close relatives (Weyrich et al. 2017; Fellows Yates et al. 2021). Despite offering a window into the past of oral host-associated microbiomes, the main focus of dental calculus research so far has been on ancient humans and other primates (Warinner et al. 2014; Warinner et al. 2015; Modi et al. 2020; Fellows Yates et al. 2021; Granehäll et al. 2021; Moraitou et al. 2022). As a consequence, there are only a few molecular studies investigating dental calculus of non-human animals (Ottoni et al. 2019; Brealey et al. 2020; Brealey et al. 2021; Ozga and Ottoni 2023; Moraitou et al. 2025).

Historical changes in microbiome composition of non-primates have not been studied so far. Dental calculus from ruminants may present particularly good opportunities to gain new insights into host-associated microbiomes that do not otherwise survive *post-mortem*. Ruminants, like reindeer, regurgitate their rumen content into their oral cavity for repeated mastication. This way microbiomes from the rumen and reticulum compartments of the digestive tract are introduced into the oral environment (Kittelmann et al. 2015; Tapio et al. 2016; Amin et al. 2021). There they may become incorporated into dental calculus (Brealey et al. 2020). Previous studies on the rumen microbiome of reindeer found high abundances of methanogenic archaea (Sundset et al. 2009a), numerous novel rumen uncultured bacteria (RUGs) (Glendinning et al. 2021), and yielded 18 metagenome-assembled genomes (MAGs) characterized to the species level (Kamenova et al. 2023; Kamenova et al. 2025). In a pioneering study of the dental calculus of reindeer, (Brealey et al. 2020) found that reindeer oral microbiome diversity is overall lower, with greater numbers of uncharacterized taxa, than that of other mammals like the brown bear (*Ursus arctos*) and the eastern gorilla (*Gorilla beringei*). Several typical mammalian oral microbiome members, e.g. *Streptococcus,* were determined to be almost entirely absent. Further, they identified reindeer-specific *Methanobrevibacter* related to rumen-associated species such as *M. ruminantium* and *M. olleyae*, which indicates that some portion of the rumen microbiome is captured in reindeer dental calculus (Brealey et al. 2020). Finally, Brealey et al. (2020) identified DNA from several known dietary items of reindeer, such as willow (*Salix sp.*), suggesting that dental calculus may also reveal dietary differences among historical populations.

Here, we used ancient dental calculus material from the teeth of reindeer (*Rangifer tarandus,* Linnaeus, 1758) to generate ancient metagenome-assembled genomes (MAGs). We demonstrate the suitability of paleo-genomic dental material for providing information about the oral microbiomes of prehistoric mammals and also explore the dietary information contained in ancient reindeer dental calculus, focusing on plant and lichen DNA identified in the metagenomes. We investigate the stability of the wild animal oral microbiome and dietary conservation across large temporal scales by comparing ancient metagenomes to 21 dental calculus metagenomes generated from contemporary wild reindeer. Our aim is to establish dental calculus preserved DNA as a tool to study mammalian ecology and evolution during the Pleistocene.

## Materials and Methods

### Ancient reindeer dental calculus sampling

The ancient reindeer material used in this study consists of 19 museum specimens loaned from the French National Museum of Prehistory (Musée National de Préhistoire, MNP) in Les Eyzies, Dordogne, France (see Table 1). The material was excavated from Cro-Magnon settlement sites situated in rock shelters (French: *abris*), and is believed to have been transported there from off-site hunting grounds. The archaeological sites are situated in the Vézère Valley in the near vicinity of the museum in Department Dordogne. The Dordogne region was likely a glacial refugium during the last glacial maximum (Sommer and Nadachowski 2006). Five samples were excavated in the early twentieth century by J. A. Le Bel and J. Maury from middle and upper Magdalenien layers from Abri Laugerie-Basse (sample code ‘LB’, (Maury and Le Bel 1925)). Another 14 samples (sample code ‘U’) were excavated in the early twentieth century by Denis Peyrony either at Abri de la Madeleine (middle and upper Magdalenien) or Abri Laugerie-Haute Ouest (Solutrean), but the exact records have been lost to time, therefore attribution to the sites remains uncertain. The archaeological context does not allow precise individual assignment of bones or teeth. Therefore, we exclusively used lower right third molars (M_3_) of which reindeer typically only have one, to ensure that only unique individuals were included in the study.

**Table 1.**
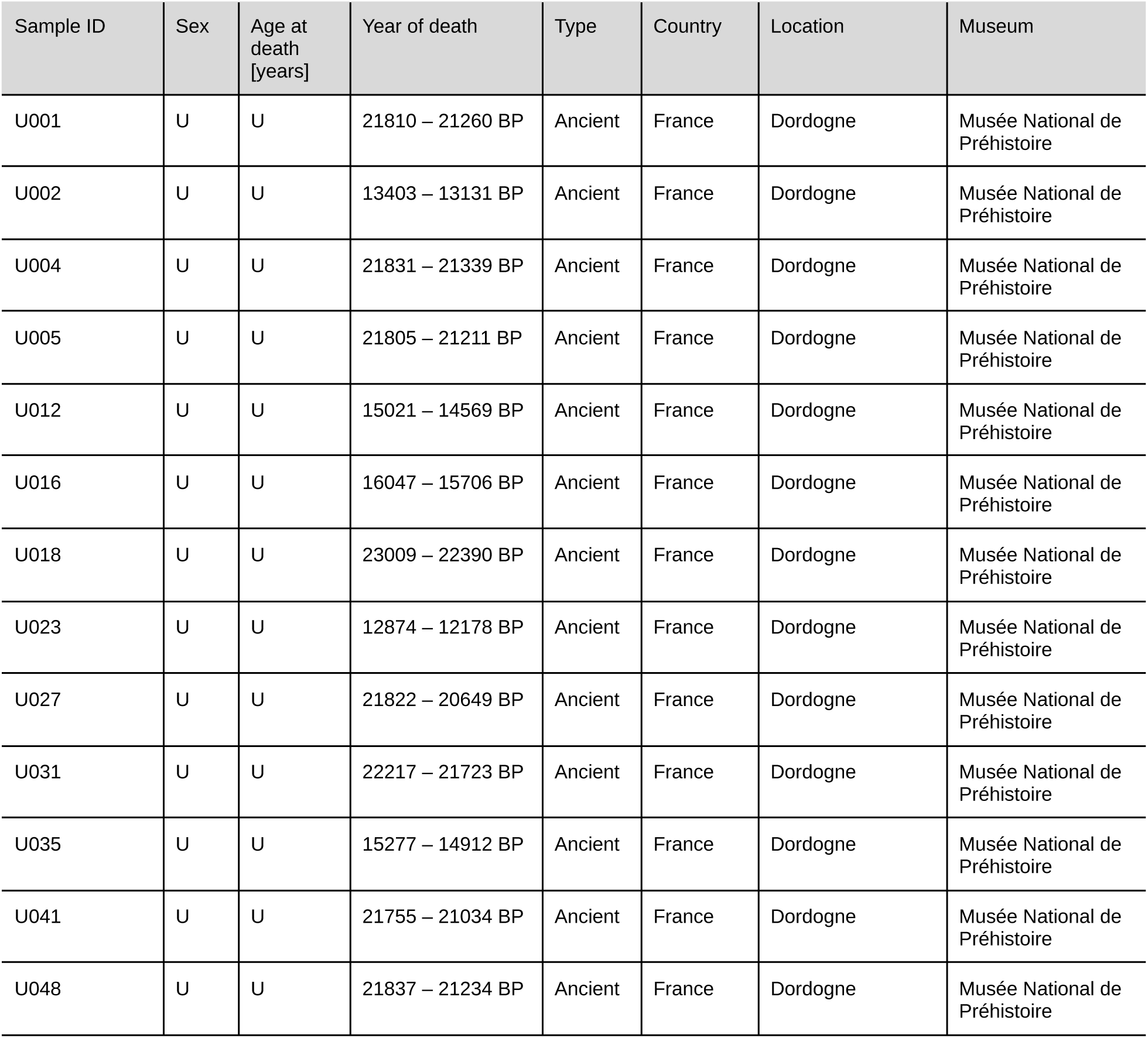
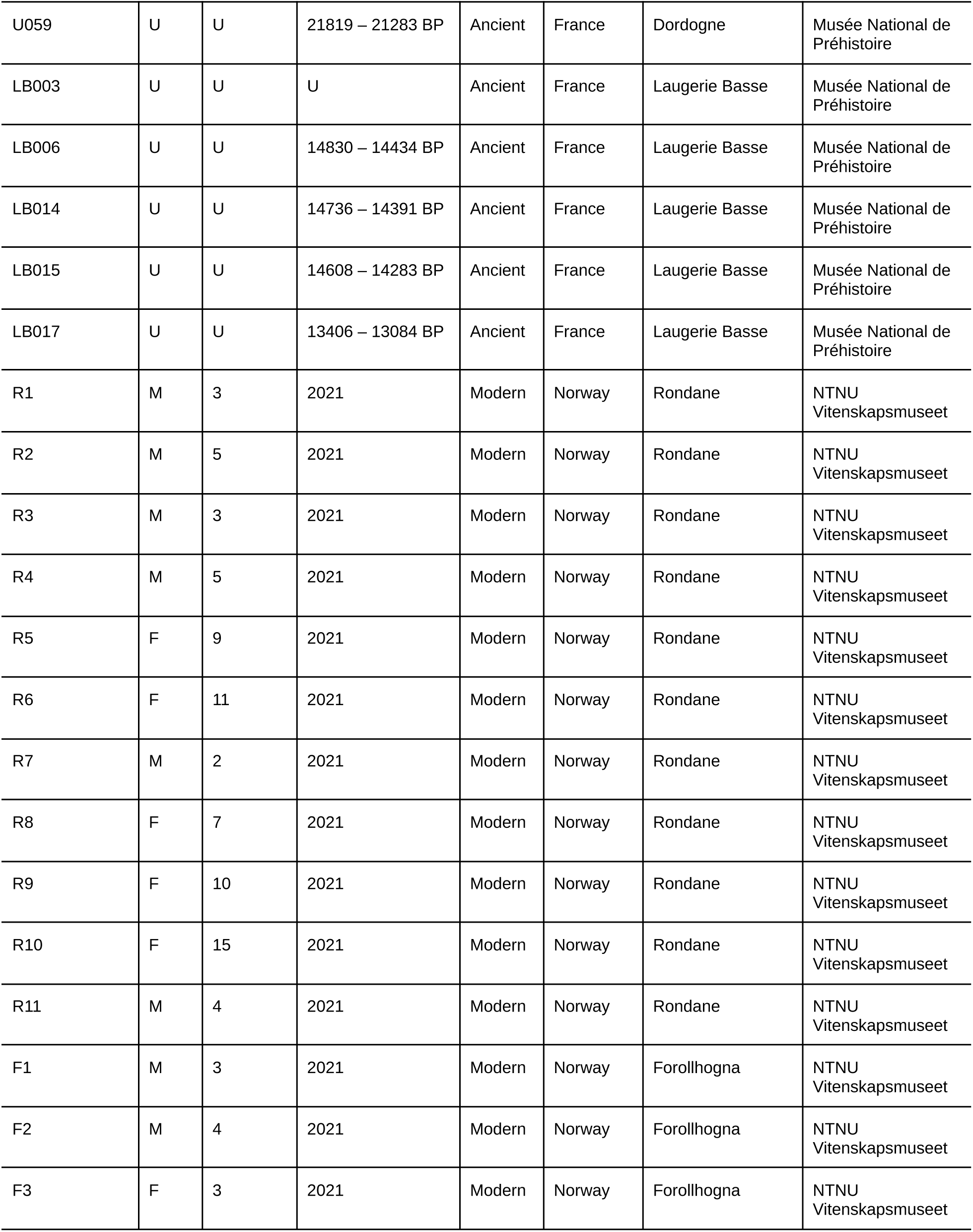
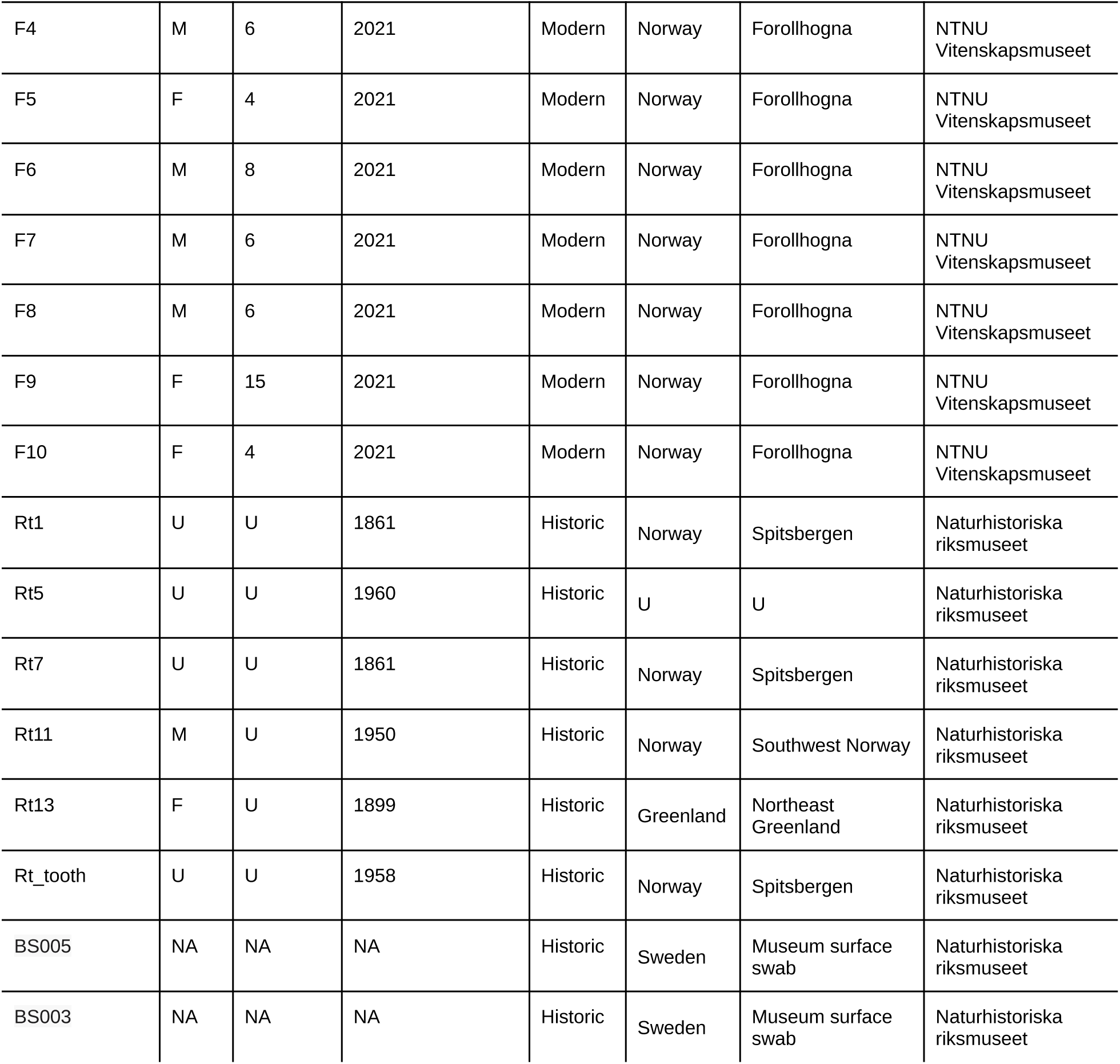
Overview of samples in this study. Laboratory negative controls (blanks) omitted. Year of death is reported in calibrated years before present (calBP) for ancient samples calculated with OxCal 4.4.4 using the calibration curve IntCal20. Year of death is reported as the calendar year for historic and modern samples. U =unknown, M = male, F = female.

| Sample ID | Sex | Age at death [years] | Year of death | Type | Country | Location | Museum |
| --- | --- | --- | --- | --- | --- | --- | --- |
| U001 | U | U | 21810 – 21260 BP | Ancient | France | Dordogne | Musée National de Préhistoire |
| U002 | U | U | 13403 – 13131 BP | Ancient | France | Dordogne | Musée National de Préhistoire |
| U004 | U | U | 21831 – 21339 BP | Ancient | France | Dordogne | Musée National de Préhistoire |
| U005 | U | U | 21805 – 21211 BP | Ancient | France | Dordogne | Musée National de Préhistoire |
| U012 | U | U | 15021 – 14569 BP | Ancient | France | Dordogne | Musée National de Préhistoire |
| U016 | U | U | 16047 – 15706 BP | Ancient | France | Dordogne | Musée National de Préhistoire |
| U018 | U | U | 23009 – 22390 BP | Ancient | France | Dordogne | Musée National de Préhistoire |
| U023 | U | U | 12874 – 12178 BP | Ancient | France | Dordogne | Musée National de Préhistoire |
| U027 | U | U | 21822 – 20649 BP | Ancient | France | Dordogne | Musée National de Préhistoire |
| U031 | U | U | 22217 – 21723 BP | Ancient | France | Dordogne | Musée National de Préhistoire |
| U035 | U | U | 15277 – 14912 BP | Ancient | France | Dordogne | Musée National de Préhistoire |
| U041 | U | U | 21755 – 21034 BP | Ancient | France | Dordogne | Musée National de Préhistoire |
| U048 | U | U | 21837 – 21234 BP | Ancient | France | Dordogne | Musée National de Préhistoire |
| U059 | U | U | 21819 – 21283 BP | Ancient | France | Dordogne | Musée National de Préhistoire |
| LB003 | U | U | U | Ancient | France | Laugerie Basse | Musée National de Préhistoire |
| LB006 | U | U | 14830 – 14434 BP | Ancient | France | Laugerie Basse | Musée National de Préhistoire |
| LB014 | U | U | 14736 – 14391 BP | Ancient | France | Laugerie Basse | Musée National de Préhistoire |
| LB015 | U | U | 14608 – 14283 BP | Ancient | France | Laugerie Basse | Musée National de Préhistoire |
| LB017 | U | U | 13406 – 13084 BP | Ancient | France | Laugerie Basse | Musée National de Préhistoire |
| R1 | M | 3 | 2021 | Modern | Norway | Rondane | NTNU Vitenskapsmuseet |
| R2 | M | 5 | 2021 | Modern | Norway | Rondane | NTNU Vitenskapsmuseet |
| R3 | M | 3 | 2021 | Modern | Norway | Rondane | NTNU Vitenskapsmuseet |
| R4 | M | 5 | 2021 | Modern | Norway | Rondane | NTNU Vitenskapsmuseet |
| R5 | F | 9 | 2021 | Modern | Norway | Rondane | NTNU Vitenskapsmuseet |
| R6 | F | 11 | 2021 | Modern | Norway | Rondane | NTNU Vitenskapsmuseet |
| R7 | M | 2 | 2021 | Modern | Norway | Rondane | NTNU Vitenskapsmuseet |
| R8 | F | 7 | 2021 | Modern | Norway | Rondane | NTNU Vitenskapsmuseet |
| R9 | F | 10 | 2021 | Modern | Norway | Rondane | NTNU Vitenskapsmuseet |
| R10 | F | 15 | 2021 | Modern | Norway | Rondane | NTNU Vitenskapsmuseet |
| R11 | M | 4 | 2021 | Modern | Norway | Rondane | NTNU Vitenskapsmuseet |
| F1 | M | 3 | 2021 | Modern | Norway | Forollhogna | NTNU Vitenskapsmuseet |
| F2 | M | 4 | 2021 | Modern | Norway | Forollhogna | NTNU Vitenskapsmuseet |
| F3 | F | 3 | 2021 | Modern | Norway | Forollhogna | NTNU Vitenskapsmuseet |
| F4 | M | 6 | 2021 | Modern | Norway | Forollhogna | NTNU Vitenskapsmuseet |
| F5 | F | 4 | 2021 | Modern | Norway | Forollhogna | NTNU Vitenskapsmuseet |
| F6 | M | 8 | 2021 | Modern | Norway | Forollhogna | NTNU Vitenskapsmuseet |
| F7 | M | 6 | 2021 | Modern | Norway | Forollhogna | NTNU Vitenskapsmuseet |
| F8 | M | 6 | 2021 | Modern | Norway | Forollhogna | NTNU Vitenskapsmuseet |
| F9 | F | 15 | 2021 | Modern | Norway | Forollhogna | NTNU Vitenskapsmuseet |
| F10 | F | 4 | 2021 | Modern | Norway | Forollhogna | NTNU Vitenskapsmuseet |
| Rt1 | U | U | 1861 | Historic | Norway | Spitsbergen | Naturhistoriska riksmuseet |
| Rt5 | U | U | 1960 | Historic | U | U | Naturhistoriska riksmuseet |
| Rt7 | U | U | 1861 | Historic | Norway | Spitsbergen | Naturhistoriska riksmuseet |
| Rt11 | M | U | 1950 | Historic | Norway | Southwest Norway | Naturhistoriska riksmuseet |
| Rt13 | F | U | 1899 | Historic | Greenland | Northeast Greenland | Naturhistoriska riksmuseet |
| Rt_tooth | U | U | 1958 | Historic | Norway | Spitsbergen | Naturhistoriska riksmuseet |
| BS005 | NA | NA | NA | Historic | Sweden | Museum surface swab | Naturhistoriska riksmuseet |
| BS003 | NA | NA | NA | Historic | Sweden | Museum surface swab | Naturhistoriska riksmuseet |

All ancient samples were processed at a dedicated paleo-genomics facility at the NTNU University Museum. To remove surface contamination, the teeth were irradiated with UV light for 10 minutes on each side and subsequently decontaminated using EDTA, following the recommendations by (Brealey et al. 2020). Dental calculus powder (10-20 mg) was collected by grinding the tooth surface with a Dremel multitool equipped with a file head. To avoid outside and cross-contamination, subsampling was conducted inside a laminar flow-hood, which was decontaminated between samples with 5% (v/v) bleach and 70% EtOH. Dental calculus powder was collected using clean aluminum foil, which was exchanged between samples. All ancient samples were radiocarbon-dated at the NTNU University Museum’s National Laboratory for Age Determination (Trondheim, Norway), except sample LB003 which was subsequently removed from the metagenomic analyses (see Methods section “Metagenomic analysis”). Samples were prepared and measured as described in (Seiler et al. 2019). In summary, collagen was extracted using a modified Longin protocol (Longin 1971). Tissue pieces were cleaned in an ultrasonic bath with water and acetone to remove soil particles and fats. After demineralization with HCl, humic substances from the soil were removed with NaOH, before the samples were hydrolyzed at pH=3, 70°C overnight. The extracted gelatin was filtered through quartz filters and freeze-dried. The dried collagen was subsampled and about 3 mg was used for further treatment. The subsamples were combusted in an elemental analyzer to measure CN contents for quality control and the CO2 was captured from the gas stream and reduced to graphite. The graphite was then measured by accelerator mass spectrometry against an oxalic acid standard (NIST SRM-4990C) that had followed the same treatment. Radiocarbon dates were calibrated with OxCal v4.4.2 (Bronk Ramsey 2009) and the terrestrial calibration curve IntCal20 (Reimer et al. 2020).

### Historical reindeer samples

We complemented our dataset of ancient reindeer dental calculus with previously published short-read data from specimens of six historical reindeer that died between 1856 and 1960 (mainland Norway n = 1, Svalbard n = 3, Greenland n = 1, Unknown n = 1, Table 1) and are stored in the collections of the Swedish Museum of Natural History (Stockholm) (Brealey et al. 2020). One of these samples (“Rt_tooth”) was a fragment of reindeer tooth crown which serves as control for tooth surface and external contamination in contrast to authentic dental calculus. Two samples (“BS005”, “BS003”), surface swabs from the museum storage area, were taken as controls of environmental contamination. The negative blank controls from this study were also downloaded and processed, to account for study-specific laboratory contamination. The sequence data was retrieved from the European Nucleotide Archive (ENA) at EMBL-EBI under accession number PRJEB33363 (https://www.ebi.ac.uk/ena/browser/view/PRJEB33363).

### Modern reindeer dental calculus sampling

In order to contextualize the results gained from the ancient and historical samples, we obtained 21 dental calculus samples from present-day reindeer populations. This modern wild reindeer material (see Table 1) was collected by hunters as part of the Norwegian national monitoring program for cervids, managed by the Norwegian Institute for Nature Research (NINA) on behalf of the Norwegian Environment Agency. The reindeer were hunted from wild populations in Rondane Sør (61.88°N, 9.80°E; *n*=11) and Forollhogna (62.68°N, 10.79°E; *n*=10), two of six monitoring areas for wild reindeer in mainland Norway, during the annual reindeer hunt between August and September 2021. Hunters register information, including kill date, location, and animal sex in a national database (www.hjorteviltregisteret.no). Hunters deliver the pre-cleaned and dried lower mandibles, which are then measured and used to assess age at death at NINA in Trondheim, Norway. Age for individuals two years and older was determined based on readings of cementum annuli in decalcified and stained longitudinal sections (Veiberg et al. 2020). To allow easy and non-destructive extraction of the incisors used for age determination, the jaws were boiled for approximately 90 minutes. After boiling, dental calculus samples were collected from one half of an air-dried mandible of each reindeer individual in the wild animal processing room at NINA. During sampling, all surfaces were decontaminated with bleach and then wiped with ultra-pure molecular grade water. A fresh aluminum foil sheet was used as the working surface for each sample. Gloves and scalpel blades were changed between each sample and protective clothing was worn, including one-way lab coats, double gloves taped to the lab coat so that no skin was exposed during sampling, and face masks. Material was collected by scraping off the dental calculus deposits from all teeth in the mandible, including both lingual and buccal sides. In most cases, the majority of the material was collected from molars and premolars, only rarely the incisors were included as well, aiming for ca. 50 mg of material from each individual. Afterwards, samples were stored in microcentrifuge tubes at room temperature for 1 week before being frozen at -20°C until DNA extraction.

### DNA extraction, library building and sequencing

For all samples, all pre-PCR steps were performed in laboratories that are free of PCR products. Ancient samples were additionally processed in separate, dedicated, positively pressurized clean-room facilities, while modern samples were processed in a conventional molecular biology laboratory.

DNA extraction from dental calculus powder was conducted using a protocol especially developed by (Brealey et al. 2020) for non-human, mammalian dental calculus based on an adaptation of the protocol from (Dabney et al. 2013). Each extraction batch was accompanied by negative blank controls containing no sample material, for the subsequent identification of laboratory contaminants. The extracted modern genomic DNA was sheared to a mean fragment length of 350 to 500 bp using a Covaris ME220 focused ultrasonicator. For each sample and the extraction blanks, 20 μL of DNA extract were built into single-stranded libraries, following the *Santa Cruz Reaction* library preparation method (Kapp et al. 2021). DNA extraction blanks were included, as well as a library blank (no-template control during library building). The optimal number of PCR cycles for each library was determined via qPCR on a QuantStudio3 instrument (ThermoFisher), then each library was amplified using a dual indexing approach (Kircher et al. 2012). The indexing PCR buffer mix consisted of (per sample) 0.8 µL 25 mM dNTPs (final conc. 0.2 mM), 5 U/µL AmpliTaq Gold polymerase (final conc. 0.05 U/µL), 10 uL 10X AmpliTaq Gold buffer (final conc. 1X), 10 µL 25 mM MgCl2 (final conc. 2.5 mM), 0.8 µL 50 mg/mL BSA (final conc. 0.4 mg/mL) and 63.4 µL ultra-pure molecular grade H_2_O, 10 µL DNA template and 2 µl of 100 µM dedicated P5 and P7 index each. Based on the qPCR results, in the indexing PCR modern samples were amplified with 10 cycles, modern blanks with 14 cycles, ancient samples with 5 to 14 cycles and ancient blanks with 21 cycles. The indexing PCR settings consisted of a 10 minute hot-start at 95 °C, followed by a sequence of 30 seconds at 95 °C, 60 seconds at 60 °C, and 45 seconds at 72 °C, repeated for the given number of cycles. Final extension lasted for 5 minutes at 72 °C. The PCR product was purified using SeraMag beads in bead:DNA ratio of 9:10 and eluted in 33 µL Qiagen EBT buffer. The amplified libraries were quantified on 4200 TapeStation instrument (Agilent Technologies) using High Sensitivity D1000 ScreenTape. The libraries were then pooled in equimolar concentrations and subsequently subjected to several rounds of PE 150 bp sequencing on the Illumina NovaSeq 6000 platform (Novogene UK).

### Sequencing data processing

Sequence data from the pooled libraries were demultiplexed according to their unique dual-index combinations. Read pairs with the default overlap length of ≥ 11 bp were collapsed, residual adapter sequences were removed and reads shorter than 25 bp were discarded with AdapterRemoval v2.2.4. Trimmed sequencing reads were mapped against a caribou (*R. t. caribou*) reference genome assembly (Taylor et al. 2019) with the PALEOMIX pipeline v1.7 (Schubert et al. 2014) using the ‘mem’ algorithm of the Burrows-Wheeler Aligner (BWA) v0.7.16a (Li 2013) and no mapping quality (MAPQ) score filtering. We discarded reads that mapped to the caribou reference and extracted unmapped reads with *samtools* v1.12. From these unmapped reads, PCR duplicate reads were removed with *seqkit* v2.3.0. We then mapped only the collapsed reads against the human nuclear and mitochondrial reference genomes (genbank accession number GCF_000001405) to further remove possible contamination from processing the samples. We chose to only use the collapsed reads because we expect the DNA in dental calculus to be fragmented, therefore longer fragments are likely to be modern contamination (see (Brealey et al. 2020)). As before, we discarded reads mapping to the reference and extracted reads with samtools. We considered these unmapped reads to be of putative microbial origin (henceforth referred to as ‘*microbial reads*’).

### Metagenomic analysis

#### MAG assembly

In order to accurately characterize the microbiome at lower taxonomic levels, we generated metagenome-assembled genomes (MAGs) following the *metaWRAP* v1.3.2 pipeline (Uritskiy et al. 2018), processing ancient and modern samples separately throughout. First, we assembled the reads into longer contigs. We opted for a co-assembly strategy using *MEGAHIT* v1.2.9 (Li et al. 2015), in which reads from multiple samples are assembled together, since this strategy increases the usefulness of very low coverage data. We ran two co-assemblies, one each for all ancient samples (including the downloaded historical metagenomes, ‘ancient assembly batch’) and all modern samples (‘modern assembly batch’), with a minimum contig length of 1000 (*--min-contig-len* 1000) base pairs (bp) and the ‘meta-sensitive’ preset (*--presets meta-sensitive*). We chose this grouping of samples since historic samples and ancient samples share characteristic post-mortem damage patterns that the freshly collected modern samples do not have. To avoid polluting the co-assembly with putative contaminant reads, microbial reads from blank control samples were not included in the assemblies, but were mapped back to the assemblies during binning to identify putative laboratory contaminants (see below).

We checked for reads mapping to the common Illumina sequencing control, phiX 174, by mapping the microbial reads against the phiX reference genome (RefSeq acc. No. GCF_000819615.1) with PALEOMIX v1.7 as described above. Since the total number of reads mapping to PhiX in each sample were negligible (min = 0, max = 10, mean = 0.8) we did not remove those reads before further processing.

To account for miscalls caused by aDNA damage in the ancient co-assembly, we used the method described in (Klapper et al. 2023), which is implemented as part of the nf-core/mag pipeline (v2.5.4) (Krakau et al. 2022) within the nf-core framework (Ewels et al. 2020). Since the aDNA subworkflow is currently not implemented to work with co-assembly input as of version 2.5.4, we concatenated our input reads into a single file and ran the pipeline in single-assembly mode. Apart from the aDNA subworkflow, all other parts of nf-core/mag were skipped.

The assembled contigs were then processed with the *Anvi’o* pipeline (Eren et al. 2021). Open-reading frames within contigs were identified with *Prodigal* (Hyatt et al. 2010). Contigs were then annotated using Hidden-Markov-Models (HMMs), searching against *Anvi’o*’s collections of bacterial single-copy genes, as well as KEGG (Kanehisa and Goto 2000) and Pfam (Mistry et al. 2021) functional annotations. *Anvi’o* sample and depth profiles for binning were generated by mapping the original microbial reads from each sample, including blank samples, against all contigs from their respective co-assembly with the *mem* algorithm in *BWA* v0.7.17-r1188, in a manner similar to competitive mapping, where the reads are mapped against a reference concatenated from multiple reference genomes (Feuerborn et al. 2020). The resulting bam files were then further profiled with *anvi-profile* using a minimum contig length of 1000 bp. The resulting sample profiles were hierarchically clustered based on depth profiles into a single merged profile database for downstream analysis with *Anvi’o*.

To assess ancient DNA degradation, we ran *mapDamage2* (Jónsson et al. 2013) on the microbial reads for every ancient and historical sample with the collection of *Anvi’o* contigs acting as reference. In addition, in the case of the six taxa that appeared in both ancient and modern samples, we mapped the ancient microbial reads against the respective MAGs with the *paleomix* v1.7 bioinformatic pipeline (same settings as above), including built-in damage assessment with *mapDamage2* with default settings. We used the decOM pipeline (Duitama González et al. 2023) with the default decOM source matrix to track the putative sources of the metagenomic sequences in our dental calculus. The pipeline was run separately for the ancient and modern sample groups.

To ensure high accuracy for taxonomic classification, we binned (i.e. sorting the contigs into likely operational taxonomic units (OTUs)) contigs with three independent, automated metagenomic binning softwares: *Metabat* v2.12.1 (Kang et al. 2014) with default parameters*, Maxbin* v2.0 (Wu et al. 2016), with default parameters, and *Concoct* v1.1.0 *(Alneberg et al. 2014),* with default parameters and number of clusters set to 100, which we chose based on preliminary analysis of the data. The results of the three binning softwares were refined and dereplicated with *metaWRAP*’s built-in refinement module, using a 50% completion threshold and a 20% redundancy/contamination threshold as calculated by *checkM* (Parks et al. 2015).

#### Short read classification with KRAKEN

In order to test how our MAG analysis compares to a read-based classification method, we analyzed the dataset with *KrakenUniq* (Breitwieser et al. 2018; Pockrandt et al. 2022). We were interested in the quantity of reads and therefore how much of the community diversity is captured by each method. Furthermore, we wanted to assess how abundance and community composition is estimated by the two approaches, and whether they change when focusing on only oral or gastro-intestinal taxa (GIT) microbial taxa. To facilitate a comparison between the methods, we used exactly the same input reads as for the MAG analysis (see above). Reads were searched against the 2023 version of GenBank Microbial Nucleotide database (microbial NT). To be assigned species-level taxonomy, reads required a minimum of 1000 unique k-mers per taxon per sample, as well as a minimum of 50 reads per species per sample. Relative abundances of taxa at different taxa ranks were generated from a MPA style report in *KrakenUniq*. To retrieve the most up-to-date taxonomy nomenclature, the taxon identification tags (taxIDs) of classified taxa were additionally searched against the NCBI Entrez database. Only taxa within Bacteria or Archaea were retained for downstream analysis.

#### Statistical analysis of microbial communities

A Wilcoxon test in *R* (v4.2.3) (R Core Team 2021) was used to test for differences in MAG completion, redundancy and guanine-cytosine (GC) content between the two co-assembly batches. The finished MAGs were then taxonomically classified with the Genome Taxonomy Database (release 207_v2) using the associated Genome Taxonomy Database Toolkit (*gtdb-tk* v2.2.4, (Chaumeil et al. 2019; Chaumeil et al. 2022)).

We then used the ‘compare’ function of *dRep* (Olm et al. 2017) to find MAGs which were assigned the same taxonomy in the modern and ancient batches. MAGs were considered of identical taxonomy when the primary algorithm (*Mash*, (Ondov et al. 2016)) assigned more than 90% shared average nucleotide identity (ANI) and the secondary clustering algorithm (*ANImf*, (Richter and Rosselló-Móra 2009)) assigned at least 90% shared ANI. If one or both methods yielded lower results than the threshold, we considered the MAGs sufficiently distinct to represent different taxa. Summary statistics were computed by *Anvi’o*. In *Anvi’o*, abundance is calculated as the mean coverage of a contig divided by the mean coverage of all contigs in that sample. In addition, *Anvi’o* calculates a detection value, which equals the proportion of nucleotides in any one bin that have at least 1X coverage in a sample. We considered a bin as sufficiently well detected in a sample when the detection value was above 0.3.

The *Anvi’o* summary statistics for detected MAGs in the modern and ancient datasets, including blank controls, were then combined and analyzed in *R* (v4.2.3) (R Core Team 2021) with the *phyloseq* (McMurdie and Holmes 2013)*, decontam* (Davis et al. 2018) and *maaslin2* (Mallick et al. 2021) packages. *Decontam* was used to identify and remove likely laboratory contaminant taxa based on their prevalence in the blank controls, using the default probability threshold of 0.1. After removing contaminants, we log-transformed filtered abundance values and used the *vegan (Oksanen et al. 2013)* package to calculate a matrix of dissimilarity indices (based on Euclidean distance) which was used as input for a principal coordinate analysis (PCoA) with the *ape* package (Paradis and Schliep 2019), as well as to conduct a permutational multivariate analysis of variance (PERMANOVA) using age class (ancient, historical, modern) and sequencing depth (number of microbial reads) as variables. To explore possible differences between the sampling areas of the modern reindeer populations, we additionally conducted a PERMANOVA using individual age, area and number of microbial reads as variables. We calculated a projected PCA onto the PCoA with *R* code developed by Matthias Grenié (https://gist.github.com/Rekyt/), which itself uses the method from (Legendre and Legendre 1998). This method is similar to the native method in the *ape* package, but was chosen over the latter for compatibility with *R’s ggplot2* package (Wickham et al. 2016). We investigated whether microbiome community composition is affected by the age of the host individual by constructing a PCoA of modern samples with known ages at death. Further, we used the *vegan* package for an analysis of multivariate homogeneity of group dispersions and a Tukey multiple comparisons of means with a 95% family-wise confidence level. We generated a clustered heatmap of filtered, log-transformed abundances with the *pheatmap R* package (Kolde 2019). For a differential abundance analysis between ancient and modern samples, *Maaslin2* was used with the not-log-transformed abundance values with minimum abundance of 0 and a minimum prevalence of 1%, and the default linear model (LM) method with *Maaslin2’s* built-in log-transformation (log2) but no additional normalization. Blank samples, historical samples and the samples LB003, U035 and U027 were removed from this analysis due to the lack of authentic microbial reads.

Given the more sensitive (but likely less accurate) classification of *KrakenUniq*, we employed a more conservative contamination investigation compared to the MAG analysis, where we could use damage and binning statistics to verify findings. We used decOM to exclude samples with < 5% cumulative oral proportions (Supp. Fig. 1A). Samples were filtered by read count, removing any sample with < 100,000 species-level assigned prokaryotic reads (Supp. Fig. 1B). Using our control samples, we measured contaminant prevalence (threshold = 0.5) and frequency (threshold = 0.1) with *decontam*, taking into account sampling batch (ancient, historical, modern) with the option *batch.combine = “minimum”* (Supp. Fig. 2). We removed all taxa with higher relative abundance in museum controls (swabs, tooth) than dental calculus, including taxa that were only found in controls and not at all in calculus (Supp. Fig. 3). Many of the removed taxa are known environmental and human-skin associated species, like *Cutibacterium acnes*. *Pseudomonas* spp. remained at high abundance in the reindeer tooth sample and some blanks and low abundance in some dental calculus samples, despite the previous filtration steps. Since it is a common and well known environmental and lab contaminant ((Salter et al. 2014; Weyrich et al. 2019), we filtered all *Pseudomonas* taxa from our samples (Supp. Fig. 4).

For statistical analysis of the data generated with *KrakenUniq*, the read counts in the final dataset were transformed using the “robust CLR” transformation in the *microbiome* package (Lahti and Sudarshan 2012-2019) for R. Samples were projected onto a PCoA using Euclidean distances of microbiome dissimilarity, using the *ordination* function in *phyloseq*. All control samples clustered together away from the dental calculus samples (Supp. Fig. 5); thus we removed the control samples from the remainder of the Kraken analyses. PCoAs were repeated with just the dental calculus samples, using both Euclidean distances of robust CLR transformed abundances, and Jaccard distances of taxa presence/absence. To compare taxa detected in ancient vs modern samples, we generated Venn diagrams with the R package *ggVennDiagram (Gao 2025)*.

Following the procedures described in (Moraitou et al. 2022), we used species- and genus-level sets of predefined taxa lists to identify putative oral-associated taxa, based on the Human Oral Microbiome Database (accessed April 6, 2025) and other primate research (Chen et al. 2010; Fellows Yates et al. 2021). We also included a list of genus- and family-level taxa previously found in the rumen and GIT of Norwegian mainland and Svalbard reindeer (Sundset et al. 2009a; Sundset et al. 2009b; Pope et al. 2012; Salgado-Flores et al. 2016; Zielińska et al. 2016; Kamenova et al. 2023). Since no such prior information is available for the reindeer oral microbiome, we used a combination of the primate-derived oral-associated taxa list, along with ‘core reindeer dental calculus microbes’ based on our modern dental calculus samples. Following the methodology from (Fellows Yates et al. 2021) a taxon was considered a core taxon if it was detected in 50% of individuals of the target population (>= 10 modern individuals). Based on these taxa lists, we calculated the proportion of reads assigned to reindeer oral and GIT microbes (or both, when a taxon appeared in both lists) using either *KrakenUniq* or the MAG approach. We also repeated the PCoAs on the dental calculus samples for the *KrakenUniq* dataset, using only reindeer oral taxa or only reindeer GIT taxa.

To compare the assignment and relative abundances of taxa between the *KrakenUniq* and MAG approaches, we calculated the proportion of reads assigned by *KrakenUniq* to each taxon across all dental calculus samples, compared to the proportion of reads mapped to MAG bins for the corresponding taxon. Each taxon was summarised to the lowest taxonomic level classified by both approaches. For example, *Streptococcus chenjunshii* and *Streptococcus gallolyticus* were detected by both approaches and proportions were therefore calculated at the species-level for these taxa; however other *Streptococcus* species were detected by only one approach, and were thus summarised at the genus-level as ‘other *Streptococcus*’. This procedure was continued up to the class level. Taxa without a matching class detected by both methods were summarised together as ‘taxa only detected in one method’.

### Dietary profiling analysis

For dietary profiling, all non-host reads were competitively mapped with bowtie2 (options: -k 1000 -D 15 -R 2 -N 0 -L 22 -i S,1,1.15) against three reference databases: the NCBI RefSeq database (release 213, excluding bacteria), the PhyloNorway plant database (Wang et al. 2021), and the NCBI nt database (release 251, cf. (Sayers et al. 2025)). The PhyloNorway database was recently reported to contain substantial bacterial contamination (Oskolkov et al. 2025), despite prior efforts to pre-filter microbial reads (Wang et al. 2021). To account for this, we additionally mapped all non-host reads against a combined reference genome database comprising the GTDB microbial database and RefSeq chloroplast and mitochondrial genomes using bowtie2 (options: -k 1000 -D 15 -R 2 -N 1 -L 22 -i S,1,1.15 --np 1 --mp ‘1,1’ --rdg ‘0,1’ --rfg ‘0,1’ --score-min ‘L,0,-0.1’). We then ran the tool *getRtax* (https://github.com/genomewalker/get-reads-taxonomy) to distinguish microbial (Archaea, Virus and Bacteria) reads from eukaryotic reads. Microbial reads were removed from the alignments to the eukaryotic reference databases, after which the remaining eukaryotic alignments were merged and the BAM file headers cleaned of non-hit references using the compressbam function in *metaDMG* (Michelsen et al. 2022).

The resulting alignment files were processed with bamfilter reassign to assign reads to the most likely reference using the built-in expectation–maximization (EM) algorithm (Fernandez-Guerra et al. 2023). Alignments were then passed through *bamfilter* twice: first to obtain mapping statistics and second to apply filtering criteria, retaining alignments that (i) had an expected breadth ratio > 0.8 or (ii) showed a normalized GINI < 0.6 and normalized entropy > 0.6. We further excluded references with a mean read ANI < 95% and references with fewer than three aligned reads (using options: -g 0.6 -b 0.8 -e 0.6 -m 8G -t 4 -n 3 -A 92 -a 95 -N --include-low-detection). The *metaDMG* lca/dfit/aggregate functions were then applied to first taxonomically classify reads and parse mismatch matrices, then fit and estimate the mismatches due to DNA damage, and lastly to aggregate the resulting taxonomic profiles with the damage statistics. All taxa are presented at genus-level for Viridiplantae and Fungi. Since most fungi are likely environmental, fungi assignments were classified with *FUNGuildR (Nguyen et al. 2016)* to identify lichenized fungi, and all other fungi were excluded from further analysis. We note that this filtering step excludes many mushrooms that could be food sources for reindeer; however, we were unable to identify fungi with likely fruiting bodies from the taxonomic assignments. In addition to the above filters, we required a minimum of 100 reads and an average read length 35 bp for a taxon to be classed as detected in a sample. Potential laboratory contaminants were identified and removed using prevalence in control samples (extraction and library blanks, museum swabs, and the museum reindeer tooth sample) using *decontam* with a threshold of 0.3. In addition, taxa were excluded if their relative abundance was higher in any of the control samples, compared to any of the dental calculus samples. Finally, taxa detected in ancient samples were required to have an average damage score of 0.05 to be further considered in the dataset. Given the young age of the historical samples (1861–1960), we were unable to use DNA damage as a filtering criteria for those samples (Supplementary Table 3). For some analyses, genera with between 10 and 99 reads in a sample were classed as “present with low confidence”, regardless of damage score, due to the low numbers of plant and lichen reads in the dataset.

For analysis, we focused on genus-level taxa detections (i.e. presence/absence), rather than relative abundance in each sample, since we have no knowledge of how DNA read abundance in dental calculus relates to quantity of dietary consumption. For comparing taxa detections among age classes and modern populations, we subset the data to genera detected with high confidence in >1 sample (32 genera). Due to the small number of genera recovered and sparseness of their detections, we did not have sufficient power for alpha or beta diversity analyses. Instead, we provide a descriptive analysis of putative dietary profiles through heatmaps generated with the R package *pheatmap* and Venn diagrams generated with the R package ggVennDiagram.

## Results

### Radiocarbon dating

Radiocarbon dating placed the majority of the ancient samples at 12–23 thousand years before the present (ka BP, Table 1), a period covering the last glacial maximum (ca. 18–20 ka BP) to the end of the Pleistocene (ca. 11.7 ka BP) (Clark et al. 2009). The samples fall into two temporal groups, one older group between 21 - 23 ka BP and a younger group between 12 - 16 ka BP.

### MAG generation and authenticity

Our final dataset consisted of 21 modern and 19 ancient reindeer, as well as 6 historical samples from a previously published study (Brealey et al. 2020) (Table 1). The average number of microbial reads was 24,835,136 (min: 11,299,887, max: 38,662,827) for ancient samples, 3,642,306 (min: 47,090, max: 8,820,877) for historical samples, and 8,333,884 (min: 166,985, max: 14,451,312) for modern samples. For the ancient co-assembly batch, our *MEGAHIT* co-assembly yielded 147,775 contigs with a total length of 369,908,798 bp, with an average contig length of 2,503 bp (min = 1,000 bp (controlled by settings), max = 655,378 bp) and a N_50_ of 2,898 bp. For the modern co-assembly batch, our *MEGAHIT* co-assembly yielded 126,891 contigs with a total length of 308,341,285 bp, with an average contig length of 2,429 bp (min = 1,000 bp (controlled by settings), max = 261,408 bp) and a N_50_ of 2,699 bp. The contigs of both datasets were binned into combined 121 distinct MAGs (62 ancient, 59 modern) that met our quality thresholds. Based on prevalence in blank control samples, one contaminant MAG (*Acinetobacter fasciculus*) was removed from the ancient assembly batch and five contaminant MAGs (*Caulobacter*, *Methylobacterium*, *Nevskia, Obscuribacteraceae, Mesorhizobium terrae*) were removed from the modern assembly batch, reducing the total number of MAGs to 115 (see Supp. Table 1). *Acinetobacter fasciculus* is a member of a genus of common ancient sample contaminants (Salter et al. 2014; Eisenhofer et al. 2019) whereas *Nevskia*, *Caulobacter*, *Methylobacterium* and genus *Mesorhizobium* have been previously identified as laboratory contaminants (Salter et al. 2014). While bacteria are dominant among the 115 assembled MAGs, we also identified six species of archaea (Fig. 1). Modern MAGs had significantly higher completeness and lower redundancy than ancient MAGs but were not significantly different in their GC content (Fig. 2).

**Figure 1.**
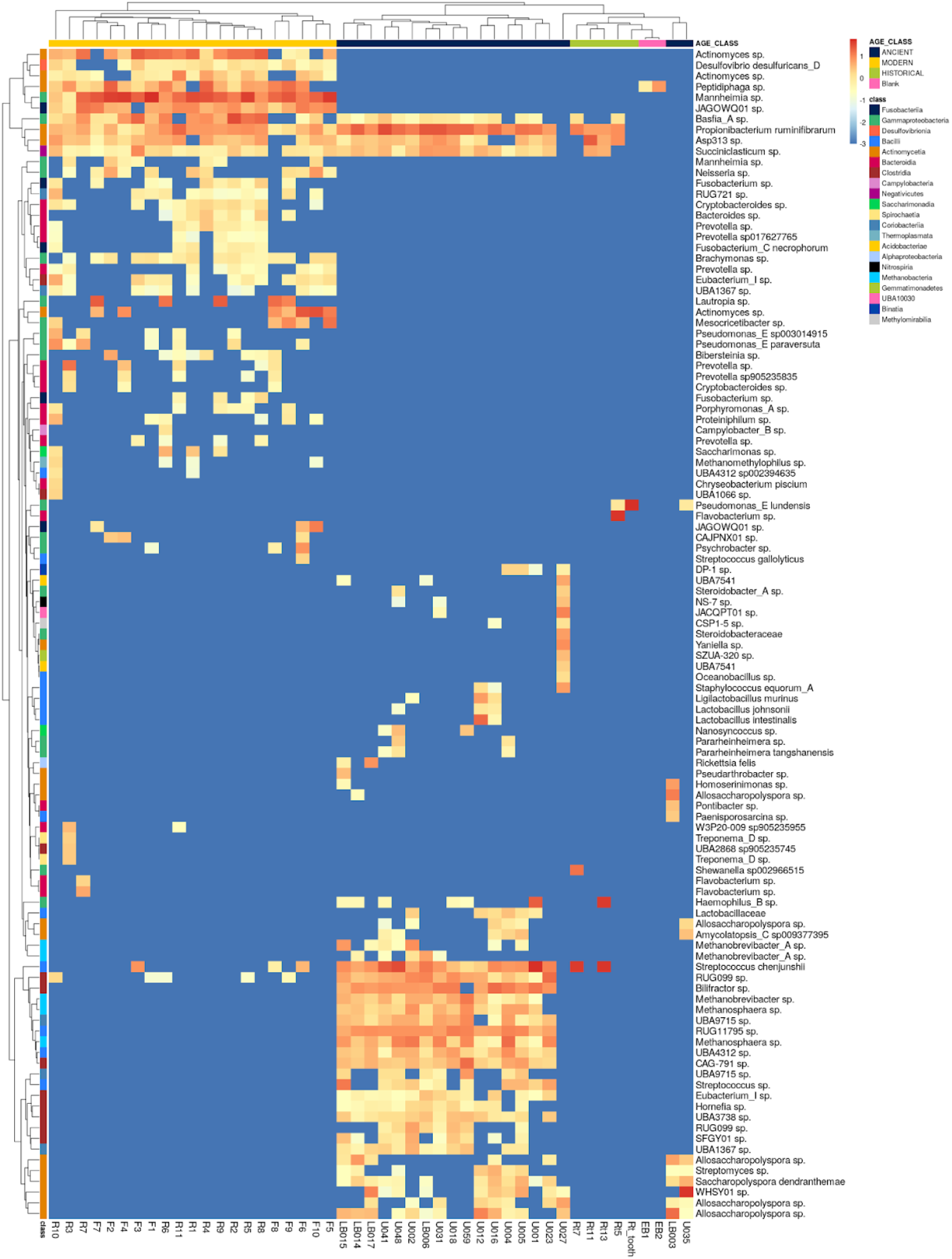
Heatmap of the abundance of the 114 MAGs in each sample. Columns are samples, rows are MAGs. Both MAGs and samples have been hierarchically clustered by abundance. Abundance is corrected for detection threshold (considering samples to be 0 abundance if under detection limit). Abundance values were log-transformed, with a pseudo-count of 0.001 for 0 abundance values (i.e. -3 after log-transformation). MAGs are labeled by their lowest-level taxonomic classification and are annotated by class-level taxonomy. Samples with no detection of any MAG after filtering are excluded from the heatmap (Rt1 and most of the blanks).

**Figure 2.**
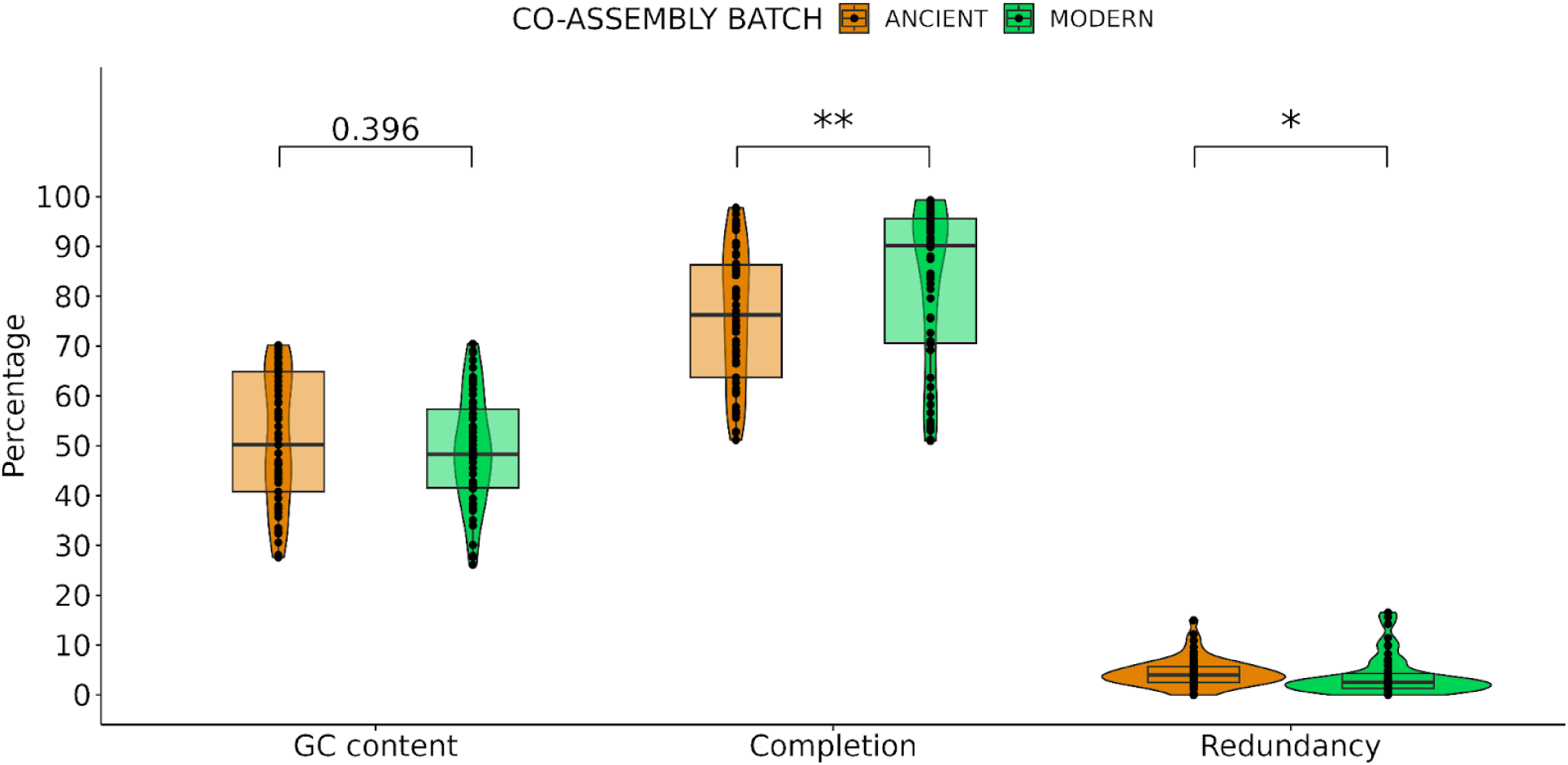
MAG assembly statistics generated by *checkM*. Relative GC content, completion and redundancy of MAGs between ancient and modern assembly batches. Each point represents one assembled MAG/bin. Point density is visualized as a violin plot. Boxes represent the 25th percentiles (lower margin), 50th percentiles (median, thick line) and 75th percentiles (upper margin) of the value distribution. Differences in batches were tested with a Wilcoxon test (p < 0.05 = *, p < 0.01 = **).

To further investigate the authenticity of the sample-specific MAGs and sequence data, we used a combination of damage pattern assessment and microbial community analysis. Prior to correcting damage with the nf-core/mag pipeline, our ancient and to a lesser degree historical samples showed high frequencies of cytosine-to-thymine transitions at the ends of reads, a damage pattern characteristic for aDNA (Supp. Fig. 6). Comparable damage patterns were detected with *mapDamage2* (Supp. Fig. 6) and the *PyDamage* (Borry et al. 2021) submodule built into the nf-core/mag pipeline’s aDNA workflow (Supp. Fig. 7). Microbial reads from the ancient samples LB003 and U035 show unusually low levels of aDNA damage given their age (Supp. Fig. 6, Supp. Fig. 8). These samples group with historical samples and blanks in the PCoA (Fig. 4). Furthermore, these samples also show a different microbial community than other ancient samples, with none of the top 20 taxa being present in LB003 (Fig. 3) and with U035 being dominated by taxon *WHSY01 sp*. of family Pseudonocardiaceae. *WHSY01 sp*. was recently discovered to colonize fossil remains and was first recovered from DNA retrieved from fossilized *Centrosaurus* bones (Saitta et al. 2019; Liang et al. 2020) a member of a more general microbial community of *post-mortem* bone colonizers involved in taphonomic processes (Liang et al. 2020). We also found *WHSY01* in lower abundance in a few other ancient samples (Fig. 1).

**Figure 3.**
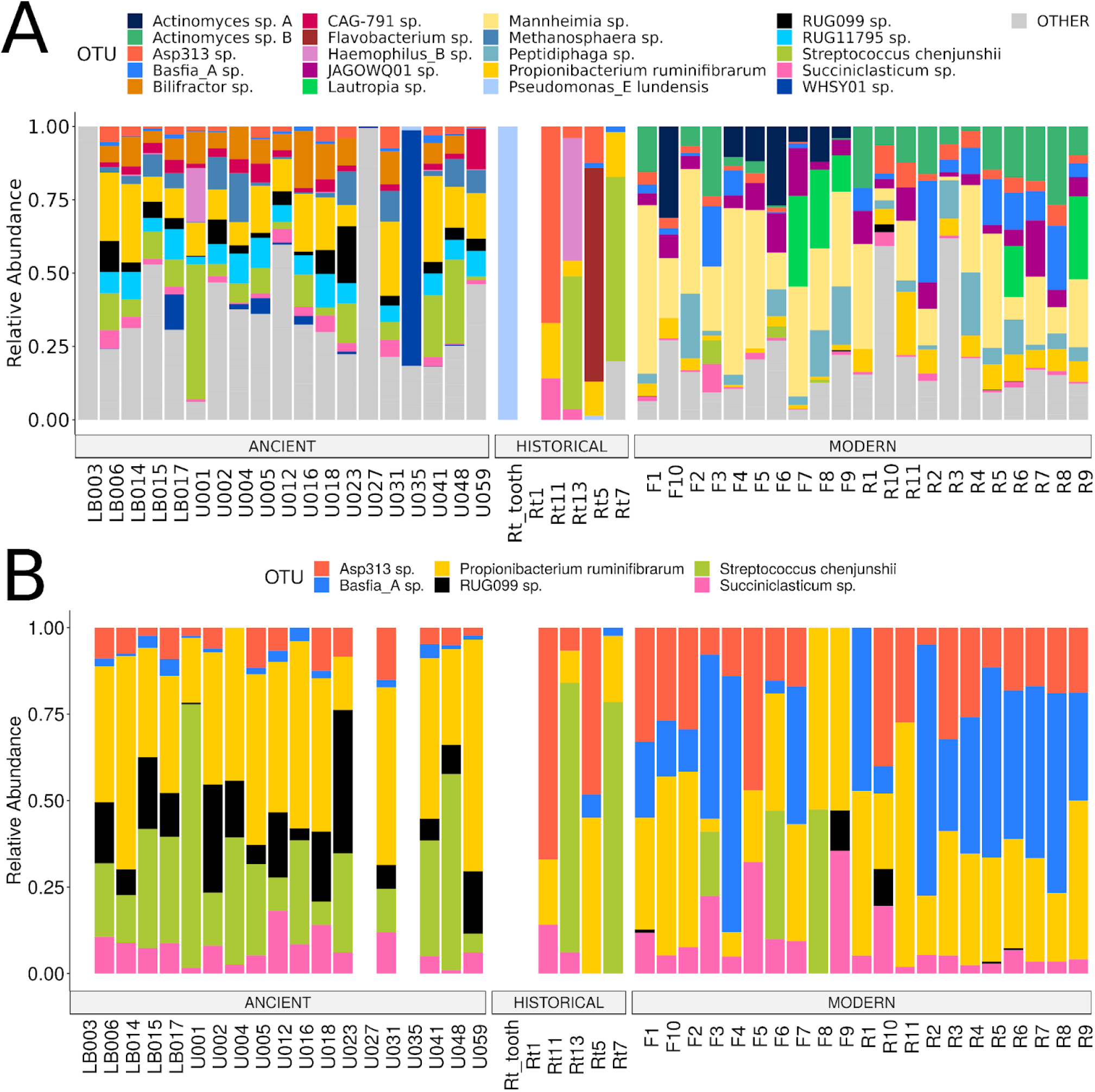
Relative abundance of selected taxa. **A:** Top 20 most abundant taxa across all samples. All other taxa are combined under ‘other’ **B:** Only taxa present in both ancient/historical and modern samples. OTU names show the lowest-level assigned taxonomy (genus and/or species). Abundance is corrected for detection threshold (considering samples to be 0 abundance if under detection limit).

**Figure 4.**
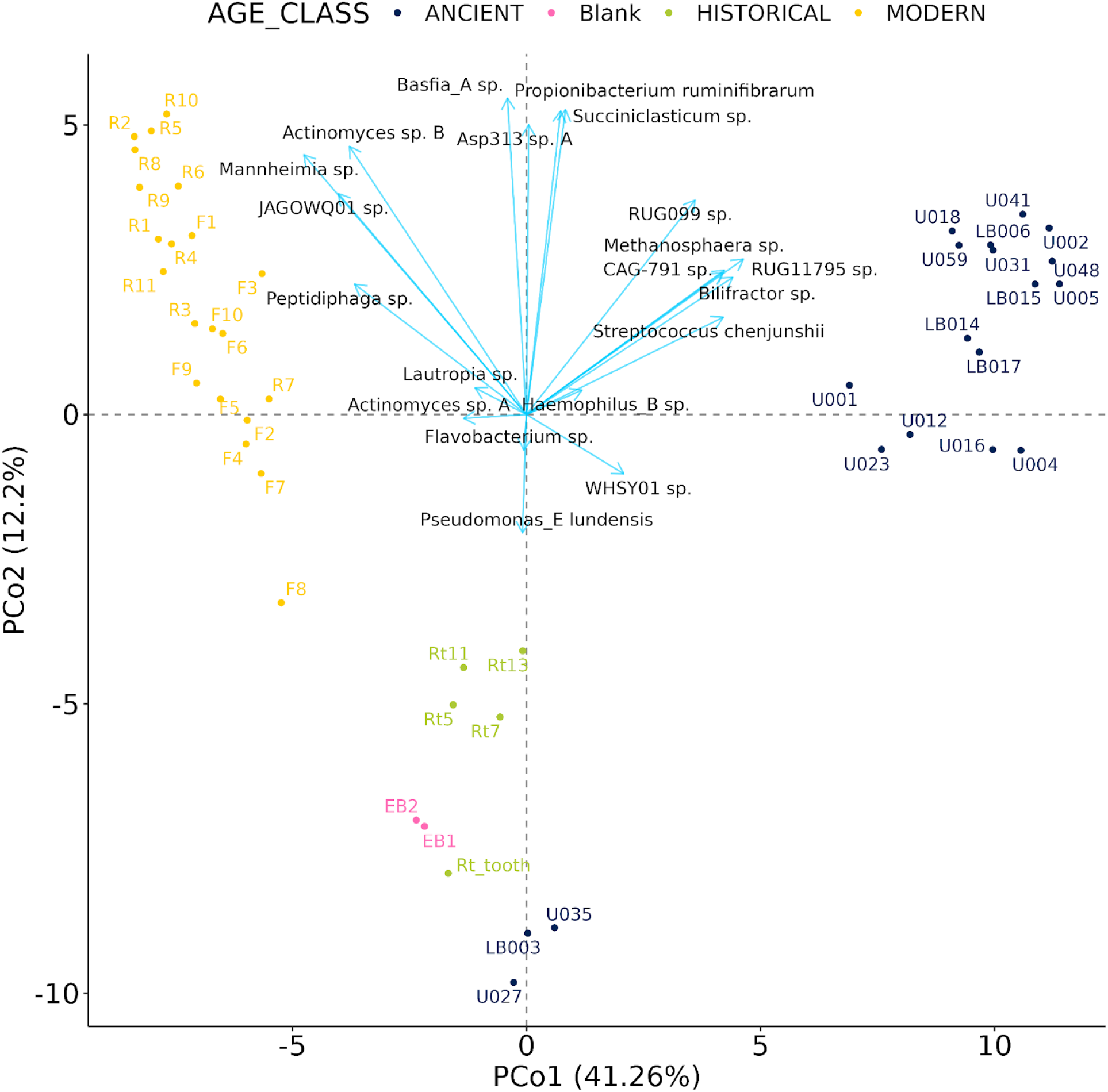
PCoA biplot of the relationship between microbial taxon abundance and samples. Samples are divided into ancient, modern, historical and laboratory blank/control samples. Abundance is corrected for detection threshold (considering samples to be 0 abundance if under detection limit.) Abundance values were log-transformed, with a pseudo-count of 0.001 for 0 abundance values (i.e. -3 after log-transformation). Samples with abundance equal to 0 are omitted. Samples are labeled in the color of their respective age class. The PCoA is overlaid with a projected principal components analysis (PCA) showing the direction of influence of the top 20 most abundant taxa on the distribution (blue arrows). Arrows are labeled with taxon names in black. Arrow lengths have been scaled by factor 20 to increase visibility.

MAGs present in both ancient and modern samples show high degrees of typical aDNA damage in the ancient samples (Supp. Fig. 9). The extraction blanks accompanying the modern sample batches have high abundances of a species in the genus *Peptidophaga*, which also occurs in relatively high abundance in most modern samples. The blanks might have been cross-contaminated by the samples; however, since we can not safely rule out that this taxon originates from a different source, we consider it contamination. *Pseudomonas lundensis (Molin et al. 1986)*, which is found in high abundance in the tooth surface control sample (Rt_tooth) from (Brealey et al. 2020), is very likely a contaminant as well, since it is commonly associated with spoiled meat, i.e. animal carcasses (Molin et al. 1986; Molin and Ternstrom 1986). These taxa were not excluded from statistical analysis, but were excluded from interpretation of results.

Using *decOM* we confirmed that most ancient, historical and modern samples contained discernible proportions of communities that resemble human oral microbiomes, with higher proportions being recognisably human oral in modern samples (Supp. Fig. 10). The proportion of soil communities was higher in ancient samples than in both historical and modern samples, consistent with their origin from excavations. A large proportion of each microbiome remained unclassified, which is expected for samples from wild, non-model organisms.

### Microbial community comparison across time

In general, we observe high differentiation by age class (ancient / historical / modern), with most taxa being found only in either the modern, ancient or historical samples (Fig. 1, Fig. 3). In ancient samples, we identified two archaeal species in the genus *Methanobrevibacter* as well as two species in the genus *Methanosphaera* (Fig. 1). These taxa are absent from modern samples. The only species of archaea identified in modern samples was from genus *Methanomethylophilus,* which in turn is absent from ancient samples (Fig. 1). In the PCoA, samples cluster distinctively into an ancient cluster, a modern cluster, and a third cluster containing blanks and historical samples, as well as the three ancient samples U027, U035 and LB003 along PC 1 (Fig. 4). This third cluster is weakly sub-clustered into samples with adequate MAG detection (Rt5, Rt7, Rt11 and Rt13) and those without (U027, U035, LB003, Blanks, Rt_tooth, Fig. 4). The biplot shows that the separation of modern and ancient samples on PCo1 is chiefly driven by the taxa which are exclusively present in the corresponding samples, while taxa present in both ancient and modern samples are driving the separation of these samples along PCo2 (Fig. 4). Modern samples show weak clustering by sampling location (Rondane Sør and Forollhogna), which becomes more obvious when subsetting the modern samples by themselves (Supp. Fig. 11). The modern samples did not cluster by age at death (Supp. Fig. 11 & 12).

PERMANOVA analyses confirmed that microbial communities of samples of different age classes were significantly different (*r*^2^ = 0.200, p < 0.001), without an effect of the number of microbial reads (*r*^2^ = 0.019, p = 0.184) when modelling these variables together. However, when investigating the effect of these variables independently, both age class and read number were significant and explained more variance (age class: *r*^2^ = 0.431, p < 0.001; microbial reads: *r*^2^ = 0.249, p < 0.001). In modern samples, we detected a significant effect of sampling area (*r*^2^ = 0.156, p < 0.001), a weak but significant effect of individual age at death (*r*^2^ = 0.085, p < 0.05), but no effect of the number of microbial reads (*r*^2^ = 0.059, p = 0.096). This result remained when looking at the variables independently (area: *r*^2^ = 0.168, p < 0.001; age: *r*^2^ = 0.087, p < 0.05; microbial reads: *r*^2^ = 0.068, p = 0.113). The dispersion (average distance to median) was 9.168 for ancient samples, 8.568 for modern samples and 5.525 for historical samples. While modern samples are slightly more clustered than ancient samples, the difference is not significant (Tukey-test, adjusted p = 0.717 [modern / ancient]). Historical samples are weakly, but significantly more clustered than modern samples (Tukey-test, adjusted p < 0.05 [modern / historic]), and significantly more clustered than ancient samples (Tukey-test, adjusted p < 0.05 [ancient / historic]).

The community clustering of reindeer samples is similar when assigning taxonomy with *KrakenUniq.* In the whole community ordinations, samples cluster into distinct ancient and modern clusters, with historical samples occupying the space in between (Supp. Fig. 13A-B). The clustering is somewhat weaker and less pronounced than in MAG ordinations (Fig. 4). Clustering in *KrakenUniq* ordinations becomes more pronounced when focusing on the reindeer oral microbiome and to a lesser extent on the reindeer gut microbiome (Supp. Fig. 13C-E). Overall, more putative oral taxa appear to be at higher abundances in modern samples, while more putative GIT taxa are at higher abundances in ancient samples (Supp. Fig. 14B, D).

### Comparison between MAG-based analysis and read-based analysis

The taxon recovery of the read-based approach with KrakenUniq overall matched the taxa classified with the MAG-based approach using Anvi’o, albeit with few key differences. On average, 42% of reads mapped to bins in the MAG-based approach (range: 11-60%), whereas only 12% of reads were classified with the *KrakenUniq* read-based approach (range: 2-35%; Supp Fig. 15). When comparing the classification by habitat (oral vs. GIT), Kraken tended to capture more putative oral-only microbes, while MAGs tended to capture more GIT-only microbes, particularly in the ancient samples (Supp Fig. 13). However, ancient samples generally had higher abundances of GIT-associated microbes, regardless of classification method (Supp. Fig. 13).

When comparing genus-level microbial community diversity, KrakenUniq yielded an overall greater diversity. Of the 684 unique genera detected with either of the two methods, 618 (90%) were detected only by KrakenUniq, 33 (5%) were detected only by MAGs and 33 (5%) were detected by both methods (Supp. Fig. 16B). However, the 618 genera detected only by KrakenUniq represent merely 16% of all reads classified with KrakenUniq, while the 33 genera only detected by MAGs represent 36% of all reads classified with the MAG-based approach (Supp. Fig. 16B). As expected, KrakenUniq outperforms MAG-based approach in the detection of rare low-abundance taxa. This pattern remains consistent when comparing ancient and modern samples (Supp. Fig. 17A-B). The abundance at which taxa were detected varied between taxa and the detection method, however, most high abundance taxa were detected at similar abundances by both methods (Supp Fig. 17A). At taxonomic levels higher than class, 5.6% of the KrakenUniq assignments were not detected by the MAG approach, whereas 0.52% of the MAG assignments were not present in the KrakenUniq dataset (black point in Supp. Fig. 17A). Of the top 50 highest abundance taxa detected by each method, 32 (64%) were detected by both methods, 26 (52%) were detected only by the MAG-based approach and 21 (42%) were detected only by the short-read approach with KrakenUniq. Of the 20 most abundant species found with *KrakenUniq,* four were also identified at the species level in the MAGs, while 14 were identified at the genus level in the MAGs (Supp. Fig. 17C).

### Taxa overlapping temporal datasets

Among the 115 putatively non-contaminant MAGs, we identified six that were present in all age classes, and therein were present in almost all samples (*Propionibacterium ruminifibrarum*, Rumen uncultured bacterium 99 (*RUG099 sp.*), *Basfia_A sp., Succinclasticum sp.*, *Streptococcus chenjunshii* and *Asp313 sp,* Fig. 3B). However, none of these taxa were detected in four samples (U027, U035, Rt_tooth, Rt1). All of these taxa were first isolated from non-reindeer ruminants, specifically their gastrointestinal tracts or feces. Ancient microbial reads mapping to these MAGs show high degrees of typical ancient DNA damage (Supp. Fig. 9).

To further investigate strain-level differences between ancient and modern communities, differential abundance analysis with Maaslin2 (Mallick et al. 2021) was performed on the MAG dataset. For this analysis, we excluded blanks, historical samples (due to the low sample size) and the three ancient samples with outlying/different communities and low signs of ancient DNA damage (U027, U035 and LB003). The analysis yielded 56 significant taxa (FDR < 0.05), 50 of which are bins present only in either ancient or modern samples but not in both (Supp. Table 2). *Asp313 sp.* is present and about equally abundant in both datasets (FDR = 0.759, coefficient = 0.161, p = 0.759, Fig. 5).

**Figure 5.**
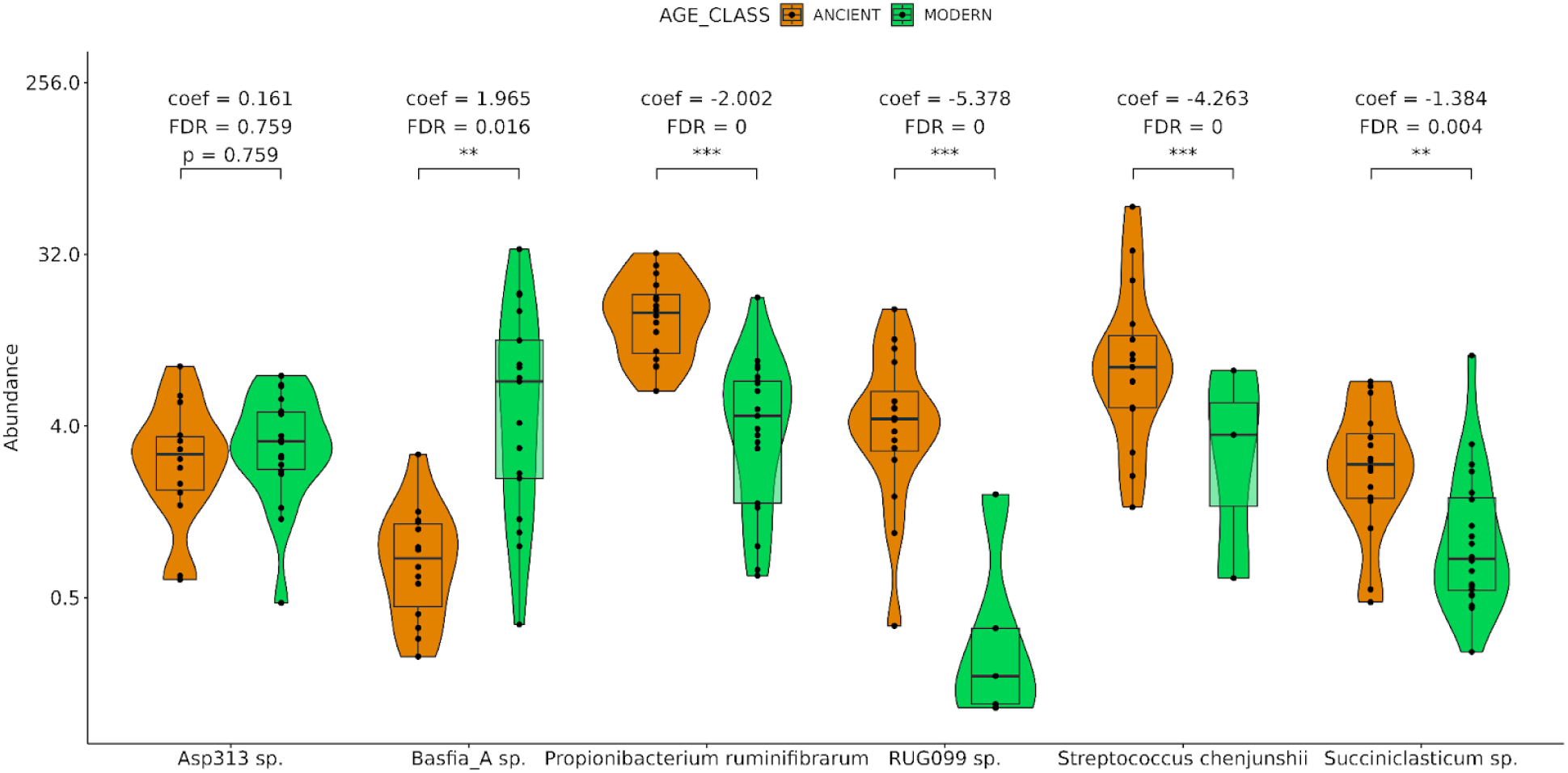
Differential abundance of six selected rumen microbes. Filtered (detection > 0.3) abundance is compared between six taxa present in both ancient (orange) and modern (green) samples. The scale is log2 transformed, which is the default log transformation performed by MaAsLin2 (Mallick et al. 2021). Each point represents one sample. Samples in which the taxon is not present are omitted. Point density at any given abundance value is visualized as a violin plot. Boxes represent the 25th percentiles (lower margin), 50th percentiles (median, thick line) and 75th percentiles (upper margin) of the value distribution. Maaslin2 metrics (coefficient (coef), p-value (p), false discovery rate (FDR)) are noted in brackets on the top. Metrics are rounded to three decimal points (p < 0.01 = **, p < 0.001 = ***).

The other five taxa that are present in both modern and ancient samples have significantly different abundances in the age groups (Fig. 5, Supp. Tab. 2). *Propionibacterium ruminifibrarum* (FDR = 0, coefficient = -2.002, p < 0.001), *RUG099 sp*. (FDR = 0, coefficient = -5.378, p < 0.001), *Succiniclasticum sp*. (FRD = 0.004, coefficient = -1.384, p = 0.01), *S. chenjunshii* (FDR = 0, coefficient = -4.263, p < 0.001) are more abundant in ancient samples, *Basfia_A sp*. (FDR = 0.016, coefficient = 1.965, p < 0.01) is more abundant in modern samples.

Overall, *KrakenUniq* was more successful in detecting taxa which are shared between both the ancient and modern datasets. *KrakenUniq* detected 122 (19% of total) genera in both sets vs. 9 (14%) in the MAG analysis (Supp. Fig. 17C-D). At the species-level, KrakenUniq detected 241 (13% of total) species in both ancient and modern samples (vs. 6 in the MAG analysis; Supp. Fig. 17E-F), of which 19 species (8%) rank among the top 20 most abundant species (vs. 6 out of 6 in the MAG analysis). However, some of the detected species might be bacterial strains of the same species, which we can not clearly distinguish. For taxa that can be matched between the detection methods, results are relatively consistent. Within the 241 species detected by KrakenUniq in both ancient and modern samples, 87 (36%) are GIT-associated, 54 (22%) are oral-associated and 49 (20%) are associated with both habitats.

### Recovery of dietary information from dental calculus

For the investigation of putative dietary items, based on prevalence in blank and museum control samples, twelve genera were identified as potential contaminants and excluded from further analysis. These were as follows: *Callitriche, Capsicum, Cichorium, Citrus, Cucurbita, Dioscorea, Fumaria, Helianthus, Hippuris, Pinus, Triticum* and *Zea*. *Capsicum* and *Citrus* were only detected in blank samples. Thus the final list of putative dietary taxa identified among all samples consisted of 75 plant genera and 1 lichen genus. Only six genera were identified with high confidence in the ancient samples, although many genera were present at low confidence, as there were too few reads to evaluate damage patterns (Fig. 6A). Only one taxon, *Veronica*, was detected at high confidence in both ancient and modern samples; however, it was also observed at low abundance in a museum swab even after contamination filtering, thus we remain cautious in interpreting the dietary significance of this taxon (Fig. 6A-B). Most of the genera identified in the historical samples were also identified in the modern samples, while four taxa were shared between ancient and historical samples (Fig. 6B). No genus was detected with high confidence in all three age classes (Fig. 6B). Within the modern age class, nine genera were detected in both sampling areas (Fig. 6C), with *Salix* and *Carex* detected more frequently in Forollhogna, *Ceraphylium* detected more frequently in Rondane, and *Zannichelia* and the lichen genus *Cladonia* detected similarly in both (Fig. 6A).

**Figure 6.**
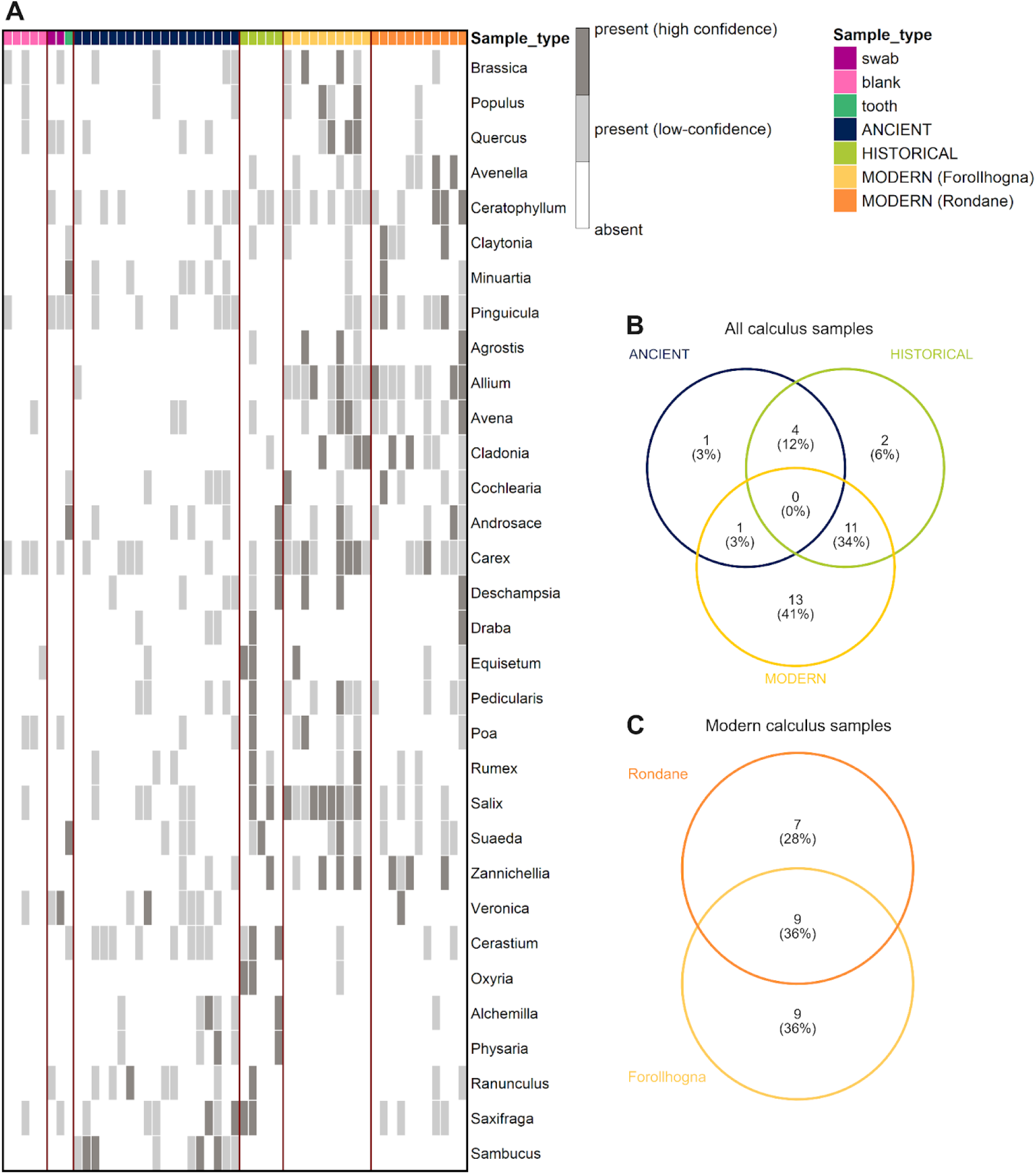
Plant and lichen genera detected among sample age classes. **A.** Heatmap showing the presence or absence of each genus within each sample, for genera detected in at least two samples. Genera detected at low confidence in a sample (<100 reads but ≥10 reads, damage not evaluated) are indicated in light grey. Genera are ordered by shared and unique taxa identified in B. **B**. Venn diagram showing shared and unique high-confidence genera among ancient, historical and modern age classes. **C.** Venn diagram showing shared and unique high-confidence genera between the Rondane and Forollhogna populations within the modern age class. In all plots, genus-level taxa only detected in one sample are excluded.

## Discussion

Here, we characterized the oral microbiome and putative dietary components captured in the dental calculus of contemporary and prehistoric reindeer. We showed that reindeer dental calculus is a suitable source material for the long-term recovery of oral and gut microbiota, as well as plants that are known components of reindeer diets. While we observe a turnover in microbial community membership and composition between archaeological and fresh material, parts of the gut microbiome appear to be remarkably stable throughout time and space. Specifically, we assembled six partial genomes of such microbial taxa, all with putative roles in the reindeer gastrointestinal tract.

### Dental calculus can be used to identify modern and ancient rumen microbes

Although we detected major differences between sample age classes, six MAGs (*Propionibacterium ruminifibrarum,* Rumen uncultured bacterium 099 *(RUG099 sp.), Basfia_A sp., Succinclasticum sp., Streptococcus chenjunshii* and *Asp313 sp.*) were identified in a majority of samples from both groups. All six taxa are associated with the ruminant gastrointestinal tract. Since they have been previously found in other ruminant host species, it is likely that these taxa, or closely related strains thereof, have a conserved function in the reindeer. Since these taxa were found in both ancient and modern calculus, we suggest that these six microbes are true members of the reindeer oral and/or rumen microbiome. The fact that these are the only taxa we found in common across time and space suggests that they might be important, highly conserved microbiome members that are also likely highly abundant in the rumen. However, preservation bias might play a role (see below). A future comparative pangenomic analysis of our study MAGs with strains from other ruminants might confirm our hypotheses.

The six taxa of interest are variably well understood, but each taxon or a closely related strain has been isolated before from ruminant gastrointestinal tracts or connected systems such as feces. *Propionibacterium ruminifibrarum (Vaidya et al. 2019)* is a described, cultured, gram-positive species which was first isolated from the rumen of a dairy-cow. In the rumen, *P. ruminifibrarum* ferments propionate and a range of other carbohydrates and amino acids from plant-fiber material, which are useful substrates for the host (Vaidya et al. 2019). MAGs of rumen uncultured bacterium 099 *(RUG099 sp.* from the order Peptostreptococcales) have been assembled from cow rumen sequences (Stewart et al. 2018)*. Basfia_A sp.* is a member of the family Pasteurellaceae. *Basfia* and its type species *Basfia succiniproducens* were first isolated and described from cow rumina, where they anaerobically produce succinic acid from glucose (Kuhnert et al. 2010). The type species of genus *Succinclasticum* is *Succiniclasticum ruminis* (van Gylswyk 1995), was first isolated from cow rumina where it converts succinate to propionate. Other unclassified species or strains in this genus have been found in cattle (Gharechahi et al. 2021) as well as other ruminants, including reindeer rumen (Glendinning et al. 2021). *Streptococcus chenjunshii (Tian et al. 2019)* was first isolated from feces of Tibetan antelope (*Pantholops hodgsonii*), which suggests an association with the gut microbiome. Tibetan antelope is a bovid from the alpine grasslands of Tibet (Wei et al. 2023), not unlike the tundra habitat inhabited by the reindeer. Tibetan antelope nematode gut parasites *Trichostrongylus*, *Marshallagia* and *Nematodirus spp.* have also been found in Svalbard reindeer (*R. t. platyrhynchus*) (Stien et al. 2002; Cao et al. 2022), suggesting similarities between the gastrointestinal environments of these two hosts. *Asp313 sp*. is a member of the family Actinomycetaceae which has been isolated from feces of tibetan antelope (BioSample: SAMN08967870) and goat (*Capra hircus*) (BioSample: SAMN11294320, (Peng et al. 2021)).

GIT-associated taxa were more frequently shared between ancient and modern samples, compared to oral taxa, using either the MAG-based or *KrakenUniq* approach. GIT-associated taxa were also more abundant in ancient samples than in modern samples independent of the approach used. One possible explanation for this pattern is that since the oral microbiome is more directly interacting with the environment of the host, it is more strongly affected by changes in the environment, such as changes in diet, climate, or increase in human-made antibiotics in the environment (Brealey et al. 2021; Moraitou et al. 2022). In contrast, gut or rumen microbes may be more conserved due to their important role in gut function in ruminants.

Rumen-associated prokaryotic and eukaryotic microbes were also identified in the original analysis of the historical samples by (Brealey et al. 2020), which did not utilize MAGs. However, the authors demonstrated that authentic MAGs can be recovered from non-model animal dental calculus and suggested the more extensive use of MAGs to improve the future analysis of ancient reindeer dental calculus. Here, we were successful in recovering reindeer oral and rumen bacteria by assembling MAGs from a wider variety of samples, further revealing the utility of metagenome-assembled genomes. None of the six rumen microbes shared between ancient and modern samples have been previously reported as members of reindeer microbiomes (Sundset et al. 2009a; Pope et al. 2012; Salgado-Flores et al. 2016; Zielińska et al. 2016; Kamenova et al. 2023). Conversely, taxa in the family *Ruminococcaceae*, which were reported as highly abundant in previous studies (Pope et al. 2012; Salgado-Flores et al. 2016; Zielińska et al. 2016), are completely absent from our MAG assembly results (see Supp. Tab. 1) and only present at very low abundances in the *KrakenUniq* results. One important reason for this discrepancy may be that most of these previous studies did not employ whole-genome shotgun sequences as done here, but instead used 16S rRNA amplicon sequencing. To date, MAGs have only been assembled from the rumen, small intestine and colon of reindeer inhabiting the high-Arctic Svalbard archipelago (Kamenova et al. 2023), a population exhibiting strong genetic and dietary divergence (Dussex et al. 2025). We did not detect any of those 18 species-level MAGs among our species-level MAGs. Furthermore, only the genera *Bacteroides*, *Prevotella*, and *Methanobrevibacter* could be identified from MAGs in both our study and in (Kamenova et al. 2023). *KrakenUniq* detected 14 genera at low abundances which were also detected in (Kamenova et al. 2023). Both shotgun metagenomics and 16S rRNA can be biased toward detecting certain taxa in ancient metagenomic datasets (Ziesemer et al. 2015; Warinner et al. 2017; Velsko et al. 2018), thus neither perfectly detect the entire microbiome. Furthermore, many of the previous studies were based on feces, food matter that passed through multiple stomachs and sequentially altered by each local microbiome, making it a less accurate proxy for the rumen microbiome (Tapio et al. 2016). On the other hand, rumen content is directly regurgitated into the mouth.

### Diversity in reindeer oral microbiome and putative dietary profiles

We observed clear differences in the composition of oral and GIT microbes, and to a lesser extent, plants and lichens, both between ancient and modern reindeer dental calculus and between the two modern reindeer populations. The technical biases discussed above cannot explain all the differences observed between the modern populations, which instead likely have ecological and evolutionary explanations. The Forollhogna population is descended from domesticated reindeer, while the southern Rondane population has experienced little introgression with domestic populations (Kvie et al. 2019). These genetic differences could influence microbial exposure via herd dynamics and feeding behaviour, and microbial colonization via immune processes (Ezenwa et al. 2012; Ryu and Davenport 2022), leading to differing oral microbiome compositions. Since the populations are geographically separated, differences in the microbiome and putative dietary profiles could also be driven by habitat. Forollhogna is the most productive wild reindeer area in Norway. Unlike the majority of the other wild reindeer areas, Forollhogna is mostly located in the subalpine region. The area is characterized by vegetation types with grass heaths, heather and willow providing important forage resources during summer and snow free periods (Kjørstad et al. 2017). Forollhogna also possess abundant lichen resources which generally represent an important winter forage for reindeer (Kjørstad et al. 2017). Regarding forage availability, the southern Rondane region shares many of the qualities and characteristics of Forollhogna although the landscape is more heterogeneous and with a larger proportion of alpine areas. Various species of herbaceous plants and grasses were observed in both populations, including alpine and tundra genera like *Androsace* and *Cochlearia*, as well as the lichen *Cladonia*, which is one of the primary winter foods of present-day reindeer populations, excepting the reindeer of Svalbard (Staaland et al. 1988; Nieminen and Heiskari 1989). We found slightly more plant taxa in the Forollhogna population compared to Rondane, with increased detection of the willow genus *Salix* and the sedge *Carex*. Diet can directly affect gut microbiome composition (Storeheier et al. 2002; Salgado-Flores et al. 2016; Kamenova et al. 2025), and it is possible that diet may affect oral microbiome composition (Moraitou et al. 2022). Together, our results suggest that ecological factors play a role in shaping the reindeer oral microbiome (Brealey et al. 2021; Moraitou et al. 2022). Because we are able to partly explain some of the differences between two contemporaneous populations based on their ecology, it might be that some of the differences between the modern populations and ancient populations are at least partly due to ecological factors.

There are intra-population differences between samples from different individual reindeer, which may be due to individual variation in genetics, physiology, behaviour, feeding ecology, or individual age at death. Since dental calculus forms continuously over the entire lifetime of an individual, its microbial community is also individual- and age-dependent (Utter et al. 2016). However, as we were not able to identify the age at death of the ancient individuals, we could not investigate whether the differences between ancient and modern samples were due to the effect of age. We do not know exactly how the ancient reindeer in this study were selected for killing and transport from the kill site to the site of consumption by their prehistoric human hunters. However, research on prehistoric humans in France (Boyle 1993; Kuntz and Costamagno 2011) and Neanderthals in Germany (Gaudzinski and Roebroeks 2000) and France (Niven et al. 2012) indicates that adult reindeer were preferentially selected for hunting. The modern samples, for which we know the age at death, were all adults (≥ 2 years of age) and did not cluster according to their age at death in the microbial ordinations, suggesting that adult age played no major role in the differences we observed.

### Eco-evolutionary dynamics as possible explanations for spatio-temporal differences in the reindeer metagenome

We hypothesize that the large differences in microbial and plant-taxon composition in dental calculus among the ancient and modern samples could possibly be explained by the following ecological and evolutionary factors or their interactions: 1) The local and temporal availability of vegetation to feed on and 2) plasticity in reindeer foraging behavior. It has been shown that variation in the gut microbiome in reindeer (Salgado-Flores et al. 2016) and oral microbiome in other mammals (Moraitou et al. 2022) is driven by ecological factors, specifically diet. The oral cavity is a point of entry for microbes, through interactions with microbes in plants, soil and water during feeding, and thus the microbiome preserved in dental calculus may more strongly reflect changes in the local environment than other host-associated microbiomes (Brealey et al. 2021; Moraitou et al. 2022). The role of the environment for determining diet availability is indicated by the presence of certain plants such as *Salix ssp.* in our samples. *Salix ssp.* represents high quality forage linked to higher autumn body mass, a key survival factor, in reindeer (Kamenova et al. 2025). We detected *Salix ssp.* with high confidence in historical samples and in the modern Forolhogna population, but with only low confidence in ancient samples and the modern Rondane population. Our pattern of detection concurs that the consumption of this plant is subject to local availability, which in turn is influenced by annual climate variation as well as long-term climatic trends such as arctic greening (Kamenova et al. 2025).

While many plant genera were detected in modern and historical samples, only six genera were detected in ancient samples, the majority of which are herbaceous. We found no evidence of reindeer specialising on certain plant taxa, even in the modern populations. Thus, despite our small dataset, our results generally agree with previous studies suggesting that late Pleistocene reindeer from south France were mixed feeders with high levels of plasticity in dietary preference (Merceron and Madelaine 2008; Drucker et al. 2011; Rivals and Semprebon 2017; Rivals et al. 2020; Britton et al. 2023). Dietary plasticity has been suggested as a contributing factor in the species continued survival through changing climate at the end of the last glacial period (Rivals et al. 2020). Furthermore, there are documented examples of reindeer adapting their diet to local conditions from South Georgia Island (Mathiesen et al. 1999) and Svalbard (Hansen et al. 2019).

We hypothesize that differing environmental conditions in the late Pleistocene could be contributing to the differing dietary and oral microbiome profiles that we observed in the ancient French reindeer in comparison to the modern Norwegian populations. However, rumen-derived microbes that remained stable over this long time period may have conserved functions within the host. This combination of dietary plasticity with potentially conserved rumen microbes may have contributed to reindeer adaptation to the changing vegetation associated with climatic change and range shifts across the last glacial maximum and the transition to the Holocene. Investigating this hypothesis would require additional research into the microbial and dietary profiles of prehistoric and contemporary reindeer. We note that the strong turnover we observe in the oral microbiome of reindeer is quite unlike the rather high evolutionary stability reported in studies of the hominid oral microbiome (Fellows Yates et al. 2021). Thus, the transfer of insights between species may be limited due to their intrinsic characteristics. Further comparative research, which is presently outside the scope of this study, is needed to elucidate the factors at play.

### Technical challenges for ancient metagenomics

There are several important challenges to overcome when working with ancient metagenomics, including the identification and removal of external contamination and controlling for potential biases introduced by differences in sample preservation levels (Salter et al. 2014; Mann et al. 2018; Eisenhofer et al. 2019). Dietary reconstruction from ancient metagenomic data presents a further challenge due to incomplete reference databases and the potential for microbial DNA to dominate the datasets and obscure the proportionally smaller number of dietary DNA sequences (Mann et al. 2023). We used multiple techniques to address these issues, including utilization of blank samples and museum controls, assessment of damage patterns, microbial source-tracking and controls for sequencing depth. Furthermore, sequences in non-microbial reference databases might be heavily contaminated with microbial reads, such as was the case with the PhyloNorway database we used in this study (Oskolkov et al. 2025). However, it is difficult or impossible to distinguish microbial contamination in a database from actual plant-associated microbes, such as endosymbionts. Therefore, decontamination measures might remove valid targets as false-positives, erroneously lowering the detection rate when using such a database.

We used microbial source-tracking to explore putative sources of contaminating and endogenous microbes in our dental calculus samples. Human skin microbes as a source accounted for only a small fraction of reads in the reindeer samples, providing further confidence that our strategies to minimize contamination were successful. Ancient samples contained a larger fraction of reads sourced from soil microbes, likely reflecting their thousands of years in soil before being excavated, whereas modern samples contained higher proportions of oral microbiome sources, possibly due to these samples’ freshness. However, the majority of reindeer dental calculus sequences were assigned an ‘unknown’ source. This large fraction presumably includes residual DNA from the reindeer host, as well as DNA from reindeer-specific oral and gut microbes and plant sources, which the source-tracking database, originally built for human microbiome data, did not contain (see also (Moraitou et al. 2025)). Biases and limitations from human-centric reference databases are a common challenge in microbiome research in non-model animals, which highlights the need for expanded sequencing efforts of underrepresented organisms (Brealey et al. 2020; Mann et al. 2023).

Differences in levels of preservation and *post-mortem* decay can introduce additional biases into ancient metagenomic analyses (Ziesemer et al. 2015; Mann et al. 2018). Some evidence suggests that microbial species with higher GC content (and therefore higher DNA stability) could be better preserved over time (Mann et al. 2018), and the same may be true for eukaryotic DNA. While we did observe slightly higher GC content in MAGs assembled from ancient and historical samples compared to modern samples, the difference was not statistically significant. *Post-mortem* decay also results in short DNA fragments, complicating taxonomic assignment, particularly for eukaryote DNA (Mann et al. 2023). In ancient metagenomes, taxa with smaller genomes may be more accurately assigned to lower taxonomic levels than taxa with large, highly repetitive genomes (Mann et al. 2023), whereas the longer DNA fragments in modern samples may reduce this problem. Finally, *post-mortem* decay can lead to overall lower DNA concentrations in ancient samples, reducing the possibility to identify rare taxa and thus artificially lowering estimates of alpha diversity. In our datasets, sequencing depth (per-sample number of reads matching a particular target) generally explained only around 3% of the variation among samples in the microbial ordinations, much less than age class (ca. 15-20%). Nevertheless, preservation biases may be driving some differences in community compositions between ancient and modern samples, particularly for taxa that were detected primarily in the ancient samples. We therefore remain cautious in our interpretations of the microbes and putative dietary taxa detected only in the ancient reindeer.

In addition to our main MAG-based analysis, we conducted a read-based analysis with *KrakenUniq* to assess possible methodological biases influencing the microbial results of the study. Overall, the MAG-based approach was able to classify a slightly higher proportion of sequences than *KrakenUniq*. On the microbial community level, *KrakenUniq* performed better at identifying rare taxa at low abundances. In terms of host body habitats, MAGs seemed to favor the capture of gut-only microbes, while *KrakenUniq* favored the capture of oral-only microbes. Abundance estimates were heavily dependent on the taxon, therefore no clear methodological comparison can be drawn. When comparing the whole microbial communities of individual reindeer, the community patterns are consistent: samples cluster by age class and source population, regardless of the metagenomic approach. The clustering changes slightly when focusing only on putative oral or GIT microbiomes. Some oral-only microbes are found at higher abundances in modern samples, while many GIT-only microbes are found at higher abundance in ancient samples.

### Conclusion and future directions

Here, we conducted a comparative study of modern and ancient dental calculus of reindeer. With advanced techniques in ancient DNA sequencing and metagenome assembly, it was possible to retrieve microbial genomes and putative dietary profiles from samples surpassing 20 ka BP in age. Our work shows the potential for utilizing dental calculus to not only study temporal changes in the ecology and evolution of the oral microbiome, but also of the ruminant gastrointestinal tract. Our study also suggests that certain rumen-associated taxa have been more temporally stable than the oral microbiome.

This study sets the stage for future research on the ecology and evolution of reindeer as a holobiont (Bordenstein et al. 2024). Future studies could dramatically expand both spatial and temporal range and density of sampling. *Rangifer tarandus* is subdivided into numerous subspecies which occupy varying habitats and dispersal ranges throughout the northern

Holarctic. We hypothesize that various reindeer subspecies could differ in the composition of their microbiomes, as has been observed in other mammalian species (Moraitou et al. 2022). By sampling reindeer dental calculus from ancient samples from additional former habitats and time frames, it will be possible to trace the evolution of the microbiome through time and space and link it to past reindeer range expansions and contractions caused by ice sheet retractions and expansions. However, better understanding of the contemporary reindeer microbiome is needed to give context to past microbial changes. Research on the microbiomes of reindeer and other ruminants could therefore be further advanced by an integrative approach combining the sampling of dental calculus, buccal swabs, direct gut sampling, bolus and feces. Such approaches would further our understanding of colonization dynamics among the microbial niches of the gut, rumen and oral cavity.

We also demonstrated that dental calculus contains plant DNA from putative food plants of reindeer. More extensive sampling efforts for metagenomics analysis could be used to better understand reindeer diet on a spatiotemporal level throughout the Pleistocene and might even be suitable as a proxy for the distribution of certain plant species (Brealey et al. 2020). Future studies could complement records of past vegetation distributions (e.g. from the pollen record) with ancient dental calculus metagenomic data to further elucidate how reindeer and other, now extinct, mammoth-steppe megafauna have adapted their foraging behavior to changes in vegetation cover, as well as the parallel effects on their microbiomes. In turn, this knowledge could help us understand and predict how arctic herbivores might react to environmental changes due to the pressing matter of future anthropogenic climate change.

## Acknowledgements

This work was funded by Norwegian Research Council award number 325589 to MDM and the Swedish Research Council (Formas) grant number 2019-00275 to KG. We thank Adrian Forsythe, Jørgen Rosvold and Laurène Lecaudey for assistance with sample collection. We thank Clara Tatalidis for assistance with modern sample wet lab work. We are grateful to Stefaniya Kamenova for a productive early discussion of reindeer dietary sources. Furthermore, we thank the staff of the Musée National de Préhistoire (Les Eyzies, France), especially Stéphane Madeleine and Jean-Jacques Cleyet-Merle for assisting with sample acquisition and providing the opportunity to conduct research on these valuable materials. Finally, we thank Reidar Andersen and Hans Stenøien for providing inspiration and logistical support to pursue this work.

## Author Contributions

Writing: FLK, JCB

Editing: all authors

Data analysis microbiome: FLK, JCB

Data analysis plants: JCB, MWP, NV

Radiocarbon dating: BP, MS

Modern lab work & sampling: SLFM, KG, VV

Ancient lab work & sampling: FLK, SLFM

Interpretation of results: all authors

Funding: MDM, KG

Conception: FLK, MDM

Development: FLK, JCB, MDM

Supervision: MDM, VCB, MWP

## Data availability statement

Raw sequence data of modern and ancient samples are archived on the European Nucleotide Archive (ENA) under project accession code PRJEB106305. Raw sequence data of historic samples is archived in the ENA under accession number PRJEB33363. Sample metadata is available in Table 1. Additional information is available in the supplementary material.

## References

Alneberg J, Bjarnason BS, de Bruijn I, Schirmer M, Quick J, Ijaz UZ, Lahti L, Loman NJ, Andersson AF, Quince C. 2014. Binning metagenomic contigs by coverage and composition. Nat. Methods [Internet] 11:1144–1146. Available from: 10.1038/nmeth.3103

Amin N, Schwarzkopf S, Kinoshita A, Tröscher-Mußotter J, Dänicke S, Camarinha-Silva A, Huber K, Frahm J, Seifert J. 2021. Evolution of rumen and oral microbiota in calves is influenced by age and time of weaning. Anim Microbiome [Internet] 3:31. Available from: 10.1186/s42523-021-00095-3

Bordenstein SR, The Holobiont Biology Network, Holobiont Biology Network. 2024. The disciplinary matrix of holobiont biology. Science [Internet] 386:731–732. Available from: https://www.science.org/doi/10.1126/science.ado2152

Borry M, Hübner A, Rohrlach AB, Warinner C. 2021. PyDamage: automated ancient damage identification and estimation for contigs in ancient DNA de novo assembly. PeerJ [Internet] 9:e11845. Available from: 10.7717/peerj.11845

Boyle KV. 1993. Upper Palaeolithic procurement and processing strategies in southwest France. Archeol. Pap. Am. Anthropol. Assoc. [Internet] 4:151–162. Available from: https://anthrosource.onlinelibrary.wiley.com/doi/10.1525/ap3a.1993.4.1.151

Brealey JC, Leitão HG, Hofstede T, Kalthoff DC, Guschanski K. 2021. The oral microbiota of wild bears in Sweden reflects the history of antibiotic use by humans. Curr. Biol. [Internet] 31:4650–4658.e6. Available from: 10.1016/j.cub.2021.08.010

Brealey JC, Leitão HG, van der Valk T, Xu W, Bougiouri K, Dalén L, Guschanski K. 2020. Dental Calculus as a Tool to Study the Evolution of the Mammalian Oral Microbiome. Mol. Biol. Evol. [Internet] 37:3003–3022. Available from: 10.1093/molbev/msaa135

Breitwieser FP, Baker DN, Salzberg SL. 2018. KrakenUniq: confident and fast metagenomics classification using unique k-mer counts. Genome Biol. [Internet] 19:198. Available from: 10.1186/s13059-018-1568-0

Britton K, Jimenez E-L, Le Corre M, Renou S, Rendu W, Richards MP, Hublin J-J, Soressi M. 2023. Multi-isotope analysis of bone collagen of Late Pleistocene ungulates reveals niche partitioning and behavioural plasticity of reindeer during MIS 3. Sci. Rep. [Internet] 13:15722. Available from: 10.1038/s41598-023-42199-7

Bronk Ramsey C. 2009. Bayesian analysis of radiocarbon dates. Radiocarbon [Internet] 51:337–360. Available from: 10.2458/azu_js_rc.51.3494

Cao Y, Foggin M, Zhao X. 2022. Tibetan antelope migration during mass calving as parasite avoidance strategy. Innovation (Camb) [Internet] 3:100326. Available from: 10.1016/j.xinn.2022.100326

Chaumeil P-A, Mussig AJ, Hugenholtz P, Parks DH. 2019. GTDB-Tk: a toolkit to classify genomes with the Genome Taxonomy Database. Bioinformatics [Internet] 36:1925–1927. Available from: 10.1093/bioinformatics/btz848

Chaumeil P-A, Mussig AJ, Hugenholtz P, Parks DH. 2022. GTDB-Tk v2: memory friendly classification with the genome taxonomy database. Bioinformatics [Internet] 38:5315–5316. Available from: 10.1093/bioinformatics/btac672

Chen T, Yu WH, Izard J, Baranova OV, Lakshmanan A, Dewhirst FE. 2010. The Human Oral Microbiome Database: a web accessible resource for investigating oral microbe taxonomic and genomic information. Database [Internet]. Available from: 10.1093/database/baq013

Clark PU, Dyke AS, Shakun JD, Carlson AE, Clark J, Wohlfarth B, Mitrovica JX, Hostetler SW, McCabe AM. 2009. The Last Glacial Maximum. Science [Internet] 325:710–714. Available from: 10.1126/science.1172873

Dabney J, Knapp M, Glocke I, Gansauge M-T, Weihmann A, Nickel B, Valdiosera C, García N, Pääbo S, Arsuaga J-L, et al. 2013. Complete mitochondrial genome sequence of a Middle Pleistocene cave bear reconstructed from ultrashort DNA fragments. Proc. Natl. Acad. Sci. U. S. A. [Internet] 110:15758–15763. Available from: 10.1073/pnas.1314445110

Davis NM, Proctor DM, Holmes SP, Relman DA, Callahan BJ. 2018. Simple statistical identification and removal of contaminant sequences in marker-gene and metagenomics data. Microbiome [Internet] 6:226. Available from: 10.1186/s40168-018-0605-2

Drucker DG, Kind CJ, Stephan E. 2011. Chronological and ecological information on Late-glacial and early Holocene reindeer from northwest Europe using radiocarbon (14C) and stable isotope (13C, 15N) analysis of bone collagen: Case study in southwestern Germany. Quat. Int. [Internet] 245:218–224. Available from: 10.1016/j.quaint.2011.05.007

Duitama González C, Vicedomini R, Lemane T, Rascovan N, Richard H, Chikhi R. 2023. decOM: similarity-based microbial source tracking of ancient oral samples using k-mer-based methods. Microbiome [Internet] 11:243. Available from: 10.1186/s40168-023-01670-3

Dussex N, Bieker VC, Sun X, Le Moullec M, Ersmark E, Røed KH, Speakman JR, Loe LE, Dalén L, Hansen BB, et al. 2025. The genomic basis of the Svalbard reindeer’s adaptation to an extreme Arctic environment. Genome Biol. Evol. [Internet]. Available from: 10.1093/gbe/evaf160

Eisenhofer R, Minich JJ, Marotz C, Cooper A, Knight R, Weyrich LS. 2019. Contamination in Low Microbial Biomass Microbiome Studies: Issues and Recommendations. Trends Microbiol. [Internet] 27:105–117. Available from: 10.1016/j.tim.2018.11.003

Eren AM, Kiefl E, Shaiber A, Veseli I, Miller SE, Schechter MS, Fink I, Pan JN, Yousef M, Fogarty EC, et al. 2021. Community-led, integrated, reproducible multi-omics with anvi’o. Nat Microbiol [Internet] 6:3–6. Available from: 10.1038/s41564-020-00834-3

Ewels PA, Peltzer A, Fillinger S, Patel H, Alneberg J, Wilm A, Garcia MU, Di Tommaso P, Nahnsen S. 2020. The nf-core framework for community-curated bioinformatics pipelines. Nat. Biotechnol. [Internet] 38:276–278. Available from: 10.1038/s41587-020-0439-x

Ezenwa VO, Gerardo NM, Inouye DW, Medina M, Xavier JB. 2012. Microbiology. Animal behavior and the microbiome. Science [Internet] 338:198–199. Available from: https://www.science.org/doi/10.1126/science.1227412

Fellows Yates JA, Velsko IM, Aron F, Posth C, Hofman CA, Austin RM, Parker CE, Mann AE, Nägele K, Arthur KW, et al. 2021. The evolution and changing ecology of the African hominid oral microbiome. Proceedings of the National Academy of Sciences [Internet] 118:e2021655118. Available from: https://www.pnas.org/doi/abs/10.1073/pnas.2021655118

Fernandez-Guerra A, Borrel G, Delmont TO, Elberling B, Eren AM, Gribaldo S, Jochheim A, Henriksen RA, Hinrichs K-U, Korneliussen TS, et al. 2023. A 2-million-year-old microbial and viral communities from the Kap København Formation in North Greenland. bioRxiv [Internet]:2023.06.10.544454. Available from: https://www.biorxiv.org/content/10.1101/2023.06.10.544454v2.abstract

Feuerborn TR, Palkopoulou E, van der Valk T, von Seth J, Munters AR, Pečnerová P, Dehasque M, Ureña I, Ersmark E, Lagerholm VK, et al. 2020. Competitive mapping allows for the identification and exclusion of human DNA contamination in ancient faunal genomic datasets. BMC Genomics [Internet] 21:844. Available from: 10.1186/s12864-020-07229-y

Flagstad Ø, Røed KH. 2003. Refugial origins of reindeer (Rangifer tarandus L.) inferred from mitochondrial DNA sequences. Evolution [Internet] 57:658–670. Available from: 10.1111/j.0014-3820.2003.tb01557.x

Gancz AS, Farrer AG, Nixon MP, Wright S, Arriola L, Adler C, Davenport ER, Gully N, Cooper A, Britton K, et al. 2023. Ancient dental calculus reveals oral microbiome shifts associated with lifestyle and disease in Great Britain. Nat. Microbiol. [Internet] 8:2315–2325. Available from: 10.1038/s41564-023-01527-3

Gao C-H. 2025. ggVennDiagram: A “ggplot2” implement of Venn Diagram. Github Available from: https://github.com/gaospecial/ggVennDiagram

Gaudzinski S, Roebroeks W. 2000. Adults only. Reindeer hunting at the middle palaeolithic site salzgitter lebenstedt, northern Germany. J. Hum. Evol. [Internet] 38:497–521. Available from: 10.1006/jhev.1999.0359

Gharechahi J, Vahidi MF, Bahram M, Han J-L, Ding X-Z, Salekdeh GH. 2021. Metagenomic analysis reveals a dynamic microbiome with diversified adaptive functions to utilize high lignocellulosic forages in the cattle rumen. ISME J. [Internet] 15:1108–1120. Available from: 10.1038/s41396-020-00837-2

Glendinning L, Genç B, Wallace RJ, Watson M. 2021. Metagenomic analysis of the cow, sheep, reindeer and red deer rumen. Sci. Rep. [Internet] 11:1990. Available from: 10.1038/s41598-021-81668-9

Granehäll L, Huang KD, Tett A, Manghi P, Paladin A, O’Sullivan N, Rota-Stabelli O, Segata N, Zink A, Maixner F. 2021. Metagenomic analysis of ancient dental calculus reveals unexplored diversity of oral archaeal Methanobrevibacter. Microbiome [Internet] 9:197. Available from: 10.1186/s40168-021-01132-8

van Gylswyk NO. 1995. Succiniclasticum ruminis gen. nov., sp. nov., a ruminal bacterium converting succinate to propionate as the sole energy-yielding mechanism. Int. J. Syst. Bacteriol. [Internet] 45:297–300. Available from: https://www.microbiologyresearch.org/content/journal/ijsem/10.1099/00207713-45-2-297

Hansen BB, Lorentzen JR, Welker JM, Varpe Ø, Aanes R, Beumer LT, Pedersen ÅØ. 2019. Reindeer turning maritime: Ice-locked tundra triggers changes in dietary niche utilization. Ecosphere [Internet] 10:e02672. Available from: https://onlinelibrary.wiley.com/doi/abs/10.1002/ecs2.2672

Hofreiter M, Stewart J. 2009. Ecological change, range fluctuations and population dynamics during the Pleistocene. Curr. Biol. [Internet] 19:R584–R594. Available from: https://www.sciencedirect.com/science/article/pii/S0960982209013062?via%3Dihub

Hold K, Lord E, Brealey JC, Le Moullec M, Bieker VC, Ellegaard MR, Rasmussen JA, Kellner FL, Guschanski K, Yannic G, et al. 2024. Ancient reindeer mitogenomes reveal island-hopping colonisation of the Arctic archipelagos. Sci. Rep. [Internet] 14:4143. Available from: 10.1038/s41598-024-54296-2

Hyatt D, Chen G-L, Locascio PF, Land ML, Larimer FW, Hauser LJ. 2010. Prodigal: prokaryotic gene recognition and translation initiation site identification. BMC Bioinformatics [Internet] 11:119. Available from: 10.1186/1471-2105-11-119

Jin Y, Yip H-K. 2002. Supragingival calculus: formation and control. Crit. Rev. Oral Biol. Med. [Internet] 13:426–441. Available from: 10.1177/154411130201300506

Jónsson H, Ginolhac A, Schubert M, Johnson PLF, Orlando L. 2013. mapDamage2.0: fast approximate Bayesian estimates of ancient DNA damage parameters. Bioinformatics [Internet] 29:1682–1684. Available from: 10.1093/bioinformatics/btt193

Kamenova S, Albon SD, Loe LE, Irvine RJ, Langvatn R, Gusarova G, de Muinck EJ, Trosvik P. 2025. Arctic greening drives changes in the diet and gut microbiome of a large herbivore with consequences for body mass. Ecol. Evol. [Internet] 15:e71731. Available from: 10.1002/ece3.71731

Kamenova S, de Muinck EJ, Veiberg V, Utsi TA, Steyaert SMJG, Albon SD, Loe LE, Trosvik P. 2023. Gut microbiome biogeography in reindeer supersedes millennia of ecological and evolutionary separation. FEMS Microbiol. Ecol. [Internet] 99. Available from: 10.1093/femsec/fiad157

Kanehisa M, Goto S. 2000. KEGG: kyoto encyclopedia of genes and genomes. Nucleic Acids Res. [Internet] 28:27–30. Available from: 10.1093/nar/28.1.27

Kang D, Froula J, Egan R, Wang Z. 2014. MetaBAT: Metagenome Binning based on Abundance and Tetranucleotide frequence. Available from: https://escholarship.org/uc/item/1wz6h4fc

Kapp JD, Green RE, Shapiro B. 2021. A Fast and Efficient Single-stranded Genomic Library Preparation Method Optimized for Ancient DNA. J. Hered. [Internet] 112:241–249. Available from: 10.1093/jhered/esab012

Key FM, Posth C, Krause J, Herbig A, Bos KI. 2017. Mining Metagenomic Data Sets for Ancient DNA: Recommended Protocols for Authentication. Trends Genet. [Internet] 33:508–520. Available from: 10.1016/j.tig.2017.05.005

Kircher M, Sawyer S, Meyer M. 2012. Double indexing overcomes inaccuracies in multiplex sequencing on the Illumina platform. Nucleic Acids Res. [Internet] 40:e3. Available from: 10.1093/nar/gkr771

Kittelmann S, Kirk MR, Jonker A, McCulloch A, Janssen PH. 2015. Buccal swabbing as a noninvasive method to determine bacterial, archaeal, and eukaryotic microbial community structures in the rumen. Appl. Environ. Microbiol. [Internet] 81:7470–7483. Available from: 10.1128/AEM.02385-15

Kjørstad M, Bøthun SW, Gundersen V, Holand Ø, Madslien K, Mysterud A, Myren IN, Punsvik T, Røed KH, Strand O, et al. 2017. Miljøkvalitetsnorm for villrein. Forslag fra en ekspertgruppe. Norsk institutt for naturforskning (NINA) Available from: http://hdl.handle.net/11250/2471598

Klapper M, Hübner A, Ibrahim A, Wasmuth I, Borry M, Haensch VG, Zhang S, Al-Jammal WK, Suma H, Fellows Yates JA, et al. 2023. Natural products from reconstructed bacterial genomes of the Middle and Upper Paleolithic. Science [Internet] 380:619–624. Available from: 10.1126/science.adf5300

Kolde R. 2019. pheatmap: Pretty Heatmaps. url: https://CRAN.R-project.org/package=pheatmap.

Krakau S, Straub D, Gourlé H, Gabernet G, Nahnsen S. 2022. nf-core/mag: a best-practice pipeline for metagenome hybrid assembly and binning. NAR Genom Bioinform [Internet] 4:lqac007. Available from: 10.1093/nargab/lqac007

Kuhnert P, Scholten E, Haefner S, Mayor D, Frey J. 2010. Basfia succiniciproducens gen. nov., sp. nov., a new member of the family Pasteurellaceae isolated from bovine rumen. Int. J. Syst. Evol. Microbiol. [Internet] 60:44–50. Available from: 10.1099/ijs.0.011809-0

Kuntz D, Costamagno S. 2011. Relationships between reindeer and man in southwestern France during the Magdalenian. Quat. Int. [Internet] 238:12–24. Available from: 10.1016/j.quaint.2010.10.023

Kvie KS, Heggenes J, Bårdsen B-J, Røed KH. 2019. Recent large-scale landscape changes, genetic drift and reintroductions characterize the genetic structure of Norwegian wild reindeer. Conserv. Genet. [Internet] 20:1405–1419. Available from: https://link.springer.com/article/10.1007/s10592-019-01225-w

Lahti L, Sudarshan S. 2012-2019. microbiome: microbiome R package. Github Available from: https://github.com/microbiome/microbiome

Legendre P, Legendre L. 1998. 1998. Numerical ecology. Second English edition. Elsevier, Amsterdam.

Liang R, Lau MCY, Saitta ET, Garvin ZK, Onstott TC. 2020. Genome-centric resolution of novel microbial lineages in an excavated Centrosaurus dinosaur fossil bone from the Late Cretaceous of North America. Environ Microbiome [Internet] 15:8. Available from: 10.1186/s40793-020-00355-w

Li D, Liu C-M, Luo R, Sadakane K, Lam T-W. 2015. MEGAHIT: an ultra-fast single-node solution for large and complex metagenomics assembly via succinct de Bruijn graph. Bioinformatics [Internet] 31:1674–1676. Available from: 10.1093/bioinformatics/btv033

Li H. 2013. Aligning sequence reads, clone sequences and assembly contigs with BWA-MEM. *arXiv [q-bio.GN]* [Internet]. Available from: http://arxiv.org/abs/1303.3997

Lin Z, Chen L, Chen X, Zhong Y, Yang Y, Xia W, Liu C, Zhu W, Wang H, Yan B, et al. 2019. Biological adaptations in the Arctic cervid, the reindeer (Rangifer tarandus). Science [Internet] 364. Available from: 10.1126/science.aav6312

Longin R. 1971. New method of collagen extraction for radiocarbon dating. Nature [Internet] 230:241–242. Available from: 10.1038/230241a0

Lorenzen ED, Nogués-Bravo D, Orlando L, Weinstock J, Binladen J, Marske KA, Ugan A, Borregaard MK, Gilbert MTP, Nielsen R, et al. 2011. Species-specific responses of Late Quaternary megafauna to climate and humans. Nature [Internet] 479:359–364. Available from: 10.1038/nature10574

Mallick H, Rahnavard A, McIver LJ, Ma S, Zhang Y, Nguyen LH, Tickle TL, Weingart G, Ren B, Schwager EH, et al. 2021. Multivariable association discovery in population-scale meta-omics studies. PLoS Comput. Biol. [Internet] 17:e1009442. Available from: 10.1371/journal.pcbi.1009442

Mann AE, Fellows Yates JA, Fagernäs Z, Austin RM, Nelson EA, Hofman CA. 2023. Do I have something in my teeth? The trouble with genetic analyses of diet from archaeological dental calculus. Quat. Int. [Internet] 653–654:33–46. Available from: https://www.sciencedirect.com/science/article/pii/S1040618220307746

Mann AE, Sabin S, Ziesemer K, Vågene ÅJ, Schroeder H, Ozga AT, Sankaranarayanan K, Hofman CA, Fellows Yates JA, Salazar-García DC, et al. 2018. Differential preservation of endogenous human and microbial DNA in dental calculus and dentin. Sci. Rep. [Internet] 8:9822. Available from: 10.1038/s41598-018-28091-9

Mathiesen SD, Aagnes Utsi TH, Sørmo W. 1999. Forage chemistry and the digestive system in reindeer (Rangifer tarandus tarandus) in northern Norway and on South Georgia. Rangifer [Internet] 19:91. Available from: https://septentrio.uit.no/index.php/rangifer/article/view/285

Maury J, Le Bel J-A. 1925. Laugerie Basse: the excavations of M.J.-A. Le Bel. Le Mans: Monnoyer Available from: https://www.worldcat.org/de/title/1742132

McMurdie PJ, Holmes S. 2013. phyloseq: an R package for reproducible interactive analysis and graphics of microbiome census data. PLoS One [Internet] 8:e61217. Available from: 10.1371/journal.pone.0061217

Merceron G, Madelaine S. 2008. Molar microwear pattern and palaeoecology of ungulates from La Berbie (Dordogne, France): environment of Neanderthals and modern human populations of the Middle/Upper Palaeolithic. Boreas [Internet] 35:272–278. Available from: https://onlinelibrary.wiley.com/doi/epdf/10.1111/j.1502-3885.2006.tb01157.x

Michelsen C, Pedersen MW, Fernandez-Guerra A, Zhao L, Petersen TC, Korneliussen TS. 2022. MetaDMG – A fast and accurate ancient DNA damage toolkit for metagenomic data. *bioRxiv* [Internet]:2022.12.06.519264. Available from: https://www.biorxiv.org/content/10.1101/2022.12.06.519264v1.abstract

Mistry J, Chuguransky S, Williams L, Qureshi M, Salazar GA, Sonnhammer ELL, Tosatto SCE, Paladin L, Raj S, Richardson LJ, et al. 2021. Pfam: The protein families database in 2021. Nucleic Acids Res. [Internet] 49:D412–D419. Available from: 10.1093/nar/gkaa913

Modi A, Pisaneschi L, Zaro V, Vai S, Vergata C, Casalone E, Caramelli D, Moggi-Cecchi J, Mariotti Lippi M, Lari M. 2020. Combined methodologies for gaining much information from ancient dental calculus: testing experimental strategies for simultaneously analysing DNA and food residues. Archaeol. Anthropol. Sci. [Internet] 12. Available from: 10.1007/s12520-019-00983-5

Molin G, Ternstrom A. 1986. Phenotypically based taxonomy of psychrotrophic Pseudomonas isolated from spoiled meat, water, and soil. Int. J. Syst. Bacteriol. [Internet] 36:257–274. Available from: https://www.microbiologyresearch.org/content/journal/ijsem/10.1099/00207713-36-2-257

Molin G, Ternstrom A, Ursing J. 1986. Notes: Pseudomonas lundensis, a new bacterial species isolated from meat. Int. J. Syst. Bacteriol. [Internet] 36:339–342. Available from: https://www.microbiologyresearch.org/content/journal/ijsem/10.1099/00207713-36-2-339

Moraitou M, Forsythe A, Fellows Yates JA, Brealey JC, Warinner C, Guschanski K. 2022. Ecology, Not Host Phylogeny, Shapes the Oral Microbiome in Closely Related Species. Mol. Biol. Evol. [Internet] 39. Available from: 10.1093/molbev/msac263

Moraitou M, Richards J, Bolyos C, Saliari K, Gilissen E, Timmons Z, Kitchener AC, Pauwels OSG, Sabin R, Kokkini P, et al. 2025. Host traits impact the outcome of metagenomic library preparation from dental calculus samples across diverse mammals. *bioRxiv* [Internet]:2025.03.19.643754. Available from: https://www.biorxiv.org/content/10.1101/2025.03.19.643754v1.abstract

Müller-Wille L, Heinrich D, Lehtola V-P, Aikio P, Konstantinov Y, Vladimirova V. 2006. Dynamics in Human-Reindeer Relations: Reflections on Prehistoric, Historic and Contemporary Practices in Northernmost Europe. In: Forbes BC, Bölter M, Müller-Wille L, Hukkinen J, Müller F, Gunslay N, Konstantinov Y, editors. Reindeer Management in Northernmost Europe: Linking Practical and Scientific Knowledge in Social-Ecological Systems. Berlin, Heidelberg: Springer Berlin Heidelberg. p. 27–45. Available from: 10.1007/3-540-31392-3_3

Mustonen T. 2022. Wild reindeer as a keystone cultural and ecological species in the Eurasian north. Glob. Chang. Biol. [Internet] 28:4225–4228. Available from: 10.1111/gcb.16203

Nguyen NH, Song Z, Bates ST, Branco S, Tedersoo L, Menke J, Schilling JS, Kennedy PG. 2016. FUNGuild: An open annotation tool for parsing fungal community datasets by ecological guild. Fungal Ecol. [Internet] 20:241–248. Available from: 10.1016/j.funeco.2015.06.006

Nieminen M, Heiskari U. 1989. Diets of freely grazing and captive reindeer during summer and winter. Rangifer [Internet] 9:17. Available from: 10.7557/2.9.1.771

Niven L, Steele TE, Rendu W, Mallye J-B, McPherron SP, Soressi M, Jaubert J, Hublin J-J. 2012. Neandertal mobility and large-game hunting: the exploitation of reindeer during the Quina Mousterian at Chez-Pinaud Jonzac (Charente-Maritime, France). J. Hum. Evol. [Internet] 63:624–635. Available from: 10.1016/j.jhevol.2012.07.002

Oksanen J, Blanchet FG, Kindt R, Legendre P, Minchin PR, O’hara RB, Oksanen M. 2013. Package “vegan.” Community ecology package, version [Internet] 2:1–295. Available from: https://mirror.ibcp.fr/pub/CRAN/web/packages/vegan/vegan.pdf

Olm MR, Brown CT, Brooks B, Banfield JF. 2017. dRep: a tool for fast and accurate genomic comparisons that enables improved genome recovery from metagenomes through de-replication. ISME J. [Internet] 11:2864–2868. Available from: 10.1038/ismej.2017.126

Ondov BD, Treangen TJ, Melsted P, Mallonee AB, Bergman NH, Koren S, Phillippy AM. 2016. Mash: fast genome and metagenome distance estimation using MinHash. Genome Biol. [Internet] 17:132. Available from: 10.1186/s13059-016-0997-x

Oskolkov N, Jin C, Clinton SL, Guinet B, Wijnands F, Johnson E, Kutschera VE, Kinsella CM, Heintzman PD, van der Valk T. 2025. Disinfecting eukaryotic reference genomes to improve taxonomic inference from ancient environmental metagenomic data. *bioRxiv* [Internet]:2025.03.19.644176. Available from: https://www.biorxiv.org/content/10.1101/2025.03.19.644176v1.abstract

Ottoni C, Guellil M, Ozga AT, Stone AC, Kersten O, Bramanti B, Porcier S, Van Neer W. 2019. Metagenomic analysis of dental calculus in ancient Egyptian baboons. Sci. Rep. [Internet] 9:1–10. Available from: 10.1038/s41598-019-56074-x

Ozga AT, Ottoni C. 2023. Dental calculus as a proxy for animal microbiomes. Quat. Int. [Internet] 653–654:47–52. Available from: https://www.sciencedirect.com/science/article/pii/S1040618221003554

Paradis E, Schliep K. 2019. ape 5.0: an environment for modern phylogenetics and evolutionary analyses in R. Bioinformatics [Internet] 35:526–528. Available from: https://academic.oup.com/bioinformatics/article-abstract/35/3/526/5055127

Parks DH, Imelfort M, Skennerton CT, Hugenholtz P, Tyson GW. 2015. CheckM: assessing the quality of microbial genomes recovered from isolates, single cells, and metagenomes. Genome Res. [Internet] 25:1043–1055. Available from: 10.1101/gr.186072.114

Peng X, Wilken SE, Lankiewicz TS, Gilmore SP, Brown JL, Henske JK, Swift CL, Salamov A, Barry K, Grigoriev IV, et al. 2021. Genomic and functional analyses of fungal and bacterial consortia that enable lignocellulose breakdown in goat gut microbiomes. Nat Microbiol [Internet] 6:499–511. Available from: 10.1038/s41564-020-00861-0

Pockrandt C, Zimin AV, Salzberg SL. 2022. Metagenomic classification with KrakenUniq on low-memory computers. J. Open Source Softw. [Internet] 7:4908. Available from: 10.21105/joss.04908

Pope PB, Mackenzie AK, Gregor I, Smith W, Sundset MA, McHardy AC, Morrison M, Eijsink VGH. 2012. Metagenomics of the Svalbard reindeer rumen microbiome reveals abundance of polysaccharide utilization loci. PLoS One [Internet] 7:e38571. Available from: 10.1371/journal.pone.0038571

R Core Team. 2021. R: A Language and Environment for Statistical Computing. Available from: https://www.R-project.org/

Reimer PJ, Austin WEN, Bard E, Bayliss A, Blackwell PG, Bronk Ramsey C, Butzin M, Cheng H, Edwards RL, Friedrich M, et al. 2020. The IntCal20 Northern hemisphere radiocarbon age calibration curve (0–55 cal kBP). Radiocarbon [Internet] 62:725–757. Available from: 10.1017/RDC.2020.41

Richter M, Rosselló-Móra R. 2009. Shifting the genomic gold standard for the prokaryotic species definition. Proc. Natl. Acad. Sci. U. S. A. [Internet] 106:19126–19131. Available from: 10.1073/pnas.0906412106

Rivals F, Drucker DG, Weber M-J, Audouze F, Enloe JG. 2020. Dietary traits and habitats of the reindeer (Rangifer tarandus) during the Late Glacial of Northern Europe. Archaeol. Anthropol. Sci. [Internet] 12. Available from: 10.1007/s12520-020-01052-y

Rivals F, Semprebon GM. 2017. Latitude matters: an examination of behavioural plasticity in dietary traits amongst extant and Pleistocene *Rangifer tarandus*. Boreas [Internet] 46:254–263. Available from: https://onlinelibrary.wiley.com/doi/abs/10.1111/bor.12205

Ryu EP, Davenport ER. 2022. Host genetic determinants of the microbiome across animals: From Caenorhabditis elegans to cattle. Annu. Rev. Anim. Biosci. [Internet] 10:203–226. Available from: https://www.annualreviews.org/content/journals/10.1146/annurev-animal-020420-032054

Saitta ET, Liang R, Lau MCY, Brown CM, Longrich NR, Kaye TG, Novak BJ, Salzberg SL, Norell MA, Abbott GD, et al. 2019. Cretaceous dinosaur bone contains recent organic material and provides an environment conducive to microbial communities. Elife [Internet] 8. Available from: 10.7554/elife.46205

Salgado-Flores A, Hagen LH, Ishaq SL, Zamanzadeh M, Wright A-DG, Pope PB, Sundset MA. 2016. Rumen and Cecum Microbiomes in Reindeer (Rangifer tarandus tarandus) Are Changed in Response to a Lichen Diet and May Affect Enteric Methane Emissions. PLoS One [Internet] 11:e0155213. Available from: 10.1371/journal.pone.0155213

Salter SJ, Cox MJ, Turek EM, Calus ST, Cookson WO, Moffatt MF, Turner P, Parkhill J, Loman NJ, Walker AW. 2014. Reagent and laboratory contamination can critically impact sequence-based microbiome analyses. BMC Biol. [Internet] 12:87. Available from: 10.1186/s12915-014-0087-z

Sayers EW, Beck J, Bolton EE, Brister JR, Chan J, Connor R, Feldgarden M, Fine AM, Funk K, Hoffman J, et al. 2025. Database resources of the National Center for Biotechnology Information in 2025. Nucleic Acids Res. [Internet] 53:D20–D29. Available from: 10.1093/nar/gkae979

Schubert M, Ermini L, Der Sarkissian C, Jónsson H, Ginolhac A, Schaefer R, Martin MD, Fernández R, Kircher M, McCue M, et al. 2014. Characterization of ancient and modern genomes by SNP detection and phylogenomic and metagenomic analysis using PALEOMIX. Nat. Protoc. [Internet] 9:1056–1082. Available from: 10.1038/nprot.2014.063

Seiler M, Grootes PM, Haarsaker J, Lélu S, Rzadeczka-Juga I, Stene S, Svarva H, Thun T, Værnes E, Nadeau M-J. 2019. Status report of the Trondheim radiocarbon laboratory. Radiocarbon [Internet] 61:1963–1972. Available from: 10.1017/rdc.2019.115

Sommer RS, Kalbe J, Ekström J, Benecke N, Liljegren R. 2014. Range dynamics of the reindeer in Europe during the last 25,000 years. J. Biogeogr. [Internet] 41:298–306. Available from: https://onlinelibrary.wiley.com/doi/10.1111/jbi.12193

Sommer RS, Nadachowski A. 2006. Glacial refugia of mammals in Europe: evidence from fossil records. Mamm. Rev. [Internet] 36:251–265. Available from: 10.1111/j.1365-2907.2006.00093.x

Staaland H, Øritsland NA, White RG. 1988. Digestion of energy and nutrients in Svalbard reindeer. Rangifer [Internet] 8:2. Available from: 10.7557/2.8.1.725

Stewart RD, Auffret MD, Warr A, Wiser AH, Press MO, Langford KW, Liachko I, Snelling TJ, Dewhurst RJ, Walker AW, et al. 2018. Assembly of 913 microbial genomes from metagenomic sequencing of the cow rumen. Nat. Commun. [Internet] 9:870. Available from: 10.1038/s41467-018-03317-6

Stien A, Irvine RJ, Ropstad E, Halvorsen O, Langvatn R, Albon SD. 2002. The impact of gastrointestinal nematodes on wild reindeer: experimental and cross-sectional studies. J. Anim. Ecol. [Internet] 71:937–945. Available from: https://besjournals.onlinelibrary.wiley.com/doi/10.1046/j.1365-2656.2002.00659.x

Storeheier PV, Mathiesen SD, Tyler NJC, Olsen MA. 2002. Nutritive value of terricolous lichens for reindeer in winter. Lichenologist [Internet] 34:247–257. Available from: 10.1006/lich.2002.0394

Sundset MA, Edwards JE, Cheng YF, Senosiain RS, Fraile MN, Northwood KS, Praesteng KE, Glad T, Mathiesen SD, Wright A-DG. 2009a. Molecular diversity of the rumen microbiome of Norwegian reindeer on natural summer pasture. Microb. Ecol. [Internet] 57:335–348. Available from: 10.1007/s00248-008-9414-7

Sundset MA, Edwards JE, Cheng YF, Senosiain RS, Fraile MN, Northwood KS, Praesteng KE, Glad T, Mathiesen SD, Wright A-DG. 2009b. Rumen microbial diversity in Svalbard reindeer, with particular emphasis on methanogenic archaea. FEMS Microbiol. Ecol. [Internet] 70:553–562. Available from: 10.1111/j.1574-6941.2009.00750.x

Tapio I, Shingfield KJ, McKain N, Bonin A, Fischer D, Bayat AR, Vilkki J, Taberlet P, Snelling TJ, Wallace RJ. 2016. Oral Samples as Non-Invasive Proxies for Assessing the Composition of the Rumen Microbial Community. PLoS One [Internet] 11:e0151220. Available from: 10.1371/journal.pone.0151220

Taylor RS, Horn RL, Zhang X, Golding GB, Manseau M, Wilson PJ. 2019. The Caribou (Rangifer tarandus) Genome. Genes [Internet] 10:540. Available from: 10.3390/genes10070540

Tian Z, Lu S, Jin D, Yang J, Pu J, Lai X-H, Bai X-N, Wu X-M, Li J, Wang S, et al. 2019. Streptococcus chenjunshii sp. nov. isolated from feces of Tibetan antelopes. Int. J. Syst. Evol. Microbiol. [Internet] 69:1237–1243. Available from: 10.1099/ijsem.0.003303

Uritskiy GV, DiRuggiero J, Taylor J. 2018. MetaWRAP-a flexible pipeline for genome-resolved metagenomic data analysis. Microbiome [Internet] 6:158. Available from: 10.1186/s40168-018-0541-1

Utter DR, Mark Welch JL, Borisy GG. 2016. Individuality, Stability, and Variability of the Plaque Microbiome. Front. Microbiol. [Internet] 7:564. Available from: 10.3389/fmicb.2016.00564

Vaidya JD, Hornung BVH, Smidt H, Edwards JE, Plugge CM. 2019. Propionibacterium ruminifibrarum sp. nov., isolated from cow rumen fibrous content. Int. J. Syst. Evol. Microbiol. [Internet] 69:2584–2590. Available from: 10.1099/ijsem.0.003544

Veiberg V, Nilsen EB, Rolandsen CM, Heim M, Andersen R, Holmstrøm F, Meisingset EL, Solberg EJ. 2020. The accuracy and precision of age determination by dental cementum annuli in four northern cervids. Eur. J. Wildl. Res. [Internet] 66:91. Available from: 10.1007/s10344-020-01431-9

Velsko IM, Frantz LAF, Herbig A, Larson G, Warinner C. 2018. Selection of Appropriate Metagenome Taxonomic Classifiers for Ancient Microbiome Research. mSystems [Internet] 3:1–23. Available from: 10.1128/msystems.00080-18

Vereshchagin NK, Baryshnikov GF. 1992. The ecological structure of the mammoth fauna. Ann. Zool. Fennici 28:253–259.

Wang Y, Pedersen MW, Alsos IG, De Sanctis B, Racimo F, Prohaska A, Coissac E, Owens HL, Merkel MKF, Fernandez-Guerra A, et al. 2021. Late Quaternary dynamics of Arctic biota from ancient environmental genomics. Nature [Internet] 600:86–92. Available from: 10.1038/s41586-021-04016-x

Warinner C, Herbig A, Mann A, Fellows Yates JA, Weiß CL, Burbano HA, Orlando L, Krause J. 2017. A Robust Framework for Microbial Archaeology. Annu. Rev. Genomics Hum. Genet. [Internet] 18:321–356. Available from: 10.1146/annurev-genom-091416-035526

Warinner C, Rodrigues JFM, Vyas R, Trachsel C, Shved N, Grossmann J, Radini A, Hancock Y, Tito RY, Fiddyment S, et al. 2014. Pathogens and host immunity in the ancient human oral cavity. Nat. Genet. [Internet] 46:336–344. Available from: 10.1038/ng.2906

Warinner C, Speller C, Collins MJ. 2015. A new era in palaeomicrobiology: prospects for ancient dental calculus as a long-term record of the human oral microbiome. Philos. Trans. R. Soc. Lond. B Biol. Sci. [Internet] 370:20130376. Available from: 10.1098/rstb.2013.0376

Wei Z, Xu Z, Qiao T, Wang S, Ishwaran N, Yang M. 2023. Habitats change of Tibetan antelope and its influencing factors on the North Tibetan Plateau from 2020 to 2050. Global Ecology and Conservation [Internet] 43:e02462. Available from: https://www.sciencedirect.com/science/article/pii/S2351989423000975

Weyrich LS, Duchene S, Soubrier J, Arriola L, Llamas B, Breen J, Morris AG, Alt KW, Caramelli D, Dresely V, et al. 2017. Neanderthal behaviour, diet, and disease inferred from ancient DNA in dental calculus. Nature [Internet] 544:357–361. Available from: https://www.nature.com/articles/nature21674

Weyrich LS, Farrer AG, Eisenhofer R, Arriola LA, Young J, Selway CA, Handsley-Davis M, Adler CJ, Breen J, Cooper A. 2019. Laboratory contamination over time during low-biomass sample analysis. Mol. Ecol. Resour. [Internet] 19:982–996. Available from: 10.1111/1755-0998.13011

Wickham H, Chang W, Wickham MH. 2016. Package “ggplot2.” Create elegant data visualisations using the grammar of graphics. Version [Internet] 2:1–189. Available from: https://citeseerx.ist.psu.edu/document?repid=rep1&type=pdf&doi=af53fd2f5b9e81b6edec0c13e1b3babd34bda399

Wu Y-W, Simmons BA, Singer SW. 2016. MaxBin 2.0: an automated binning algorithm to recover genomes from multiple metagenomic datasets. Bioinformatics [Internet] 32:605–607. Available from: 10.1093/bioinformatics/btv638

Zielińska S, Kidawa D, Stempniewicz L, Łoś M, Łoś JM. 2016. New Insights into the Microbiota of the Svalbard Reindeer Rangifer tarandus platyrhynchus. Front. Microbiol. [Internet] 7:170. Available from: 10.3389/fmicb.2016.00170

Ziesemer KA, Mann AE, Sankaranarayanan K, Schroeder H, Ozga AT, Brandt BW, Zaura E, Waters-Rist A, Hoogland M, Salazar-García DC, et al. 2015. Intrinsic challenges in ancient microbiome reconstruction using 16S rRNA gene amplification. Sci. Rep. [Internet] 5:16498. Available from: 10.1038/srep16498

