## Supplementary Material for "Dental calculus as a record of Pleistocene reindeer oral, digestive and dietary flora"

Supplementary Table 1. Overview over all assembled metagenome-assembled genomes (MAGs).

Abundance is filtered for detection (minimum 0.3) and summed up across all samples. If detection is below the threshold, abundance is set to 0 (= not present). Completion, redundancy (contamination), GC content, $N_{50}$ and total length were calculated with checkM (Parks et al. 2015). Taxa which were identified as lab contaminants with decontam (Davis et al. 2018) are removed.

| Co-Assembly batch | BIN | OTU | Sum abundance | Completion [%] | Contamination [%] | GC [%] | N50 | Total length [bp] | domain | phylum | class | order | family | genus | species |
|---|---|---|---|---|---|---|---|---|---|---|---|---|---|---|---|
| MODERN | bin_32_mod | Actinomyces sp. | 91.6 | 96.4 | 2.2 | 70.5 | 30226 | 3644339 | Bacteria | Actinobacteriota | Actinomycetia | Actinomycetales | Actinomycetaceae | Actinomyces | |
| MODERN | bin_61_mod | Actinomyces sp. | 186.7 | 81.4 | 6.6 | 67.2 | 9587 | 3175201 | Bacteria | Actinobacteriota | Actinomycetia | Actinomycetales | Actinomycetaceae | Actinomyces | |
| MODERN | bin_46_mod | Actinomyces sp. | 35.6 | 53.8 | 2.4 | 69 | 4500 | 1828876 | Bacteria | Actinobacteriota | Actinomycetia | Actinomycetales | Actinomycetaceae | Actinomyces | |
| ANCIENT | bin_19_anc | Allosaccharopolyspora sp. | 12.5 | 86.3 | 6.0 | 67.7 | 21599 | 4697505 | Bacteria | Actinobacteriota | Actinomycetia | Mycobacteriales | Pseudonocardiaceae | Allosaccharopolyspora | |
| ANCIENT | bin_23_anc | Allosaccharopolyspora sp | 14.5 | 80.9 | 3.9 | 66.7 | 18237 | 4092176 | Bacteria | Actinobacteriota | Actinomycetia | Mycobacteriales | Pseudonocardiaceae | Allosaccharopolyspora | |
| ANCIENT | bin_3_anc | Allosaccharopolyspora sp | 48.6 | 60.8 | 8.6 | 69.3 | 3491 | 5315633 | Bacteria | Actinobacteriota | Actinomycetia | Mycobacteriales | Pseudonocardiaceae | Allosaccharopolyspora | |
| ANCIENT | bin_35_anc | Allosaccharopolyspora sp | 18.3 | 85.4 | 6.8 | 68.7 | 17034 | 4732674 | Bacteria | Actinobacteriota | Actinomycetia | Mycobacteriales | Pseudonocardiaceae | Allosaccharopolyspora | |
| ANCIENT | bin_48_anc | Allosaccharopolyspora sp | 5.2 | 63.7 | 3.7 | 66.3 | 3925 | 4738523 | Bacteria | Actinobacteriota | Actinomycetia | Mycobacteriales | Pseudonocardiaceae | Allosaccharopolyspora | |
| ANCIENT | bin_6_anc | Amycolatopsis_C sp009377395 | 9.7 | 51.2 | 0.0 | 70.2 | 4322 | 3353510 | Bacteria | Actinobacteriota | Actinomycetia | Mycobacteriales | Pseudonocardiaceae | Amycolatopsis_C | Amycolatopsis_C sp009377395 |
| ANCIE | bin_ | Asp313 | 146.2 | 57.0 | 2.9 | 61. | 3104 | 15888 | Bacteri | Actino | Actino | Actino | Actino | Asp31 | |

| Co-Assembly batch | BIN | OTU | Sum abundance | Completion [%] | Contamination [%] | GC [%] | N50 | Total length [bp] | domain | phylum | class | order | family | genus | species |
|---|---|---|---|---|---|---|---|---|---|---|---|---|---|---|---|
| NT | F_dup | sp. | | | | 1 | | 55 | a | bacteriota | mycetia | mycetales | mycetaceae | 3 | |
| MODERN | bin_F_dup | Asp313 sp. | 146.2 | 95.5 | 3.1 | 61.3 | 12465 | 2503362 | Bacteria | Actinobacteriota | Actinomycetia | Actinomycetales | Actinomycetaceae | Asp313 | |
| MODERN | bin_45_mod | Bacteroides sp. | 9.0 | 97.6 | 0.1 | 47.6 | 60945 | 2853739 | Bacteria | Bacteroidota | Bacteroidia | Bacteroidales | Bacteroidaceae | Bacteroides | |
| ANCIENT | bin_A_dup | Basfia_A sp. | 162.7 | 86.5 | 4.5 | 44.6 | 3432 | 2032619 | Bacteria | Proteobacteria | Gammaproteobacteria | Enterobacterales | Pasteurellaceae | Basfia_A | |
| MODERN | bin_A_dup | Basfia_A sp. | 162.7 | 99.2 | 3.0 | 44.4 | 40885 | 2463674 | Bacteria | Proteobacteria | Gammaproteobacteria | Enterobacterales | Pasteurellaceae | Basfia_A | |
| MODERN | bin_49_mod | Bibersteinia sp. | 8.6 | 87.5 | 2.4 | 41.4 | 9861 | 1781199 | Bacteria | Proteobacteria | Gammaproteobacteria | Enterobacterales | Pasteurellaceae | Bibersteinia | |
| ANCIENT | bin_45_anc | Bilifractor sp. | 134.6 | 96.5 | 4.2 | 46.3 | 9104 | 2771729 | Bacteria | Firmicutes_A | Clostridia | Lachnospirales | Lachnospiraceae | Bilifractor | |
| MODERN | bin_11_mod | Brachymonas sp. | 14.0 | 91.8 | 1.3 | 51.1 | 20494 | 1932991 | Bacteria | Proteobacteria | Gammaproteobacteria | Burkholderiales | Burkholderiaceae | Brachymonas | |
| ANCIENT | bin_31_anc | CAG-791 sp. | 61.6 | 74.0 | 8.0 | 51.4 | 2206 | 2210786 | Bacteria | Firmicutes_A | Clostridia | Lachnospirales | Lachnospiraceae | CAG-791 | |
| MODERN | bin_16_mod | CAJPNX01 sp. | 6.7 | 71.0 | 1.5 | 51.8 | 3916 | 1346648 | Bacteria | Proteobacteria | Gammaproteobacteria | Burkholderiales | Neisseriaceae | CAJPNX01 | |
| MODERN | bin_30_mod | Campylobacter_B sp. | 1.2 | 88.0 | 2.6 | 37 | 3773 | 1857269 | Bacteria | Campylobacterota | Campylobacteria | Campylobacterales | Campylobacteraceae | Campylobacter_B | |
| MODERN | bin_22_mod | Chryseobacterium piscium | 1.6 | 51.3 | 2.0 | 38.4 | 1507 | 2718075 | Bacteria | Bacteroidota | Bacteroidia | Flavobacteriales | Weeksellaceae | Chryseobacterium | Chryseobacterium piscium |
| MODERN | bin_ | Cryptobac | 10.8 | 93.6 | 0.6 | 53. | 29385 | 19695 | Bacteri | Bacter | Bacter | Bacter | UBA93 | Crypto | |

| Co-Assembly batch | BIN | OTU | Sum abundance | Completion [%] | Contamination [%] | GC [%] | N50 | Total length [bp] | domain | phylum | class | order | family | genus | species |
|---|---|---|---|---|---|---|---|---|---|---|---|---|---|---|---|
| RN | 29_mod | teroides sp. | | | | 6 | | 42 | a | oidota | oidia | oidales | 2 | bacteroides | |
| MODERN | bin_54_mod | Cryptobacteroides sp | 1.9 | 54.2 | 2.8 | 55.5 | 1621 | 1646386 | Bacteria | Bacteroidota | Bacteroidia | Bacteroidales | UBA932 | Cryptobacteroides | |
| ANCIENT | bin_58_anc | CSP1-5 sp. | 1.4 | 56.3 | 15.0 | 64.9 | 1926 | 1462166 | Bacteria | Methylomirabilota | Methylomirabilia | Methylomirabilales | CSP1-5 | CSP1-5 | |
| MODERN | bin_12_mod | Desulfovibrio desulfuricans_D | 17.6 | 97.0 | 0.6 | 63.8 | 36484 | 2681036 | Bacteria | Desulfobacterota_I | Desulfovibrionia | Desulfovibrionales | Desulfovibrionaceae | Desulfovibrio | Desulfovibrio desulfuricans_D |
| ANCIENT | bin_52_anc | DP-1 sp. | 4.6 | 66.5 | 3.0 | 55.4 | 2224 | 5408552 | Bacteria | Desulfobacterota_B | Binatia | UBA9968 | UBA9968 | DP-1 | |
| MODERN | bin_18_mod | Eubacterium_I sp. | 14.2 | 96.3 | 2.6 | 47.9 | 20511 | 2242840 | Bacteria | Firmicutes_A | Clostridia | Lachnospirales | Lachnospiraceae | Eubacterium_I | |
| ANCIENT | bin_22_anc | Eubacterium_I sp | 13.5 | 69.3 | 4.8 | 48.5 | 2388 | 1857276 | Bacteria | Firmicutes_A | Clostridia | Lachnospirales | Lachnospiraceae | Eubacterium_I | |
| MODERN | bin_20_mod | Flavobacterium sp. | 1.9 | 54.9 | 1.5 | 38.2 | 1635 | 1813077 | Bacteria | Bacteroidota | Bacteroidia | Flavobacteriales | Flavobacteriaceae | Flavobacterium | |
| MODERN | bin_21_mod | Flavobacterium sp | 5.1 | 97.0 | 0.5 | 34 | 14395 | 3772220 | Bacteria | Bacteroidota | Bacteroidia | Flavobacteriales | Flavobacteriaceae | Flavobacterium | |
| ANCIENT | bin_61_anc | Flavobacterium sp | 53.2 | 68.9 | 2.2 | 33.5 | 2960 | 2292418 | Bacteria | Bacteroidota | Bacteroidia | Flavobacteriales | Flavobacteriaceae | Flavobacterium | |
| MODERN | bin_10_mod | Fusobacterium sp. | 3.3 | 75.6 | 4.0 | 27.7 | 4517 | 1610384 | Bacteria | Fusobacteriota | Fusobacteriia | Fusobacteriales | Fusobacteriaceae | Fusobacterium | |
| MODERN | bin_15_mod | Fusobacterium sp | 8.3 | 96.1 | 4.4 | 27.9 | 14777 | 2110346 | Bacteria | Fusobacteriota | Fusobacteriia | Fusobacteriales | Fusobacteriaceae | Fusobacterium | |
| MODERN | bin_50_mod | Fusobacterium_C necrophorum | 4.1 | 98.3 | 1.7 | 35.1 | 15284 | 2011259 | Bacteria | Fusobacteriota | Fusobacteriia | Fusobacteriales | Fusobacteriaceae | Fusobacterium_C | Fusobacterium_C necrophorum |

| Co-Assembly batch | BIN | OTU | Sum abundance | Completion [%] | Contamination [%] | GC [%] | N50 | Total length [bp] | domain | phylum | class | order | family | genus | species |
|---|---|---|---|---|---|---|---|---|---|---|---|---|---|---|---|
| ANCIENT | bin_29_anc | Haemophilus_B sp. | 72.5 | 93.3 | 3.0 | 46.5 | 9924 | 2121583 | Bacteria | Proteobacteria | Gammaproteobacteria | Enterobacterales | Pasteurellaceae | Haemophilus_B | |
| ANCIENT | bin_11_anc | Homoserinimonas sp. | 6.5 | 85.6 | 4.2 | 65 | 9895 | 2715953 | Bacteria | Actinobacteriota | Actinomycetia | Actinomycetales | Microbacteriaceae | Homoserinimonas | |
| ANCIENT | bin_47_anc | Hornefia sp. | 10.9 | 52.9 | 6.0 | 57 | 1448 | 1778449 | Bacteria | Firmicutes_A | Clostridia | Peptostreptococcales | Anaerovoracaceae | Hornefia | |
| ANCIENT | bin_46_anc | JACQPT01 sp. | 11.0 | 97.5 | 3.3 | 50.2 | 70727 | 4230797 | Bacteria | Bacteroidota | UBA10030 | UBA10030 | UBA10030 | JACQPT01 | |
| MODERN | bin_24_mod | JAGOWQ01 sp. | 16.0 | 98.9 | 1.1 | 26.1 | 31295 | 1626520 | Bacteria | Fusobacteriota | Fusobacteriia | Fusobacteriales | Leptotrichiaceae | JAGOWQ01 | |
| MODERN | bin_7_mod | JAGOWQ01 sp | 130.7 | 95.5 | 2.2 | 30.1 | 20386 | 2117629 | Bacteria | Fusobacteriota | Fusobacteriia | Fusobacteriales | Leptotrichiaceae | JAGOWQ01 | |
| ANCIENT | bin_57_anc | Lactobacillaceae | 10.8 | 52.7 | 2.5 | 27.6 | 4214 | 582164 | Bacteria | Firmicutes | Bacilli | Lactobacillales | Lactobacillaceae | | |
| ANCIENT | bin_49_anc | Lactobacillus intestinalis | 18.8 | 76.3 | 0.1 | 35.7 | 4625 | 1284317 | Bacteria | Firmicutes | Bacilli | Lactobacillales | Lactobacillaceae | Lactobacillus | Lactobacillus intestinalis |
| ANCIENT | bin_21_anc | Lactobacillus johnsonii | 1.9 | 52.7 | 12.2 | 38 | 1287 | 1914068 | Bacteria | Firmicutes | Bacilli | Lactobacillales | Lactobacillaceae | Lactobacillus | Lactobacillus johnsonii |
| MODERN | bin_51_mod | Lautropia sp. | 91.5 | 90.9 | 7.2 | 62.3 | 6775 | 3509295 | Bacteria | Proteobacteria | Gammaproteobacteria | Burkholderiales | Burkholderiaceae | Lautropia | |
| ANCIENT | bin_34_anc | Ligilactobacillus murinus | 8.0 | 79.8 | 1.3 | 40.8 | 4129 | 1250582 | Bacteria | Firmicutes | Bacilli | Lactobacillales | Lactobacillaceae | Ligilactobacillus | Ligilactobacillus murinus |
| MODERN | bin_3_mod | Mannheimia sp. | 7.0 | 84.0 | 5.9 | 39.4 | 7821 | 1829725 | Bacteria | Proteobacteria | Gammaproteobacteria | Enterobacterales | Pasteurellaceae | Mannheimia | |

| Co-Assembly batch | BIN | OTU | Sum abundance | Completion [%] | Contamination [%] | GC [%] | N50 | Total length [bp] | domain | phylum | class | order | family | genus | species |
|---|---|---|---|---|---|---|---|---|---|---|---|---|---|---|---|
| MODERN | bin_57_mod | Mannheimia sp | 509.9 | 92.8 | 9.9 | 37.5 | 17334 | 1850535 | Bacteria | Proteobacteria | Gammaproteobacteria | Enterobacterales | Pasteurellaceae | Mannheimia | |
| MODERN | bin_9_mod | Mesocricetibacter sp. | 34.2 | 98.7 | 0.8 | 47.8 | 37339 | 2757061 | Bacteria | Proteobacteria | Gammaproteobacteria | Enterobacterales | Pasteurellaceae | Mesocricetibacter | |
| ANCIENT | bin_38_anc | Methanobrevibacter sp. | 50.1 | 71.0 | 2.4 | 32.7 | 4540 | 1560997 | Archaea | Methanobacteriota | Methanobacteria | Methanobacteriales | Methanobacteriaceae | Methanobrevibacter | |
| ANCIENT | bin_33_anc | Methanobrevibacter_A sp. | 18.1 | 85.7 | 5.7 | 30.6 | 7432 | 1973885 | Archaea | Methanobacteriota | Methanobacteria | Methanobacteriales | Methanobacteriaceae | Methanobrevibacter_A | |
| ANCIENT | bin_41_anc | Methanobrevibacter_A sp | 7.8 | 78.3 | 3.2 | 28.1 | 3608 | 1516164 | Archaea | Methanobacteriota | Methanobacteria | Methanobacteriales | Methanobacteriaceae | Methanobrevibacter_A | |
| MODERN | bin_6_mod | Methanomethylophilus sp. | 2.1 | 59.8 | 8.3 | 58.8 | 1678 | 2017389 | Archaea | Thermoplasmatota | Thermoplasmata | Methanomassiliicoccales | Methanomethylophilaceae | Methanomethylophilus | |
| ANCIENT | bin_28_anc | Methanosphaera sp. | 95.1 | 88.6 | 5.0 | 32.5 | 11685 | 1792003 | Archaea | Methanobacteriota | Methanobacteria | Methanobacteriales | Methanobacteriaceae | Methanosphaera | |
| ANCIENT | bin_4_anc | Methanosphaera sp | 51.6 | 84.4 | 2.9 | 27.8 | 10793 | 2063892 | Archaea | Methanobacteriota | Methanobacteria | Methanobacteriales | Methanobacteriaceae | Methanosphaera | |
| ANCIENT | bin_42_anc | Nanosyncoccus sp. | 6.1 | 56.9 | 0.2 | 43.2 | 5751 | 533024 | Bacteria | Patescibacteria | Saccharimonadia | Saccharimonadales | Nanosyncoccaceae | Nanosyncoccus | |
| MODERN | bin_4_mod | Neisseria sp. | 22.7 | 96.5 | 11.5 | 53.9 | 7616 | 2331604 | Bacteria | Proteobacteria | Gammaproteobacteria | Burkholderiales | Neisseriaceae | Neisseria | |
| ANCIENT | bin_27_anc | NS-7 sp. | 3.2 | 86.3 | 4.6 | 58.6 | 9995 | 2678817 | Bacteria | Nitrospirota | Nitrospiria | Nitrospirales | Nitrospiraceae | NS-7 | |
| ANCIENT | bin_9_a | Oceanobacillus sp. | 1.5 | 57.8 | 2.1 | 36.6 | 1756 | 3333676 | Bacteria | Firmicutes | Bacilli | Bacillales_D | Amphibacilla | Oceanobacill | |

| Co-Assembly batch | BIN | OTU | Sum abundance | Completion [%] | Contamination [%] | GC [%] | N50 | Total length [bp] | domain | phylum | class | order | family | genus | species |
|---|---|---|---|---|---|---|---|---|---|---|---|---|---|---|---|
| | nc | | | | | | | | | | | | ceae | us | |
| ANCIENT | bin_64_anc | Paenisporosarcina sp. | 2.3 | 73.6 | 6.6 | 37.3 | 3419 | 3063473 | Bacteria | Firmicutes | Bacilli | Bacillales_A | Planococcaceae | Paenisporosarcina | |
| ANCIENT | bin_50_anc | Pararheinheimera sp. | 3.4 | 77.1 | 0.0 | 52.4 | 4858 | 3828432 | Bacteria | Proteobacteria | Gammaproteobacteria | Enterobacterales | Alteromonadaceae | Pararheinheimera | |
| ANCIENT | bin_63_anc | Pararheinheimera tangshanensis | 1.7 | 68.2 | 4.0 | 46.6 | 2091 | 2670909 | Bacteria | Proteobacteria | Gammaproteobacteria | Enterobacterales | Alteromonadaceae | Pararheinheimera | Pararheinheimera tangshanensis |
| MODERN | bin_34_mod | Peptidiphaga sp. | 136.1 | 95.4 | 0.0 | 65.7 | 64087 | 2306248 | Bacteria | Actinobacteriota | Actinomycetia | Actinomycetales | Actinomycetaceae | Peptidiphaga | |
| ANCIENT | bin_10_anc | Pontibacter sp. | 3.1 | 90.7 | 2.4 | 44 | 9918 | 3941209 | Bacteria | Bacteroidota | Bacteroidia | Cytophagales | Hymenobacteraceae | Pontibacter | |
| MODERN | bin_26_mod | Porphyromonas_A sp. | 3.3 | 94.1 | 1.7 | 48.7 | 8557 | 1849389 | Bacteria | Bacteroidota | Bacteroidia | Bacteroidales | Porphyromonadaceae | Porphyromonas_A | |
| MODERN | bin_17_mod | Prevotella sp. | 6.4 | 99.3 | 0.5 | 49.3 | 57965 | 2990096 | Bacteria | Bacteroidota | Bacteroidia | Bacteroidales | Bacteroidaceae | Prevotella | |
| MODERN | bin_42_mod | Prevotella sp | 20.5 | 94.8 | 16.6 | 50.2 | 3512 | 4734899 | Bacteria | Bacteroidota | Bacteroidia | Bacteroidales | Bacteroidaceae | Prevotella | |
| MODERN | bin_44_mod | Prevotella sp | 1.3 | 51.0 | 3.4 | 52.7 | 1848 | 1855959 | Bacteria | Bacteroidota | Bacteroidia | Bacteroidales | Bacteroidaceae | Prevotella | |
| MODERN | bin_8_mod | Prevotella sp | 4.0 | 83.5 | 3.5 | 53.8 | 12697 | 2512812 | Bacteria | Bacteroidota | Bacteroidia | Bacteroidales | Bacteroidaceae | Prevotella | |
| MODERN | bin_27_mod | Prevotella sp017627765 | 1.9 | 79.6 | 2.5 | 42.1 | 4848 | 2174706 | Bacteria | Bacteroidota | Bacteroidia | Bacteroidales | Bacteroidaceae | Prevotella | Prevotella sp017627765 |
| MODERN | bin_31_mod | Prevotella sp905235835 | 3.5 | 61.8 | 1.9 | 47.8 | 3220 | 1699011 | Bacteria | Bacteroidota | Bacteroidia | Bacteroidales | Bacteroidaceae | Prevotella | Prevotella sp9052 |

| Co-Assembly batch | BIN | OTU | Sum abundance | Completion [%] | Contamination [%] | GC [%] | N50 | Total length [bp] | domain | phylum | class | order | family | genus | species |
|---|---|---|---|---|---|---|---|---|---|---|---|---|---|---|---|
| | | | | | | | | | | | | | | | 35835 |
| ANCIENT | bin_E_dup | Propionibacterium ruminifibrarum | 403.4 | 55.6 | 3.5 | 67.8 | 2769 | 1730795 | Bacteria | Actinobacteriota | Actinomycetia | Propionibacteriales | Propionibacteriaceae | Propionibacterium | Propionibacterium sp900289195 |
| MODERN | bin_E_dup | Propionibacterium ruminifibrarum | 403.4 | 96.8 | 3.6 | 68.7 | 12018 | 2682438 | Bacteria | Actinobacteriota | Actinomycetia | Propionibacteriales | Propionibacteriaceae | Propionibacterium | Propionibacterium sp900289195 |
| MODERN | bin_2_mod | Proteiniphilum sp. | 5.6 | 90.4 | 3.0 | 45.5 | 7248 | 3496335 | Bacteria | Bacteroidota | Bacteroidia | Bacteroidales | Dysgonomonadaceae | Proteiniphilum | |
| ANCIENT | bin_39_anc | Pseudarthrobacter sp. | 3.4 | 60.8 | 4.3 | 65.4 | 2460 | 2861831 | Bacteria | Actinobacteriota | Actinomycetia | Actinomycetales | Micrococcaceae | Pseudarthrobacter | |
| ANCIENT | bin_37_anc | Pseudomonas_E lundensis | 61.0 | 97.7 | 6.8 | 58.8 | 23369 | 4855647 | Bacteria | Proteobacteria | Gammaproteobacteria | Pseudomonadales | Pseudomonadaceae | Pseudomonas_E | Pseudomonas_E lundensis |
| MODERN | bin_5_mod | Pseudomonas_E paraversuta | 15.7 | 70.5 | 15.6 | 56.4 | 3152 | 980793 | Bacteria | Proteobacteria | Gammaproteobacteria | Pseudomonadales | Pseudomonadaceae | Pseudomonas_E | Pseudomonas_E paraversuta |
| MODERN | bin_23_mod | Pseudomonas_E sp003014915 | 6.1 | 58.3 | 6.9 | 60.3 | 2515 | 7413950 | Bacteria | Proteobacteria | Gammaproteobacteria | Pseudomonadales | Pseudomonadaceae | Pseudomonas_E | Pseudomonas_E sp003014915 |
| MODERN | bin_41_mod | Psychrobacter sp. | 8.7 | 89.9 | 2.9 | 42.8 | 6134 | 2879938 | Bacteria | Proteobacteria | Gammaproteobacteria | Pseudomonadales | Moraxellaceae | Psychrobacter | |
| ANCIENT | bin_26_anc | Rickettsia felis | 8.8 | 81.4 | 1.3 | 32.9 | 3807 | 1160887 | Bacteria | Proteobacteria | Alphaproteobacteria | Rickettsiales | Rickettsiaceae | Rickettsia | Rickettsia felis |
| ANCIENT | bin_D_dup | RUG099 sp. | 89.5 | 90.7 | 4.9 | 48.5 | 10083 | 2113659 | Bacteria | Firmicutes_A | Clostridia | Peptostreptococcales | Anaerovoracaceae | RUG099 | |

| Co-Assembly batch | BIN | OTU | Sum abundance | Completion [%] | Contamination [%] | GC [%] | N50 | Total length [bp] | domain | phylum | class | order | family | genus | species |
|---|---|---|---|---|---|---|---|---|---|---|---|---|---|---|---|
| MODERN | bin_D_dup | RUG099 sp. | 89.5 | 69.2 | 4.8 | 48.4 | 3745 | 1692916 | Bacteria | Firmicutes_A | Clostridia | Peptostreptococcales | Anaerovoracaceae | RUG099 | |
| ANCIENT | bin_16_anc | RUG099 sp | 16.3 | 61.5 | 3.4 | 46.2 | 3484 | 1081831 | Bacteria | Firmicutes_A | Clostridia | Peptostreptococcales | Anaerovoracaceae | RUG099 | |
| ANCIENT | bin_36_anc | RUG11795 sp. | 122.7 | 68.0 | 2.0 | 39.5 | 3145 | 1667149 | Bacteria | Firmicutes | Bacilli | Erysipelotrichales | Erysipelotrichaceae | RUG11795 | |
| MODERN | bin_43_mod | RUG721 sp. | 9.2 | 95.0 | 4.9 | 63.2 | 28527 | 2171499 | Bacteria | Actinobacteriota | Coriobacteriia | Coriobacteriales | Atopobiaceae | RUG721 | |
| MODERN | bin_37_mod | Saccharimonas sp. | 9.0 | 63.7 | 0.4 | 48.2 | 129064 | 736272 | Bacteria | Patescibacteria | Saccharimonadia | Saccharimonadales | Saccharimonadaceae | Saccharimonas | |
| ANCIENT | bin_55_anc | Saccharopolyspora dendranthemae | 21.6 | 85.4 | 5.4 | 69.5 | 16887 | 5573339 | Bacteria | Actinobacteriota | Actinomycetia | Mycobacteriales | Pseudonocardiaceae | Saccharopolyspora | Saccharopolyspora dendranthemae |
| ANCIENT | bin_24_anc | SFGY01 sp. | 14.1 | 80.2 | 2.1 | 53.9 | 4358 | 1557077 | Bacteria | Firmicutes_A | Clostridia | Saccharofermentanales | DTU023 | SFGY01 | |
| ANCIENT | bin_15_anc | Shewanella sp002966515 | 15.4 | 70.3 | 9.6 | 46.5 | 1939 | 4592395 | Bacteria | Proteobacteria | Gammaproteobacteria | Enterobacterales | Shewanellaceae | Shewanella | Shewanella sp002966515 |
| ANCIENT | bin_59_anc | Staphylococcus equorum_A | 7.7 | 85.0 | 10.9 | 32.4 | 3102 | 3331866 | Bacteria | Firmicutes | Bacilli | Staphylococcales | Staphylococcaceae | Staphylococcus | Staphylococcus equorum_A |
| ANCIENT | bin_18_anc | Steroidobacter_A sp. | 3.2 | 75.3 | 6.8 | 63 | 2830 | 4469609 | Bacteria | Proteobacteria | Gammaproteobacteria | Steroidobacterales | Steroidobacteraceae | Steroidobacter_A | |
| ANCIENT | bin_14_anc | Steroidobacteraceae | 6.0 | 94.4 | 5.9 | 65.5 | 22248 | 4692371 | Bacteria | Proteobacteria | Gammaproteobacteria | Steroidobacterales | Steroidobacteraceae | | |

| Co-Assembly batch | BIN | OTU | Sum abundance | Completion [%] | Contamination [%] | GC [%] | N50 | Total length [bp] | domain | phylum | class | order | family | genus | species |
|---|---|---|---|---|---|---|---|---|---|---|---|---|---|---|---|
| ANCIENT | bin_B_dup | Streptococcus chenjunshii | 316.5 | 88.2 | 3.8 | 42.6 | 5709 | 2030032 | Bacteria | Firmicutes | Bacilli | Lactobacillales | Streptococcaceae | Streptococcus | Streptococcus chenjunshii |
| MODERN | bin_B_dup | Streptococcus chenjunshii | 316.5 | 95.1 | 0.2 | 41.6 | 27749 | 2162607 | Bacteria | Firmicutes | Bacilli | Lactobacillales | Streptococcaceae | Streptococcus | Streptococcus chenjunshii |
| MODERN | bin_59_mod | Streptococcus gallolyticus | 1.9 | 53.0 | 10.0 | 37.2 | 1490 | 1128409 | Bacteria | Firmicutes | Bacilli | Lactobacillales | Streptococcaceae | Streptococcus | Streptococcus gallolyticus |
| ANCIENT | bin_56_anc | Streptococcus sp. | 34.7 | 62.7 | 5.5 | 37.5 | 1774 | 1207602 | Bacteria | Firmicutes | Bacilli | Lactobacillales | Streptococcaceae | Streptococcus | |
| ANCIENT | bin_40_anc | Streptomyces sp. | 10.2 | 90.1 | 4.7 | 70.2 | 13783 | 5890215 | Bacteria | Actinobacteriota | Actinomycetia | Streptomycetales | Streptomycetaceae | Streptomyces | |
| ANCIENT | bin_C_dup | Succiniclasticum sp. | 86.9 | 94.6 | 3.9 | 46.9 | 7758 | 1694059 | Bacteria | Firmicutes_C | Negativicutes | Acidaminococcales | Acidaminococcaceae | Succiniclasticum | |
| MODERN | bin_C_dup | Succiniclasticum sp. | 86.9 | 95.6 | 2.0 | 46.7 | 29959 | 1989528 | Bacteria | Firmicutes_C | Negativicutes | Acidaminococcales | Acidaminococcaceae | Succiniclasticum | |
| ANCIENT | bin_43_anc | SZUA-320 sp. | 3.2 | 67.0 | 2.3 | 68 | 11686 | 3476177 | Bacteria | Gemmatimonadota | Gemmatimonadetes | Gemmatimonadales | GWC2-71-9 | SZUA-320 | |
| MODERN | bin_39_mod | Treponema_D sp. | 2.3 | 84.6 | 0.7 | 57.6 | 5425 | 2172720 | Bacteria | Spirochaetota | Spirochaetia | Treponematales | Treponemataceae | Treponema_D | |
| MODERN | bin_52_mod | Treponema_D sp | 2.1 | 72.7 | 0.7 | 48.5 | 3142 | 1660700 | Bacteria | Spirochaetota | Spirochaetia | Treponematales | Treponemataceae | Treponema_D | |
| MODERN | bin_40_mod | UBA1066 sp. | 1.9 | 53.8 | 3.1 | 58.6 | 1825 | 1339081 | Bacteria | Firmicutes_A | Clostridia | Lachnospirales | Lachnospiraceae | UBA1066 | |
| MODERN | bin_53_mod | UBA1367 sp. | 3.0 | 75.8 | 1.5 | 62.7 | 4744 | 1769629 | Bacteria | Actinobacteriota | Coriobacteriia | Coriobacteriales | Atopobiaceae | UBA1367 | |
| ANCIENT | bin_25_ | UBA1367 sp | 17.9 | 56.6 | 4.1 | 66.4 | 2176 | 1664139 | Bacteria | Actinobacteri | Coriobacteriia | Coriobacterial | Atopobiaceae | UBA1367 | |

| Co-Assembly batch | BIN | OTU | Sum abundance | Completion [%] | Contamination [%] | GC [%] | N50 | Total length [bp] | domain | phylum | class | order | family | genus | species |
|---|---|---|---|---|---|---|---|---|---|---|---|---|---|---|---|
| | anc | | | | | | | | | ota | | es | | | |
| MODERN | bin_19_mod | UBA2868 sp905235745 | 2.3 | 82.5 | 3.5 | 48.2 | 5319 | 1852118 | Bacteria | Firmicutes_A | Clostridia | Lachnospirales | Lachnospiraceae | UBA2868 | UBA2868 sp905235745 |
| ANCIENT | bin_53_anc | UBA3738 sp. | 26.7 | 84.2 | 5.0 | 45.1 | 6916 | 1475007 | Bacteria | Firmicutes_A | Clostridia | Peptostreptococcales | Anaerovoracaceae | UBA3738 | |
| ANCIENT | bin_13_anc | UBA4312 sp. | 46.8 | 75.0 | 5.7 | 43.4 | 5127 | 1714447 | Bacteria | Firmicutes | Bacilli | Erysipelotrichales | Erysipelotrichaceae | UBA4312 | |
| MODERN | bin_13_mod | UBA4312 sp002394635 | 1.2 | 56.6 | 14.3 | 41.5 | 1614 | 1856396 | Bacteria | Firmicutes | Bacilli | Erysipelotrichales | Erysipelotrichaceae | UBA4312 | UBA4312 sp002394635 |
| ANCIENT | bin_17_anc | UBA7541 | 5.3 | 94.0 | 3.6 | 62 | 29026 | 4496507 | Bacteria | Acidobacteriota | Acidobacteriae | Acidoferrales | UBA7541 | | |
| ANCIENT | bin_54_anc | UBA7541.1 | 2.1 | 60.5 | 2.1 | 63.9 | 3386 | 2247358 | Bacteria | Acidobacteriota | Acidobacteriae | Acidoferrales | UBA7541 | | |
| ANCIENT | bin_30_anc | UBA9715 sp. | 16.9 | 74.3 | 7.3 | 55.8 | 3258 | 2477898 | Bacteria | Actinobacteriota | Coriobacteriia | Coriobacteriales | Eggerthellaceae | UBA9715 | |
| ANCIENT | bin_7_anc | UBA9715 sp | 51.1 | 72.8 | 2.5 | 60 | 6621 | 2432159 | Bacteria | Actinobacteriota | Coriobacteriia | Coriobacteriales | Eggerthellaceae | UBA9715 | |
| MODERN | bin_60_mod | W3P20-009 sp905235955 | 3.6 | 91.2 | 1.0 | 42.5 | 6419 | 2459179 | Bacteria | Bacteroidota | Bacteroidia | Bacteroidales | W3P20-009 | W3P20-009 | W3P20-009 sp905235955 |
| ANCIENT | bin_1_anc | WHSY01 sp. | 75.3 | 97.8 | 6.0 | 70 | 14788 | 4179738 | Bacteria | Actinobacteriota | Actinomycetia | Mycobacteriales | Pseudonocardiaceae | WHSY01 | |
| ANCIENT | bin_5_anc | Yaniella sp. | 8.9 | 95.2 | 1.2 | 56.5 | 179919 | 3329375 | Bacteria | Actinobacteriota | Actinomycetia | Actinomycetales | Micrococcaceae | Yaniella | |

Supplementary Table 2. Significant results in the comparison of differential abundance between modern and ancient samples.

Results are shown for associations with false-discovery-rate (FDR) < 0.05. MAGs with a positive coefficient (> 0) had higher abundance in the modern samples, while those with a negative coefficient (< 0) were more abundant in ancient samples. Blanks, historical samples and ancient samples U027, U035 and LB003 were excluded from the analysis. Values are rounded to three decimal places.

| Bin | OTU | coefficient | standard error | p-value | FDR |
|---|---|---|---|---|---|
| bin_E_dup | Propionibacterium ruminifibrarum | -2.002 | 0.364 | 0 | 0 |
| bin_D_dup | RUG099 sp. | -5.378 | 0.416 | 0 | 0 |
| bin_C_dup | Succiniclasticum sp. | -1.384 | 0.418 | 0.002 | 0.004 |
| bin_B_dup | Streptococcus chenjunshii | -4.263 | 0.43 | 0 | 0 |
| bin_A_dup | Basfia_A sp. | 1.965 | 0.725 | 0.01 | 0.016 |
| bin_8_mod | Prevotella sp..3 | 1.069 | 0.317 | 0.002 | 0.003 |
| bin_7_mod | JAGOWQ01 sp..1 | 3.613 | 0.337 | 0 | 0 |
| bin_7_anc | UBA9715 sp..1 | -3.754 | 0.358 | 0 | 0 |
| bin_61_mod | Actinomyces sp. A.2 | 4.188 | 0.599 | 0 | 0 |
| bin_6_anc | Amycolatopsis_C sp009377395 | -0.855 | 0.323 | 0.012 | 0.018 |
| bin_57_mod | Mannheimia sp..1 | 5.713 | 0.378 | 0 | 0 |
| bin_57_anc | Lactobacillaceae | -0.788 | 0.243 | 0.003 | 0.004 |
| bin_56_anc | Streptococcus sp. | -2.396 | 0.453 | 0 | 0 |
| bin_55_anc | Saccharopolyspora dendranthemae | -2.705 | 0.463 | 0 | 0 |
| bin_53_mod | UBA1367 sp. | 1.33 | 0.362 | 0.001 | 0.002 |
| bin_53_anc | UBA3738 sp. | -2.552 | 0.237 | 0 | 0 |
| bin_50_mod | Fusobacterium_C necrophorum | 0.548 | 0.213 | 0.015 | 0.02 |
| bin_5_mod | Pseudomonas_E paraversuta | 1.155 | 0.51 | 0.03 | 0.036 |
| bin_49_mod | Bibersteinia sp. | 1.197 | 0.478 | 0.017 | 0.023 |

| Bin | OTU | coefficient | standard error | p-value | FDR |
|---|---|---|---|---|---|
| bin_48_anc | Allosaccharopolyspora sp..4 | -0.917 | 0.352 | 0.013 | 0.019 |
| bin_47_anc | Hornefia sp. | -1.987 | 0.285 | 0 | 0 |
| bin_46_mod | Actinomyces sp. B | 2.471 | 0.446 | 0 | 0 |
| bin_45_mod | Bacteroides sp. | 1.703 | 0.57 | 0.005 | 0.008 |
| bin_45_anc | Bilifractor sp. | -1.981 | 0.168 | 0 | 0 |
| bin_43_mod | RUG721 sp. | 1.496 | 0.413 | 0.001 | 0.002 |
| bin_41_anc | Methanobrevibacter_A sp..1 | -0.927 | 0.382 | 0.021 | 0.027 |
| bin_40_anc | Streptomyces sp. | -1.496 | 0.446 | 0.002 | 0.003 |
| bin_4_mod | Neisseria sp. | 1.875 | 0.537 | 0.001 | 0.003 |
| bin_4_anc | Methanosphaera sp..1 | -2.875 | 0.277 | 0 | 0 |
| bin_38_anc | Methanobrevibacter sp. | -4.05 | 0.396 | 0 | 0 |
| bin_36_anc | RUG11795 sp. | -2.178 | 0.112 | 0 | 0 |
| bin_35_anc | Allosaccharopolyspora sp..3 | -0.737 | 0.297 | 0.018 | 0.024 |
| bin_34_mod | Peptidiphaga sp. | 2.413 | 0.393 | 0 | 0 |
| bin_33_anc | Methanobrevibacter_A sp. | -1.386 | 0.472 | 0.006 | 0.009 |
| bin_32_mod | Actinomyces sp. A | 0.836 | 0.347 | 0.022 | 0.027 |
| bin_31_anc | CAG-791 sp. | -2.174 | 0.195 | 0 | 0 |
| bin_30_anc | UBA9715 sp. | -1.352 | 0.349 | 0 | 0.001 |
| bin_3_mod | Mannheimia sp. | 0.741 | 0.304 | 0.02 | 0.026 |
| bin_3_anc | Allosaccharopolyspora sp..2 | -2.823 | 0.547 | 0 | 0 |
| bin_29_mod | Cryptobacteroides sp. | 1.961 | 0.499 | 0 | 0.001 |
| bin_29_anc | Haemophilus_B sp. | -0.963 | 0.419 | 0.028 | 0.034 |
| bin_28_anc | Methanosphaera sp. | -4.288 | 0.413 | 0 | 0 |

| Bin | OTU | coefficient | standard error | p-value | FDR |
|---|---|---|---|---|---|
| bin_27_mod | Prevotella sp017627765 | 0.723 | 0.278 | 0.014 | 0.019 |
| bin_26_mod | Porphyromonas_A sp. | 0.666 | 0.293 | 0.029 | 0.036 |
| bin_25_anc | UBA1367 sp..1 | -2.025 | 0.338 | 0 | 0 |
| bin_24_anc | SFGY01 sp. | -2.153 | 0.481 | 0 | 0 |
| bin_23_anc | Allosaccharopolyspora sp..1 | -1.429 | 0.395 | 0.001 | 0.002 |
| bin_22_anc | Eubacterium_I sp..1 | -2.134 | 0.31 | 0 | 0 |
| bin_18_mod | Eubacterium_I sp. | 1.752 | 0.439 | 0 | 0.001 |
| bin_17_mod | Prevotella sp. | 1.024 | 0.374 | 0.01 | 0.015 |
| bin_16_anc | RUG099 sp..1 | -1.455 | 0.387 | 0.001 | 0.001 |
| bin_15_mod | Fusobacterium sp..1 | 1.307 | 0.404 | 0.003 | 0.005 |
| bin_13_anc | UBA4312 sp. | -3.613 | 0.257 | 0 | 0 |
| bin_12_mod | Desulfovibrio desulfuricans_D | 2.043 | 0.309 | 0 | 0 |
| bin_11_mod | Brachymonas sp. | 2.434 | 0.407 | 0 | 0 |
| bin_1_anc | WHSY01 sp. | -2.218 | 0.56 | 0 | 0.001 |

Supplementary Table 3. Summary alignment statistics of identified plant and lichen taxa (Separate document)

Supplementary Table 4. Detailed radiocarbon dating results.

Dating was undertaken at the NTNU University Museum’s National Laboratory for Age Determination (Trondheim, Norway). See (Seiler et al. 2019) for measurement reference. Dates are calibrated using OxCal 4.4.2 using the calibration curve IntCal20.

| Sample Name | Run Name | Fraction | 14C content (pMC) | 14C Age (rounded) | d13C (from AMS system) | Calibrated Age Ranges | % C | mgC | C content % by weight | N Content % by weight | C:N ratio by weight | 14C Age (not rounded) |
|---|---|---|---|---|---|---|---|---|---|---|---|---|
| LB006 | TRa-20235 | collagen | 18.25 ± 0.09 | 13665 ± 40 | -21.6 ± 0.9 ‰ | 68.3% probability 14646BC (68.3%) 14466BC 95.4% probability 14736BC (95.4%) 14391BC | 45.7 | 1.37 | 45.5707 | 16.7952 | 2.71 | 13666 +42/-42 BP |
| LB014 | TRa-20236 | 2nd PT Reindeer tooth, Collagen | 18.30 ± 0.11 | 13640 ± 55 | -18.3 ± 2.2 ‰ | 68.3% probability 14617BC (68.3%) 14422BC 95.4% probability 14726BC (95.4%) 14341BC | 38.1 | 0.8 | 38.3016 | 13.719 | 2.79 | 13640 +54/-54 BP |
| LB015 | TRa-20237 | collagen | 18.46 ± 0.09 | 13575 ± 45 | -20.2 ± 0.5 ‰ | 68.3% probability 14515BC (68.3%) 14348BC 95.4% probability 14608BC (95.4%) 14283BC | 35.5 | 1.1 | 35.594 | 12.9235 | 2.75 | 13573 +44/-44 BP |
| LB017 | TRa-20238 | collagen | 20.46 ± 0.10 | 12745 ± 45 | -22.7 ± 0.8 ‰ | 68.3% probability 13325BC (68.3%) 13181BC 95.4% probability 13406BC (95.4%) 13084BC | 37.7 | 0.83 | 37.9327 | 13.5046 | 2.81 | 12747 +46/-46 BP |
| U002 | TRa-20239 | collagen | 20.43 ± 0.09 | 12760 ± 35 | -19.4 ± 0.3 ‰ | 68.3% probability 13330BC (68.3%) 13202BC 95.4% probability 13403BC (95.4%) 13131BC | 42.3 | 1.1 | 42.473 | 15.4223 | 2.75 | 12760 +37/-37 BP |
| U004 | TRa-20240 | collagen | 8.80 ± 0.07 | 19530 ± 70 | -21.3 ± 1.4 ‰ | 68.3% probability 21799BC (11.7%) 21746BC 21648BC (56.6%) 21423BC 95.4% probability 21831BC (95.4%) 21339BC | 40.7 | 1.18 | 40.8102 | 14.984 | 2.72 | 19527 +68/-67 BP |
| U005 | TRa-20241 | collagen | 8.89 ± 0.08 | 19440 ± 80 | -21.1 ± 0.3 ‰ | 68.3% probability 21775BC (34.9%) 21597BC 21473BC (33.4%) 21300BC 95.4% probability 21805BC (95.4%) 21211BC | 31.2 | 1.03 | 31.3589 | 11.3776 | 2.76 | 19444 +78/-78 BP |
| U012 | TRa-15755 | Bein., Collagen | 16.15 ± 0.09 | 14645 ± 50 BP | -21.5 ± 0.6 ‰ | 68.3% probability 16119BC (68.3%) 15915BC 95.4% probability 16221BC (95.4%) 15841BC | 1.29 | 6.0287084 58 | 16.3695 | 2.72 | 86.9 | 14645 ± 49 BP |
| U016 | TRa-20223 | Collagen | 16.25 ± 0.08 | 14595 ± 45 | -18.9 ± 0.4 ‰ | 68.3% probability 16042BC (68.3%) 15841BC 95.4% probability 16169BC (95.4%) 15630BC | 40.4 | 1.09 | 40.5044 | 14.9079 | 2.72 | 14595 +44/-44 BP |

| Sampl e Name | Run Name | Fraction | 14C content (pMC) | 14C Age (rounded) | d13C (from AMS system) | Calibrated Age Ranges | % C | mgC | C content % by weight | N Content % by weight | C:N ratio by weight | 14C Age (not rounded) |
|---|---|---|---|---|---|---|---|---|---|---|---|---|
| U018 | TRa-20242 | collagen | 7.79 ± 0.06 | 20510 ± 70 | -18.7 ± 0.4 ‰ | 68.3% probability<br>22925BC (68.3%) 22621BC<br>95.4% probability<br>23009BC (95.4%) 22390BC | 46 | 1.38 | 46.061 | 17.0548 | 2.7 | 20507 +69/-68 BP |
| U023 | TRa-20243 | collagen | 21.52 ± 0.10 | 12340 ± 40 | -17.8 ± 2.3 ‰ | 68.3% probability<br>12842BC (15.2%) 12776BC<br>12473BC ( 7.1%) 12431BC<br>12416BC (46.0%) 12232BC<br>95.4% probability<br>12874BC (21.8%) 12745BC<br>12592BC (73.7%) 12178BC | 43.1 | 1.25 | 43.1037 | 15.7776 | 2.73 | 12342 +40/-40 BP |
| U027 | TRa-20244 | collagen | 9.17 ± 0.26 | 19200 +300/-200 BP | -17.2 ± 0.9 ‰ | 68.3% probability<br>21762BC (12.4%) 21626BC<br>21446BC (55.8%) 20973BC<br>95.4% probability<br>21822BC (95.4%) 20649BC | 24.5 | 0.27 | 24.7828 | 8.64427 | 2.87 | 19195 +250/-242 BP |
| U031 | TRa-20245 | collagen | 8.55 ± 0.14 | 19760 ± 140 | -17.7 ± 1.0 ‰ | 68.3% probability<br>21986BC (49.5%) 21773BC<br>21601BC (18.7%) 21471BC<br>95.4% probability<br>22217BC (67.5%) 21723BC<br>21678BC (28.0%) 21398BC | 36.7 | 0.55 | 36.5784 | 12.8638 | 2.84 | 19757 +139/-137 BP |
| U035 | TRa-20246 | collagen | 17.54 ± 0.08 | 13985 ± 35 | -17.6 ± 2.7 ‰ | 68.3% probability<br>15116BC (68.3%) 15026BC<br>95.4% probability<br>15277BC ( 0.5%) 15260BC<br>15171BC (95.0%) 14912BC | 41.1 | 1.44 | 41.2672 | 15.4282 | 2.67 | 13985 +37/-37 BP |
| U041 | TRa-20247 | collagen | 9.08 ± 0.08 | 19270 ± 70 | -18.6 ± 2.6 ‰ | 68.3% probability<br>21333BC (68.3%) 21084BC<br>95.4% probability<br>21755BC (10.8%) 21637BC<br>21430BC (84.7%) 21034BC | 38.7 | 1.2 | 38.6146 | 14.1958 | 2.72 | 19274 +75/-74 BP |
| U048 | TRa-20248 | collagen | 8.82 ± 0.11 | 19510 +110/-100 BP | -19.0 ± 1.9 ‰ | 68.3% probability<br>21797BC (13.8%) 21721BC<br>21676BC (54.4%) 21393BC<br>95.4% probability<br>21837BC (95.4%) 21234BC | 29.2 | 0.7 | 29.0486 | 10.4898 | 2.77 | 19505 +105/-104 BP |
| U059 | TRa-20249 | collagen | 8.83 ± 0.07 | 19490 ± 70 | -17.8 ± 2.5 ‰ | 68.3% probability<br>21787BC (13.1%) 21724BC<br>21673BC (27.9%) 21541BC<br>21522BC (27.3%) 21396BC<br>95.4% probability<br>21819BC (95.4%) 21283BC | 37.5 | 1.2 | 37.437 | 13.7553 | 2.72 | 19492 +72/-71 BP |

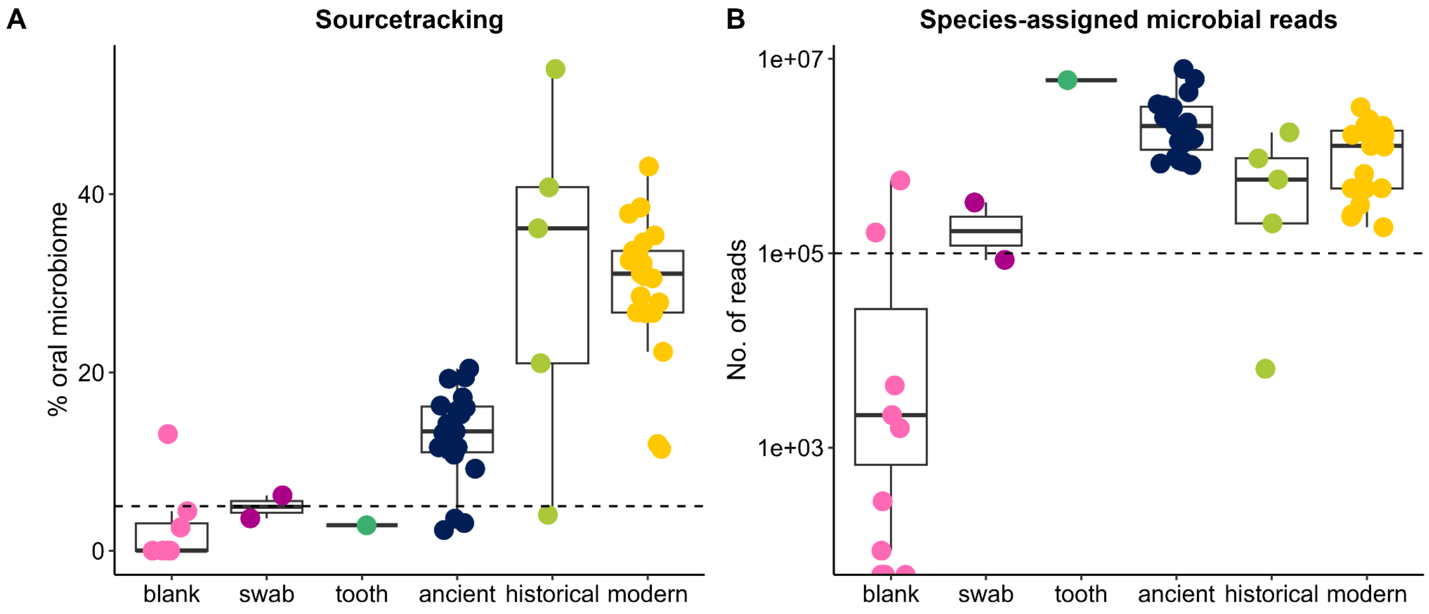

Supplementary Figure 1. Filtering of samples with low endogenous oral microbiome content.

**A.** Proportion of oral microbiome, based on decOM sourcetracking estimates (combined modern and ancient microbiome estimates). Samples below the dashed line (5%) were removed from further analysis, as not having enough endogenous oral microbiome content. **B.** Number of microbial reads assigned to the species-level by *KrakenUniq* in each sample. Samples below the dashed line (100,000 reads) were removed from further analysis, due to low read counts. In both plots, samples are coloured by sample type.

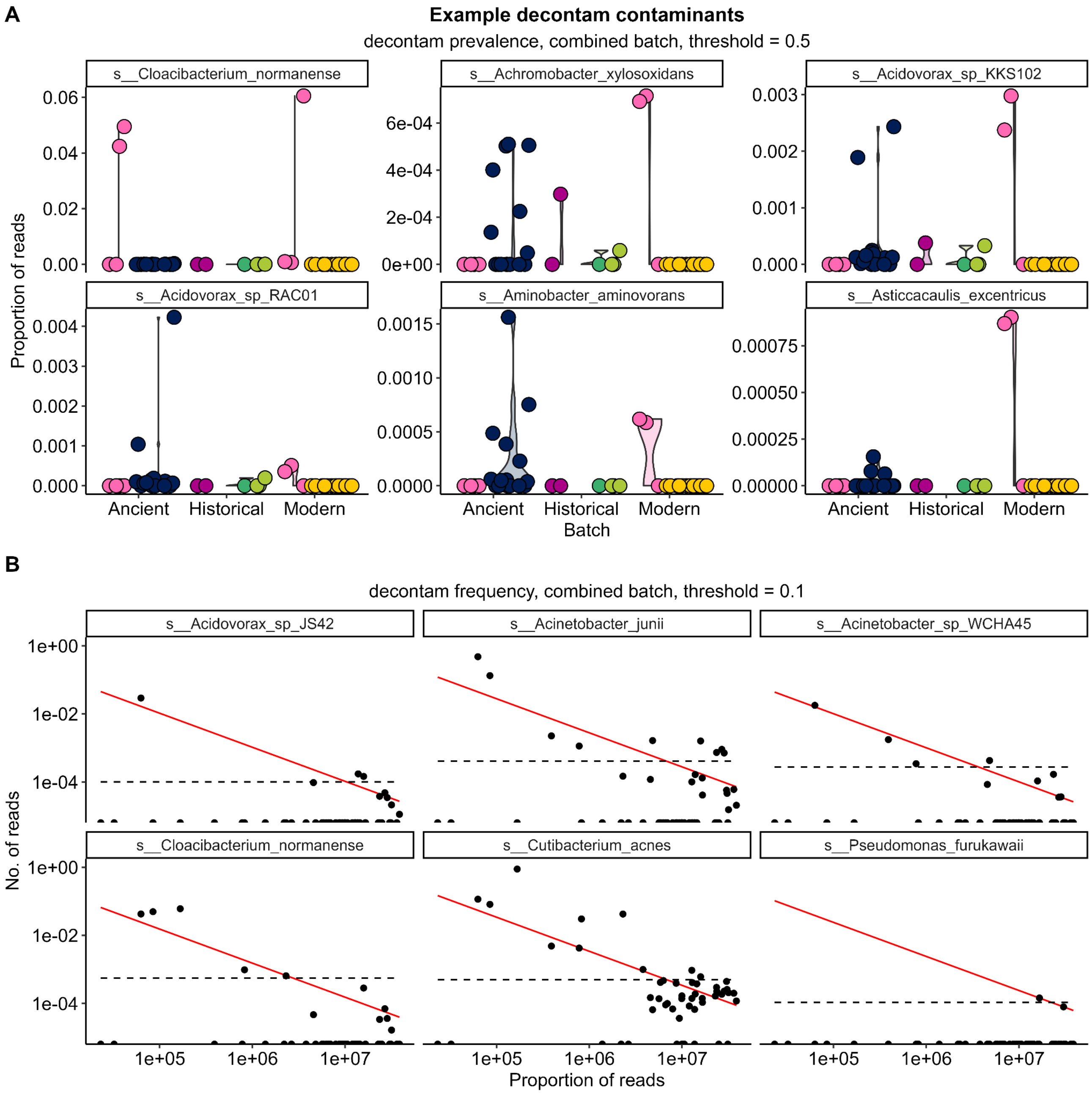

Supplementary Figure 2. Putative contaminant taxa removed during filtering with *decontam*.

**A.** Relative abundance (proportion of reads) of putative contaminants identified by the *prevalence* function, as having higher prevalence in control samples compared to dental calculus samples. Samples are grouped by extraction batch (ancient dataset, historical dataset or modern dataset) and coloured by sample type (bright pink: blanks, dark pink: museum swabs, aqua: museum tooth, dark blue: ancient dental calculus, green: historical dental calculus, yellow: modern dental calculus). **B.** Number of reads vs relative abundance (proportion of reads) of putative contaminants identified by the *frequency* function. Taxa where relative abundance is negatively correlated with sequencing depth (indicated by red lines) are likely contaminants. In both plots, only a subset of putative contaminants are shown.

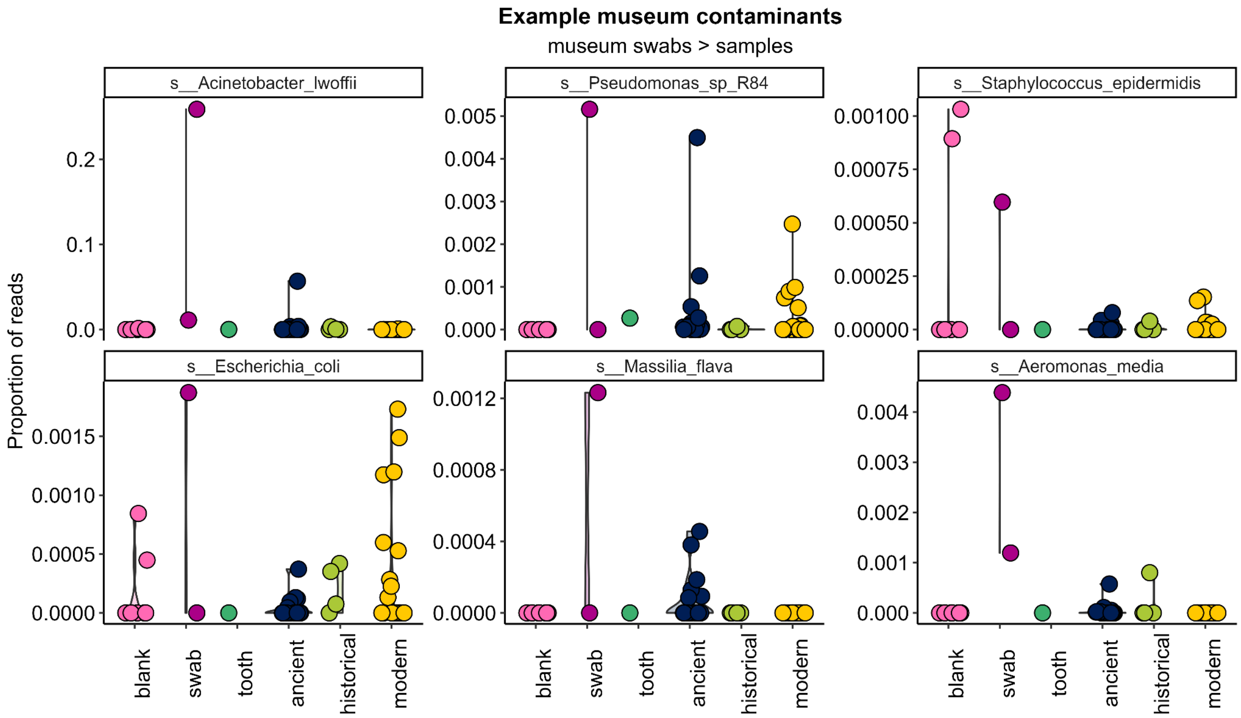

Supplementary Figure 3. Putative contaminant taxa removed as having higher relative abundance in museum controls than dental calculus.

Relative abundance of a subset of putative museum contaminants are shown. Samples are grouped by sample type.

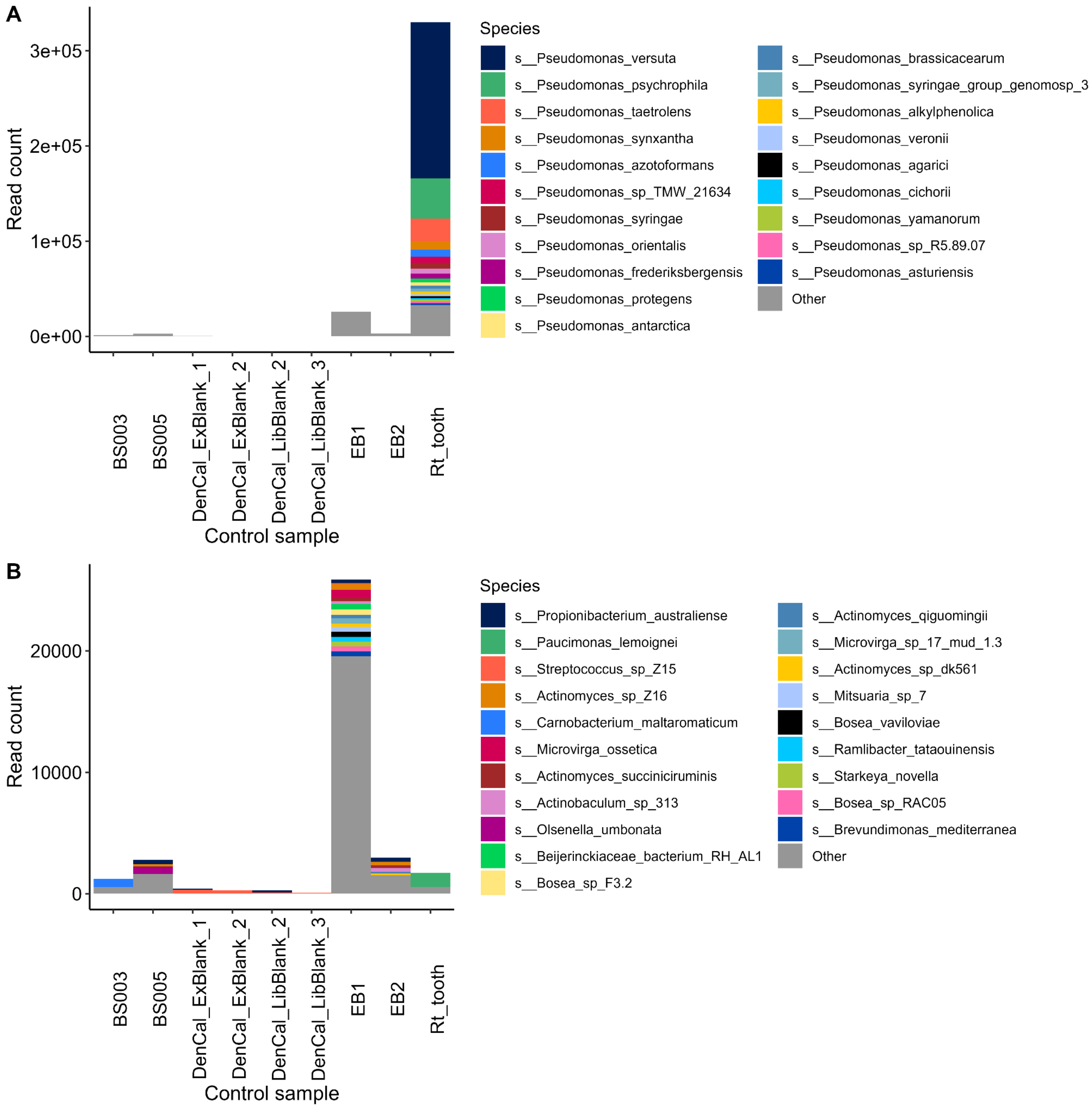

Supplementary Figure 4. Taxa remaining in control samples after other filtering steps.

**A.** After filtering taxa identified as contaminants using *decontam* and the museum swabs, various *Pseudomonas* species remain at high read counts, particularly in the museum tooth sample. **B.** After removing all *Pseudomonas* species, all remaining taxa in the control samples are at low read numbers.

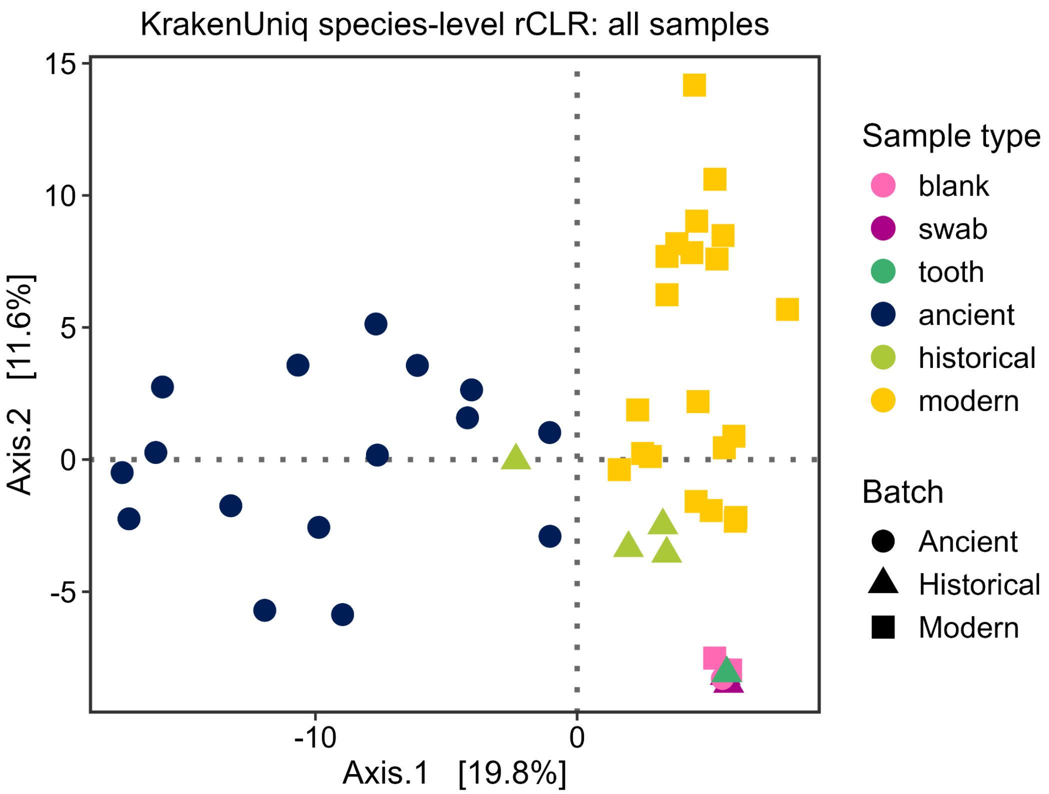

Supplementary Figure 5. Community composition ordination of control samples and dental calculus samples after contamination filtering of *KrakenUniq* classifications.

Samples were projected onto a PCoA using Euclidean distances of microbiome dissimilarity, based on the robust CLR transformation of *KrakenUniq* species-level assignments. Samples are coloured by sample type, while shapes indicate the extraction batch (dataset). All control samples are in an overlapping cluster of the PCoA, away from the dental calculus samples.

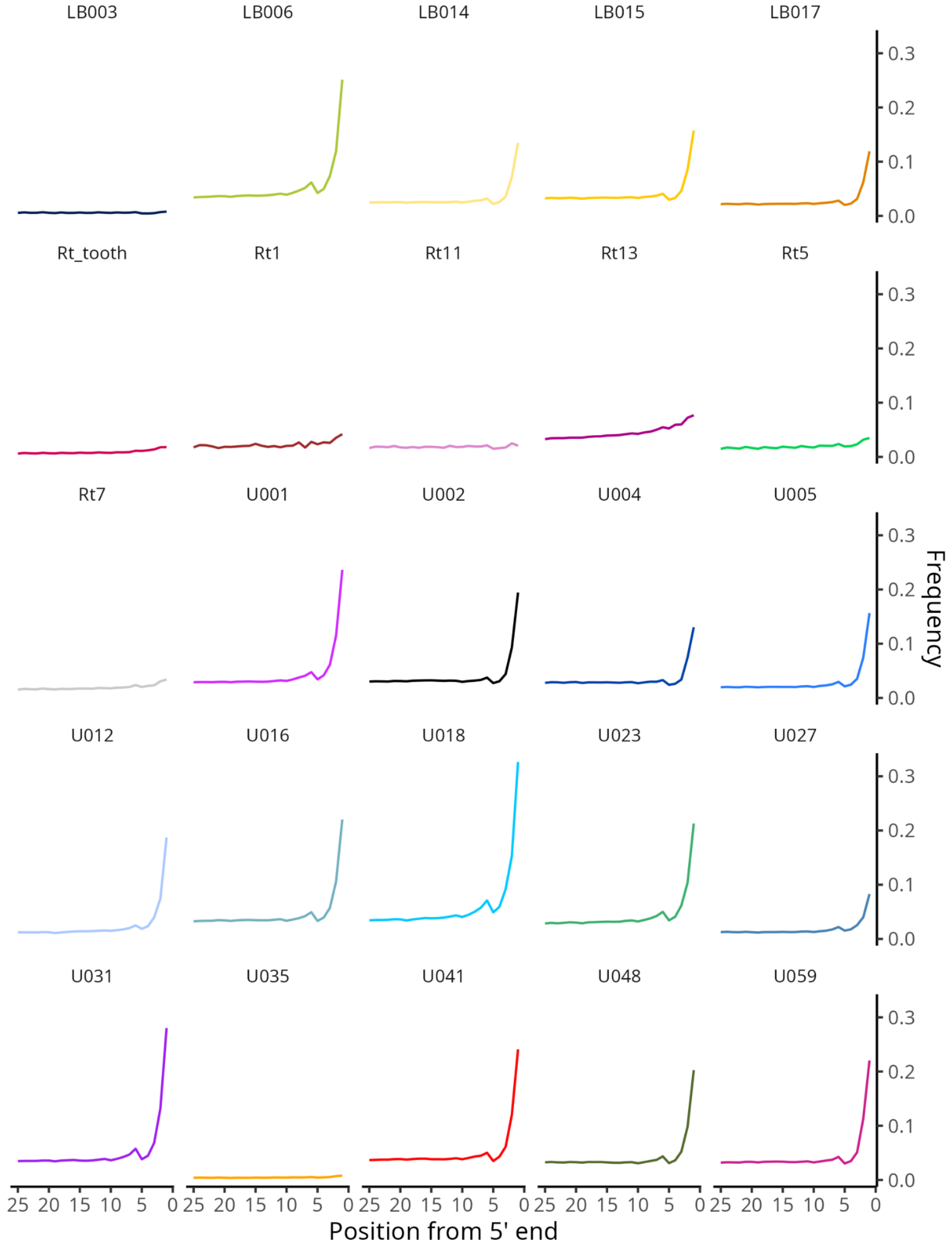

Supplementary Figure 6. Ancient DNA damage patterns.

Frequency of cytosine to thymine transitions on the 5'-end of microbial reads in ancient and historical samples, based on assembled microbial contigs, as calculated with mapDamage2 (Jónsson et al. 2013).

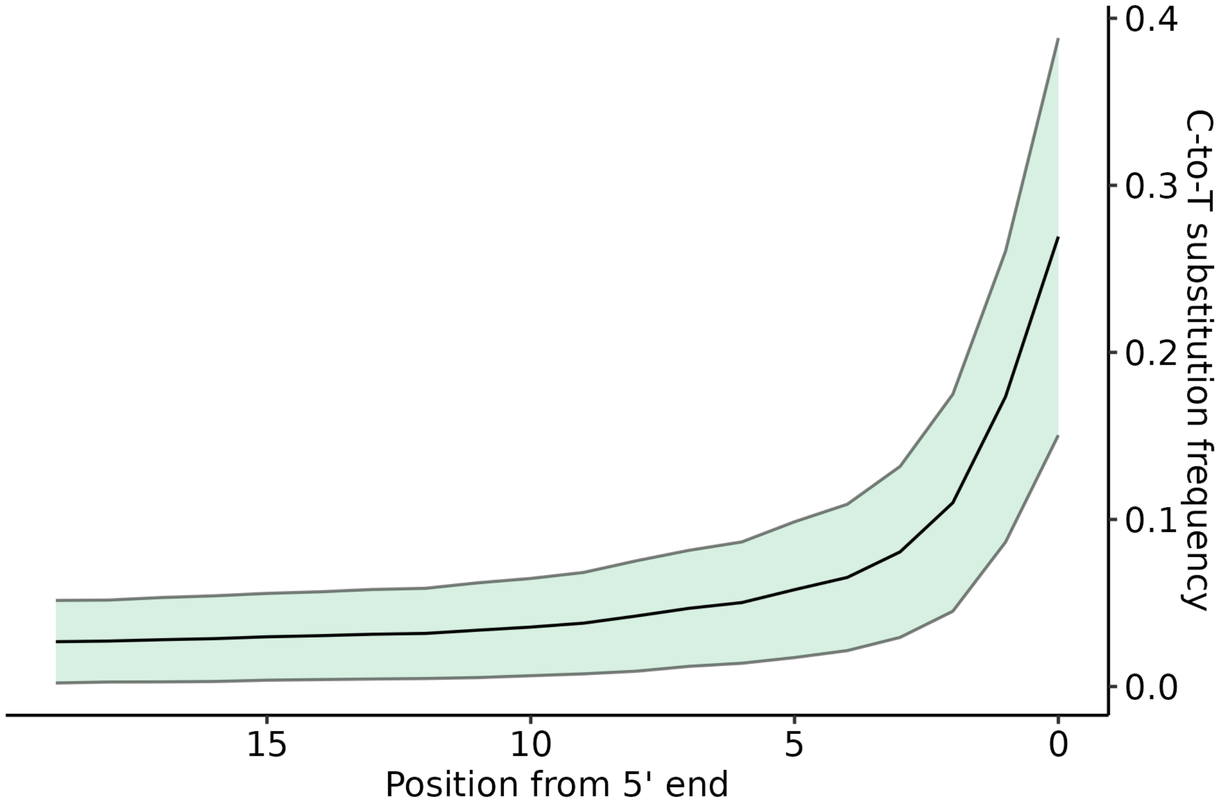

Supplementary Figure 7. Ancient DNA damage patterns in the ancient co-assembly.

Frequency of cytosine to thymine transitions as estimated with PyDamage (Borry et al. 2021). Mean substitution frequency at each position from 5’ end (black line). Gray lines bordering colored ribbon represent mean plus (upper) or minus (lower) one standard deviation.

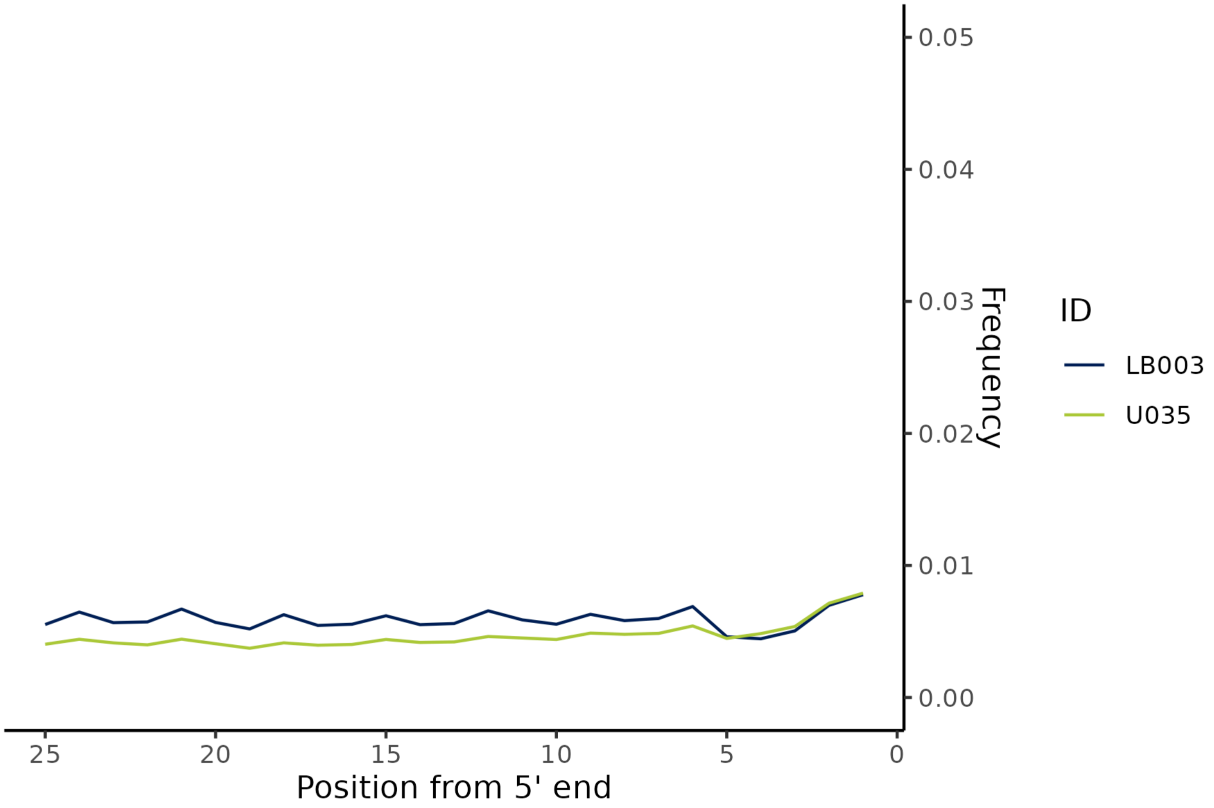

Supplementary Figure 8. Ancient DNA damage patterns for two ancient samples with unusually low levels of ancient DNA damage.

Frequency of cytosine to thymine transitions on the 5’-end of microbial reads have been calculated with mapDamage2 (Jónsson et al. 2013).

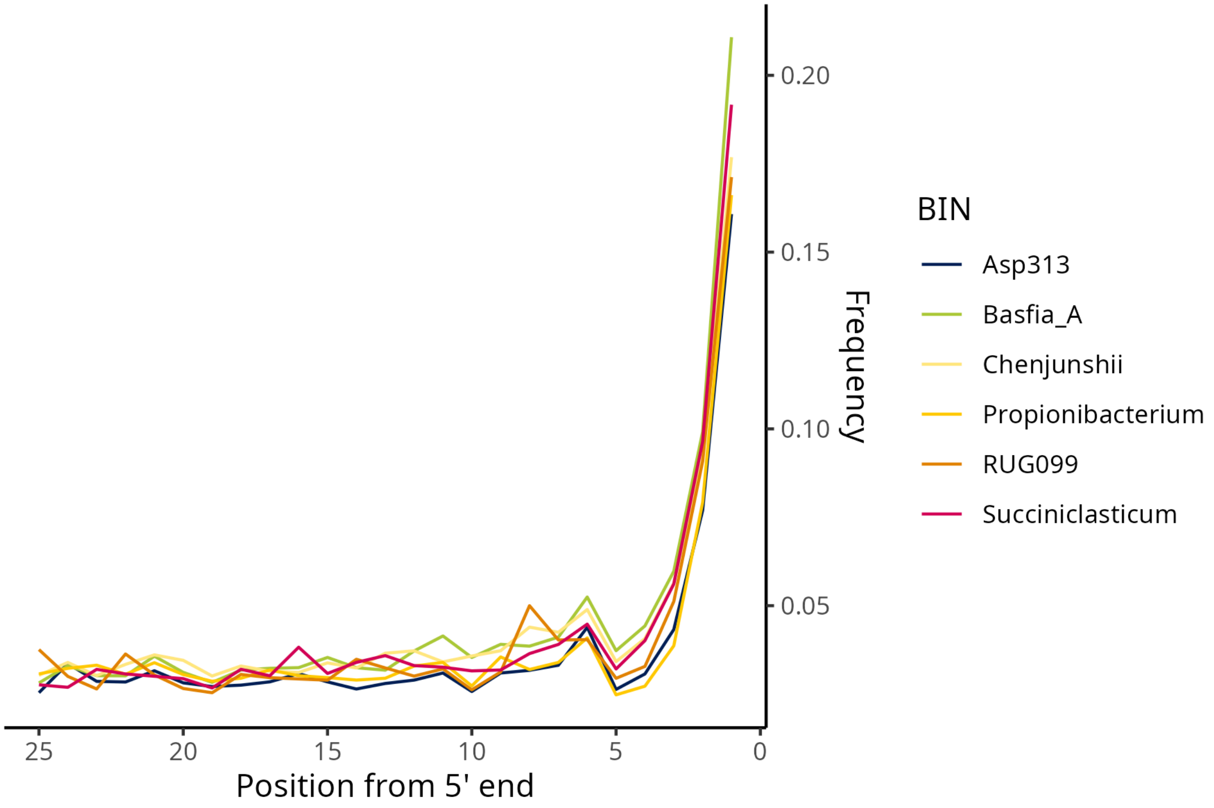

Supplementary Figure 9. Ancient DNA damage patterns.

Frequency of cytosine to thymine transitions on the 5’-end of microbial reads in in ancient samples for six bins identified in both ancient/historical and modern dental calculus samples, as calculated with mapDamage2 (Jónsson et al. 2013)

.

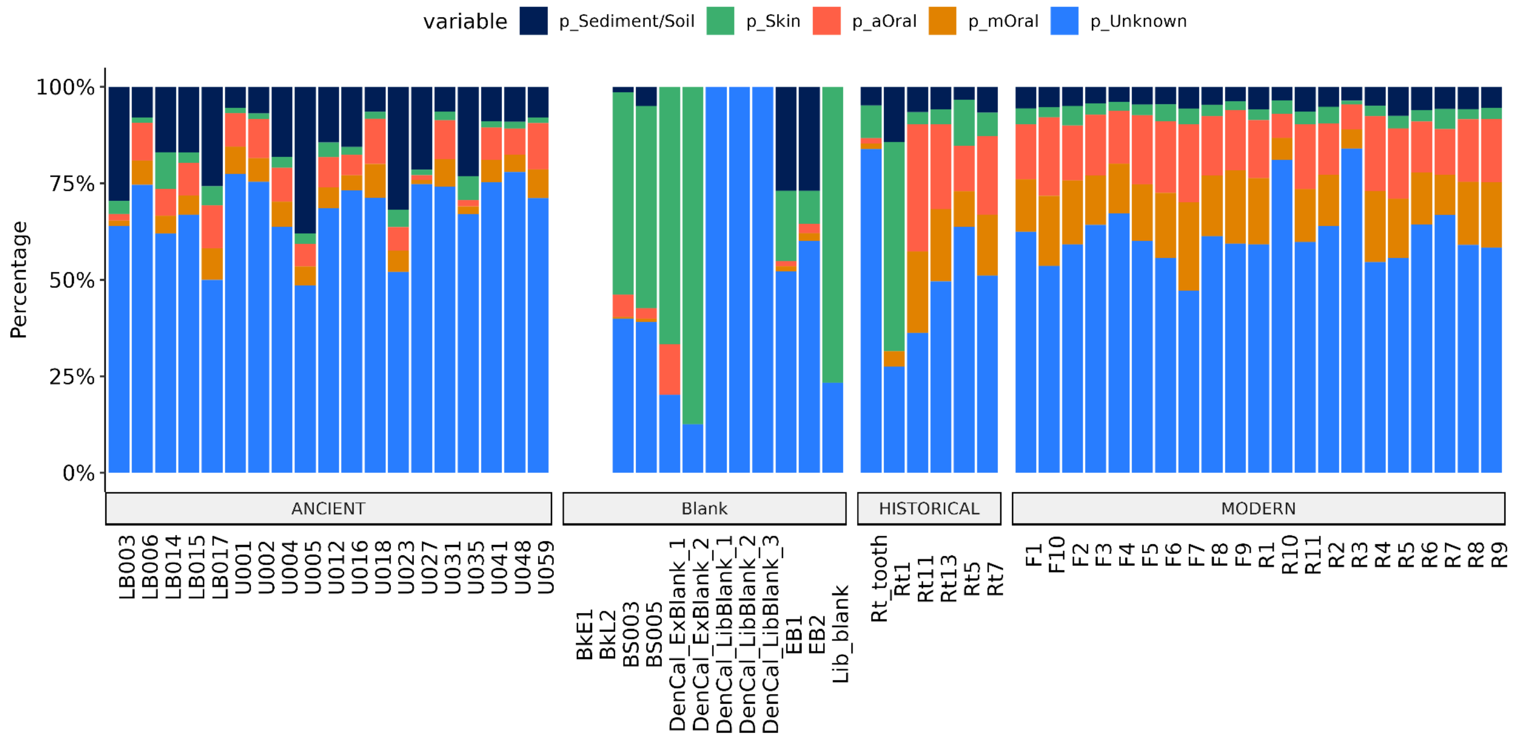

Supplementary Figure 10. Proportion of assigned putative DNA source environments per sample.

Samples are grouped by age. Classification was conducted with decOM (Duitama González et al. 2023) against the program's default database. aOral = ancient oral, mOral = modern oral

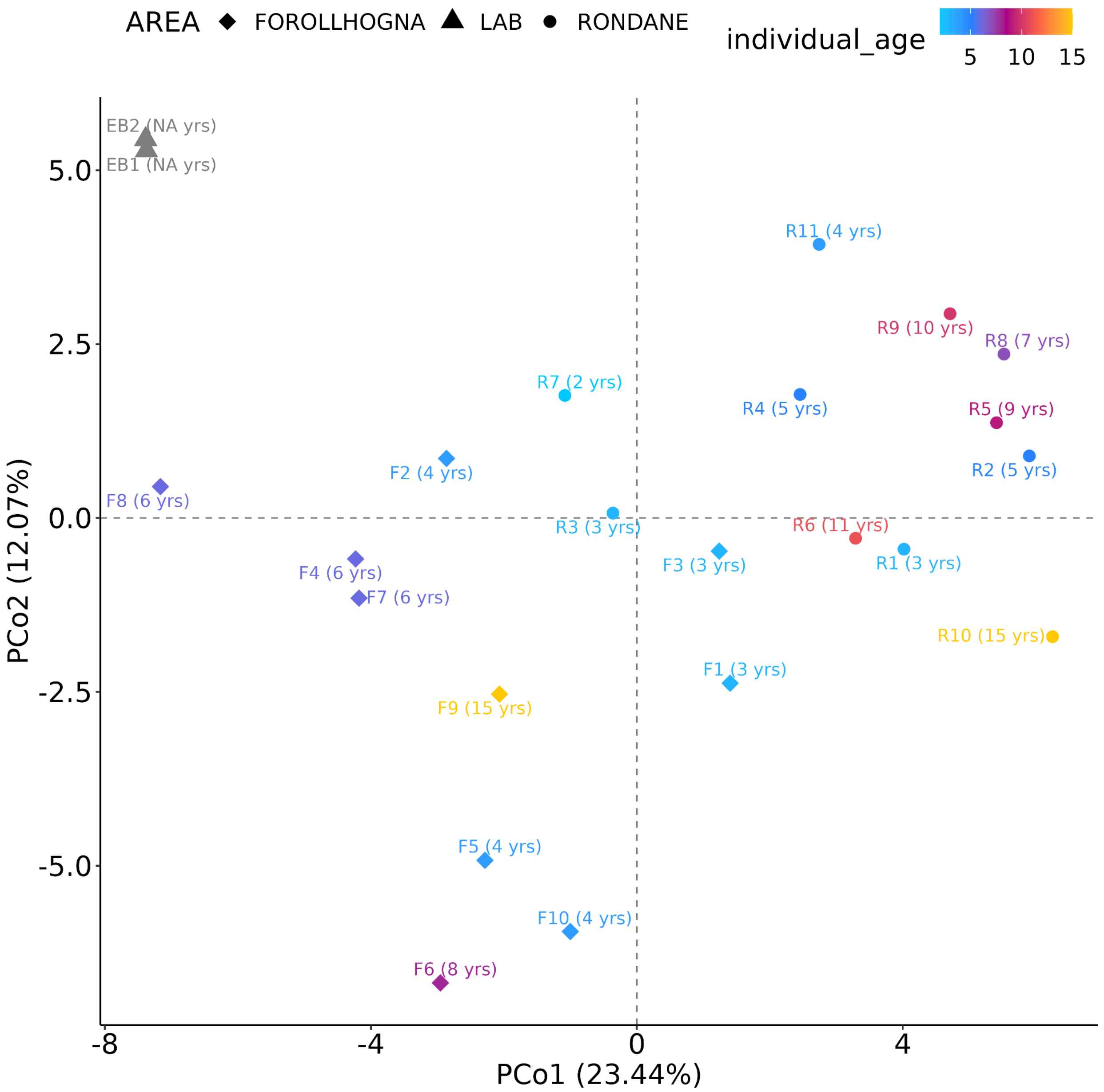

Supplementary Figure 11. PCoA plot of the relationship between taxon abundance (based on MAGs), individual age at death and geography of modern samples.

Ancient and historic samples were removed from the dataset for this analysis. Samples are colored by individual age at death on a scale from 2 to 15 years (see also Table 1). Laboratory blanks are colored in gray. Geographic origin of samples is coded by shape (diamonds = Forollhogna, Norway; circles = Rondane, Norway; triangles = laboratory blanks). Each data point is labeled with sample name followed by individual age at death in parentheses. Abundance is corrected for detection threshold (considering samples to be 0 abundance if under detection limit.) Abundance values were log-transformed, with a pseudo-count of 0.001 for 0 abundance values (i.e. -3 after log-transformation). Samples with abundance equal to 0 are omitted.

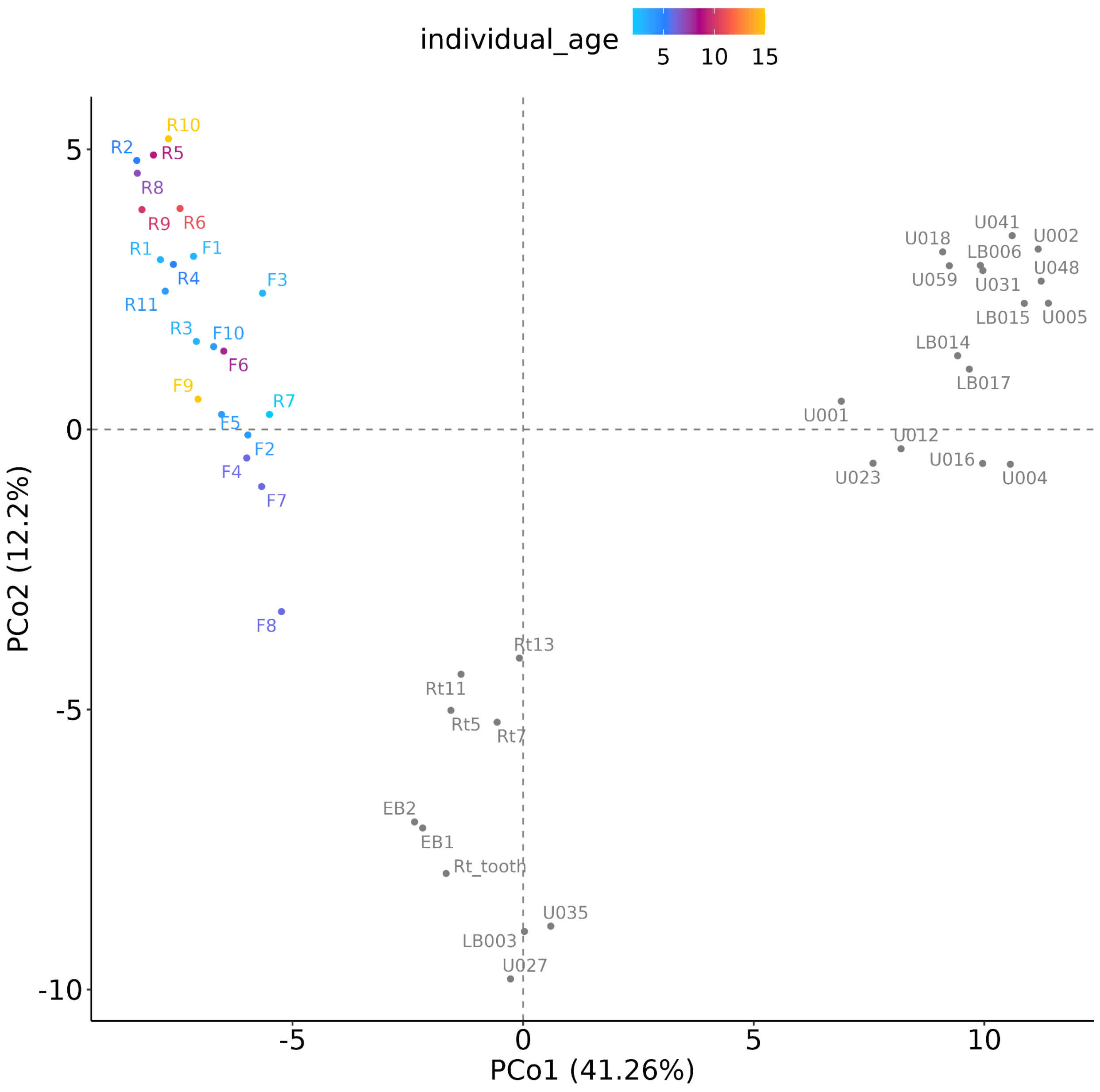

Supplementary Figure 12. PCoA plot of the relationship between taxon abundance (based on MAGs), samples and individual age at death.

Samples are colored by individual age at death (if known) on a scale from 2 to 15 years (see also Table 1). Samples with unknown age at death and laboratory blanks are colored in gray. Abundance is corrected for detection threshold (considering samples to be 0 abundance if under detection limit.) Abundance values were log-transformed, with a pseudo-count of 0.001 for 0 abundance values (i.e. -3 after log-transformation). Samples with abundance equal to 0 are omitted.

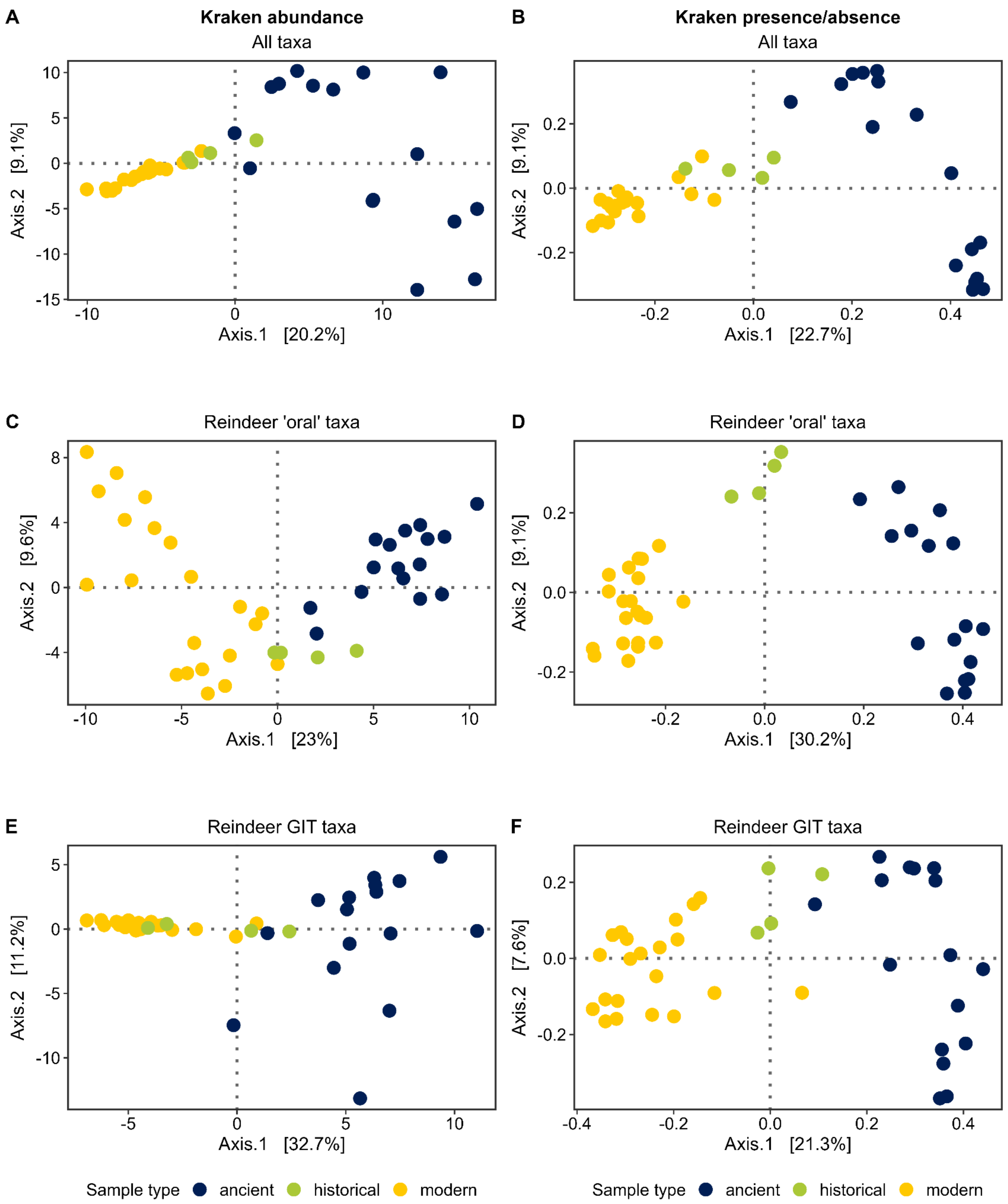

Supplementary Figure 13. Community composition ordinations using *KrakenUniq* classifications.

**A-B** Ordinations using all species-level taxa included in the final analysis. **C-D**. Ordinations using reindeer 'oral' species (species detected in 50% of the modern samples, and species- and genus-level taxa previously identified in the primate oral microbiome). **E-F**. Ordinations using reindeer GIT species (based on genus-level taxa previously identified in Norwegian reindeer rumen and GIT microbiomes). PCoAs were generated using abundance of reads (A, C, E) with the robust CLR transformation and Euclidean distances, and using presence/absence of taxa (B, D, F) with Jaccard distances.

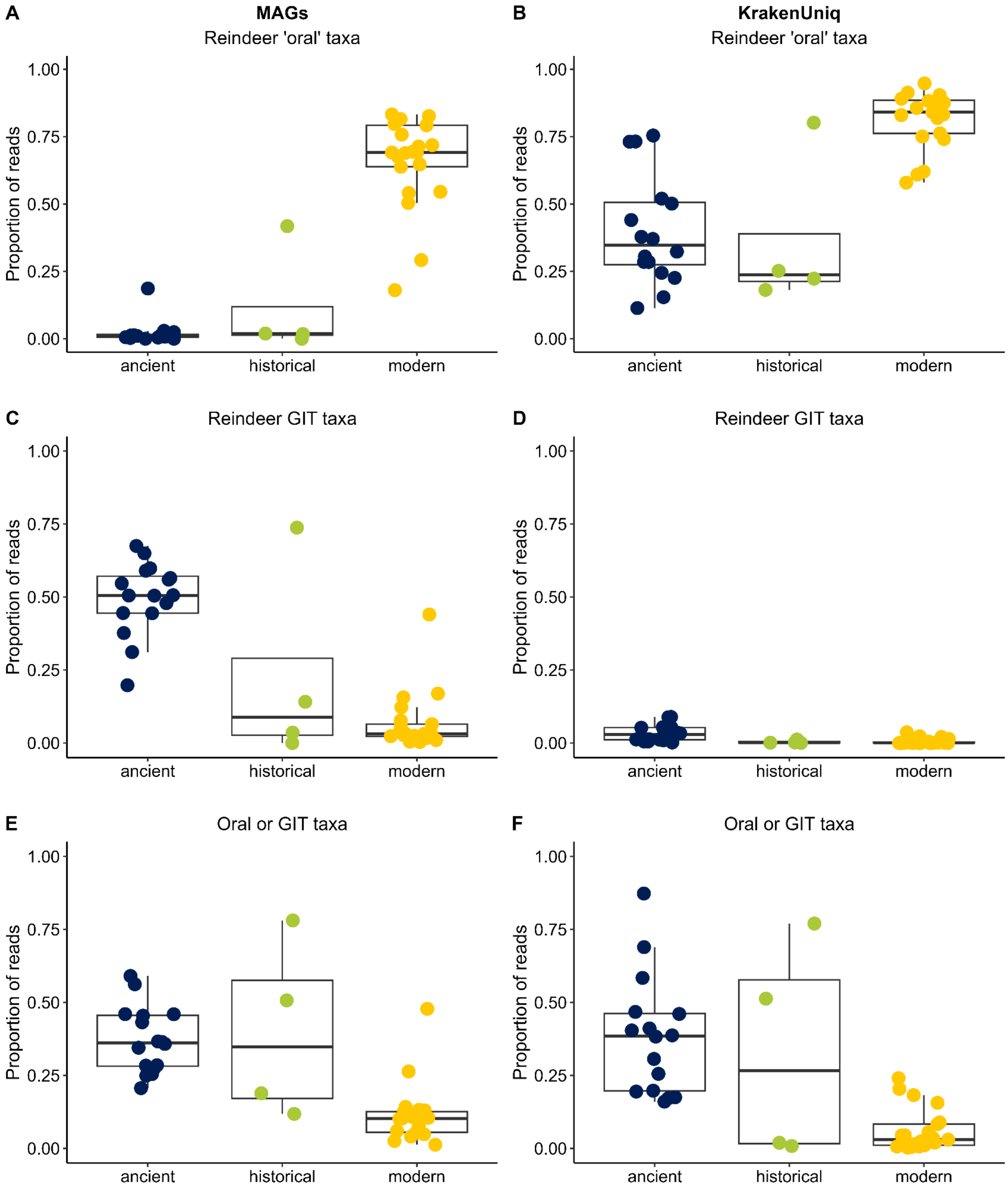

Supplementary Figure 14. Proportion of reindeer oral or GIT taxa assigned by MAG assembly (A, C, E) or *KrakenUniq* (B, D, F).

**A-B.** Reindeer ‘oral’ species (species detected in 50% of the modern samples, and species- and genus-level taxa previously identified in the primate oral microbiome). **C-D.** Reindeer GIT species (based on genus-level taxa previously identified in Norwegian reindeer rumen and GIT microbiomes). E-F. Species that were identified as both reindeer oral and GIT species.

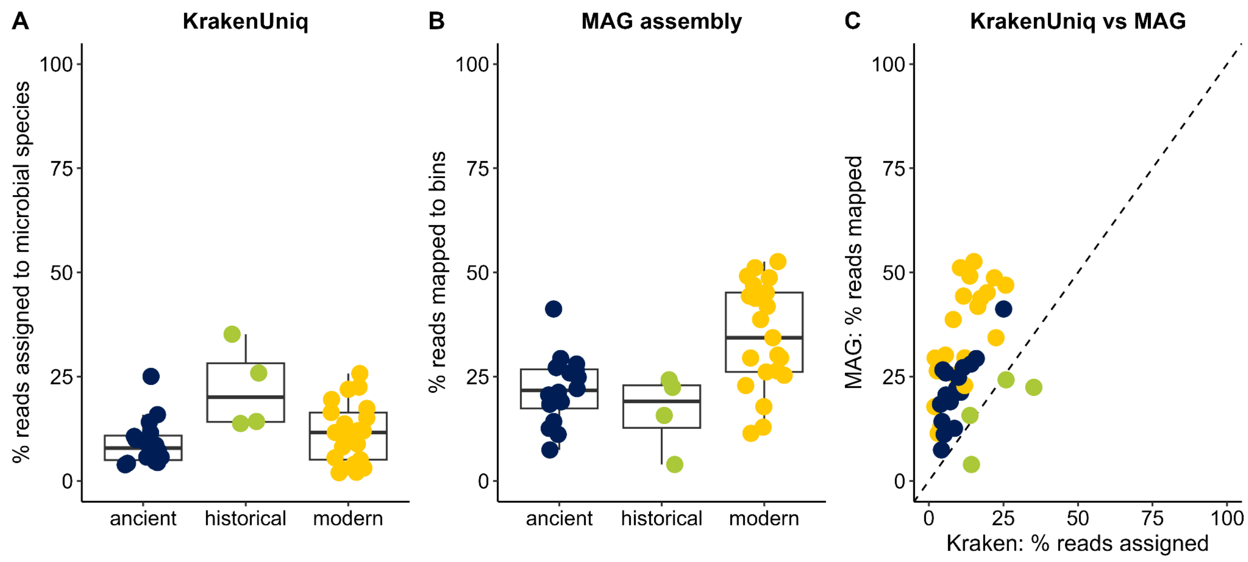

Supplementary Figure 15. Read recruitment comparison of *KrakenUniq* and MAG assembly.

**A.** Percent of reads assigned to species-level prokaryotic taxa by KrakenUniq (after all filtering steps). **B.** Percent of reads mapping to MAG bins. **C.** Comparison of percent of reads assigned by KrakenUniq and mapping to MAG bins (after all filtering steps). Samples are coloured by sample type, as indicated by the x-axis in A and B. The dashed line represents a perfect correlation, where both methods result in the same read recruitment. Most samples fall above the line, indicating higher read recruitment to MAG bins.

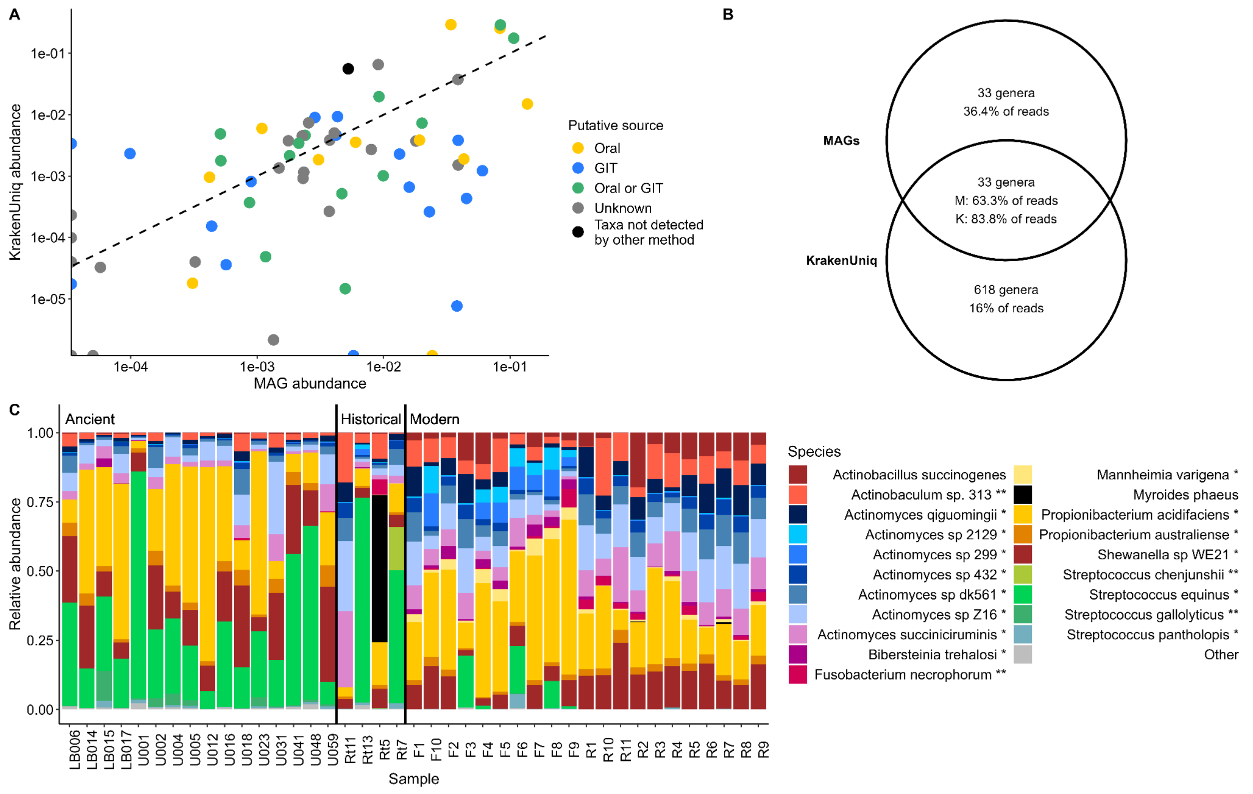

Supplementary Figure 16. Relative abundance of microbial taxa using *KrakenUniq*, compared to MAG assemblies.

**A.** Comparison of relative read abundances calculated per taxon across all dental calculus samples from KrakenUniq assignments, compared to reads mapped to MAG bins. Each taxon is summarised to the lowest taxonomic level classified by both approaches. E.g. *Streptococcus chenjunshii* and *Streptococcus gallolyticus* were detected by both approaches and are each presented as a separate data point; however other *Streptococcus* species were detected by only one approach, and are thus summarised to a single datapoint as 'other *Streptococcus*'. The dashed line represents a perfect correlation in relative abundance between the two methods. Each taxon is coloured by their putative source (oral-associated, GIT-associated, associated with either, or unknown). The black point presents all taxa that were not detected by the other approach at a taxonomic level higher than class. **B.** Venn diagram of the number of genera unique to or detected by both approaches, and summed relative abundance of the taxa in each set for each approach. **C.** Relative abundance for each sample of the 20 most abundant taxa detected by KrakenUniq. The relative abundances of all other taxa have been summarised in 'Other'. In the legend, species are annotated with ** if they were also detected at the species-level by the MAG approach, and with * if the genus (but not the species) was detected by the MAG approach.

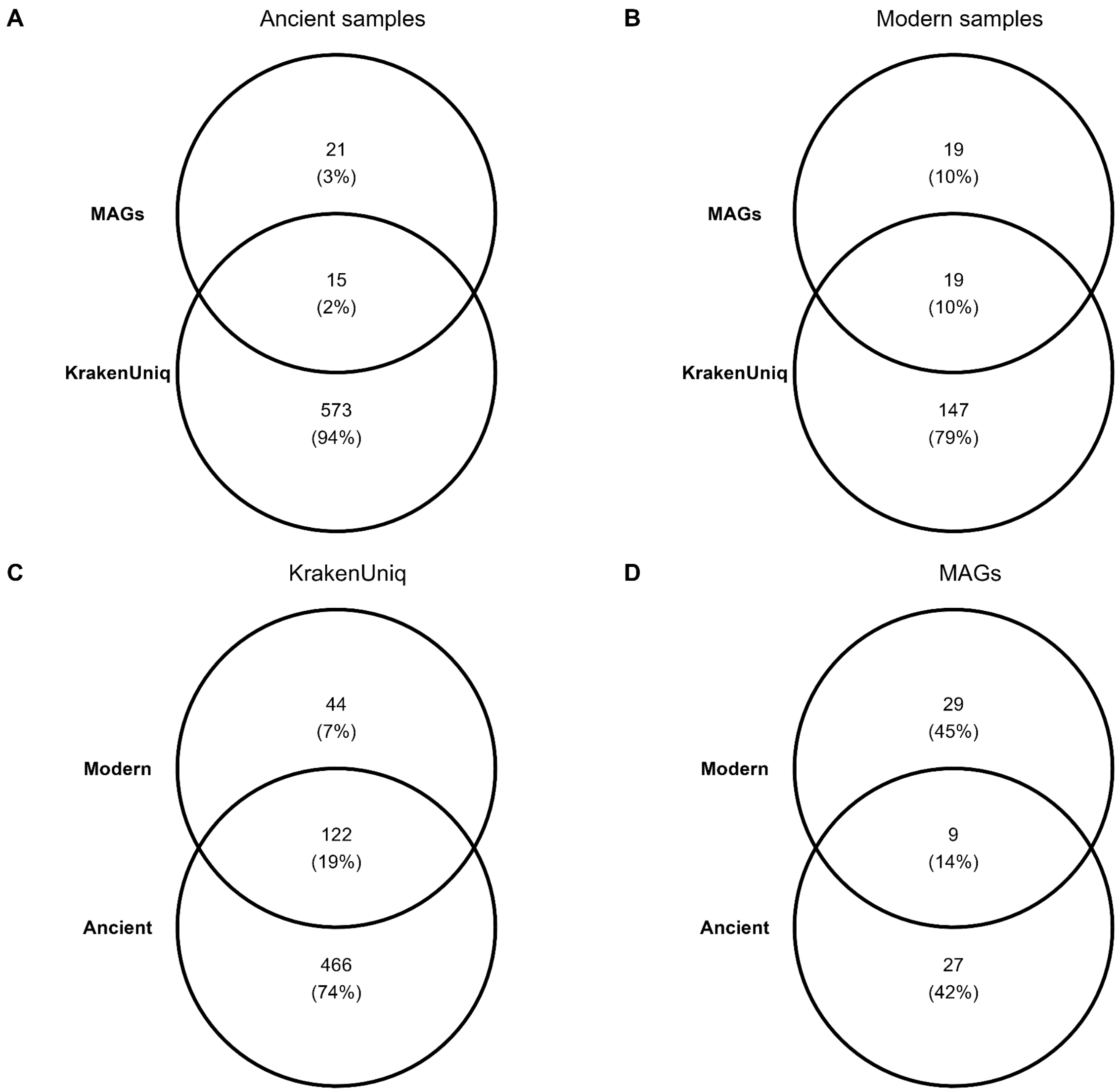

Supplementary Figure 17. Venn diagrams of KrakenUniq and MAG detection of genera in ancient and modern dental calculus samples.

**A.** Genera detected by KrakenUniq, the MAG approach or both approaches in ancient dental calculus samples. **B.** Genera detected by KrakenUniq, the MAG approach or both approaches in modern dental calculus samples. **C.** General detected by KrakenUniq and shared or unique to modern or ancient samples. **D**. General detected in the MAGs and shared or unique to modern or ancient samples. Historical samples are not shown in this figure. The number of genera in each region of each venn is shown, with the percent of all genera included in the venn in brackets.

References

Borry M, Hübner A, Rohrlach AB, Warinner C. 2021. PyDamage: automated ancient damage identification and estimation for contigs in ancient DNA de novo assembly. *PeerJ* [Internet] 9:e11845. Available from: http://dx.doi.org/10.7717/peerj.11845

Davis NM, Proctor DM, Holmes SP, Relman DA, Callahan BJ. 2018. Simple statistical identification and removal of contaminant sequences in marker-gene and metagenomics data. *Microbiome* [Internet] 6:226. Available from: http://dx.doi.org/10.1186/s40168-018-0605-2

Duitama González C, Vicedomini R, Lemane T, Rascovan N, Richard H, Chikhi R. 2023. decOM: similarity-based microbial source tracking of ancient oral samples using k-mer-based methods. *Microbiome* [Internet] 11:243. Available from: http://dx.doi.org/10.1186/s40168-023-01670-3

Jónsson H, Ginolhac A, Schubert M, Johnson PLF, Orlando L. 2013. mapDamage2.0: fast approximate Bayesian estimates of ancient DNA damage parameters. *Bioinformatics* [Internet] 29:1682–1684. Available from: http://dx.doi.org/10.1093/bioinformatics/btt193

Parks DH, Imelfort M, Skennerton CT, Hugenholtz P, Tyson GW. 2015. CheckM: assessing the quality of microbial genomes recovered from isolates, single cells, and metagenomes. *Genome Res.* [Internet] 25:1043–1055. Available from: http://dx.doi.org/10.1101/gr.186072.114

Seiler M, Grootes PM, Haarsaker J, Lélu S, Rzadeczka-Juga I, Stene S, Svarva H, Thun T, Værnes E, Nadeau M-J. 2019. Status report of the Trondheim radiocarbon laboratory. *Radiocarbon* [Internet] 61:1963–1972. Available from: http://dx.doi.org/10.1017/rdc.2019.115
